# isGWAS: ultra-high-throughput, scalable and equitable inference of genetic associations with disease

**DOI:** 10.1101/2023.07.21.550074

**Authors:** Christopher N Foley, Zhana Kuncheva, Riccardo E. Marioni, Heiko Runz, Benjamin B Sun

**Affiliations:** Optima Partners, Edinburgh, UK; Bayes Centre, University of Edinburgh, Edinburgh, UK; Centre for Genomic and Experimental Medicine, Institute of Genetics and Cancer, University of Edinburgh, Edinburgh, UK; Translational Sciences, Biogen Inc., Cambridge, MA, US

## Abstract

Genome-wide association studies (GWAS) have proven a powerful tool for human geneticists to generate biological insights or hypotheses for drug discovery. Nevertheless, a dependency on sensitive individual-level data together with ever-increasing cohort sample sizes, numbers of variants and phenotypes studied put a strain on existing algorithms, limiting the GWAS approach from maximising potential. Here we present in-silico GWAS (isGWAS), a uniquely scalable algorithm to infer regression parameters in case-control GWAS from cohort-level summary data. For any sample size, isGWAS computes a variant-disease association parameter in ∼1 millisecond, or ∼11m variants in UK-Biobank within ∼4 minutes (∼1500-fold faster than state-of-the-art). Extensive simulations and empirical tests demonstrate that isGWAS results are highly comparable to traditional regression-based approaches. We further introduce a heuristic re-sampling algorithm, leapfrog re-sampler (LRS), to extrapolate association results to semi-virtually enlarged cohorts. Owing to significant computational gains we anticipate a broad use of isGWAS and LRS which are customizable on a web interface.

## Main

Genome wide association studies (GWAS) have been immensely successful in unravelling the genetic contribution to human disease. Cost-effective genotyping and large biobank cohorts now make it possible to routinely conduct GWAS for tens of millions of variants in hundreds of thousands of individuals across thousands of phenotypes[1]. With the advent of population-scale whole genome sequencing and expansion of GWAS to research participants of non-European ancestries, these numbers can be expected to increase by another magnitude over the next few years[2], [3].

Current GWAS approaches that compute variant-disease associations in a regression framework, such as PLINK[4], fastGWA[5], BOLT-LMM[6], SAIGE[7] and REGENIE[8], require access to and input from ever increasing individual-level data (ILD). The efforts of individual-level GWAS sample collection, genotyping and data analysis tend to grow as a polynomial function of sample size[7], [8]. Moreover, the exchange of ILD between researchers is non-trivial due partly to data size but increasingly to strict – but essential -data protection regulations, which can limit the scope of collaborative analyses and biological insights gained[9]–[12]. Finally, the substantial computational and financial burden of running massive-scale GWAS, especially for binary disease outcomes, is exacerbating inequity between researchers, typically favouring already well-equipped institutions. There is therefore a pressing need for innovative approaches that help attenuate the increasing resource and financial inequities for conducting contemporary GWAS and to help decide where limited resources should best be allocated.

Here we present in-silico GWAS (isGWAS), a biobank-scalable and computationally highly efficient algorithm to infer genetic regression parameters in case-control GWAS from *j*ust four broadly ascertained cohort-level summary parameters: the counts of cases and controls within a cohort, as well as case and control minor allele frequencies (MAFs). isGWAS is highly parallelisable, exceeding efficiencies of current GWAS analysis tools by several orders of magnitude. Furthermore, we demonstrate that isGWAS yield association summary statistics highly comparable to traditional ILD regression-based approaches through extensive simulations and empirical tests in UK Biobank[13], Biobank Japan[14] and the Psychiatric Genomics Consortium cohort[15]. Owing to the sizeable computational gains, we introduce a heuristic re-sampling algorithm, called the leapfrog re-sampler (LRS), which can confidently extrapolate GWAS results to larger sample sizes, both at a locus or genome-wide scale. Our underlying methodology also leads to several desirable high-utility properties. We release a web tool available to the wider public to conduct customized isGWAS at www.optima-isgwas.com.

## Results

### Genome-wide association testing from sufficient statistics

isGWAS assumes disease-variant associations can be evaluated via a logistic-link function and, similar to widely used methods[7], [8], uses a Firth ad*j*usted maximum likelihood procedure and Newton-Raphson solver to estimate genetic effects, standard errors and association *p*-value[7], [8], [16]. isGWAS’ notable advance is based on the insight that the Newton-Raphson procedure can be simplified so that: (a) elements of the Fisher information matrix and score function vector are collapsed by taking expectation over the empirical or *a priori* distribution of a genetic variant; and (b) sufficient statistics – a specific type of summary data - are used as input variables in the score function (see **Methods** for details). We provide several options to initialise the Newton-Raphson algorithm[17], [18] that improve computational performance and reduce analysis time (**Supplementary Information**). In brief, let *y*_i_ denote disease status for the *i*-th individual and *g*_*ij,M*_ denote the *j*-th genetic variant under model *M*(e.g., additive, recessive or dominant). The sufficient statistic triple used by isGWAS is:

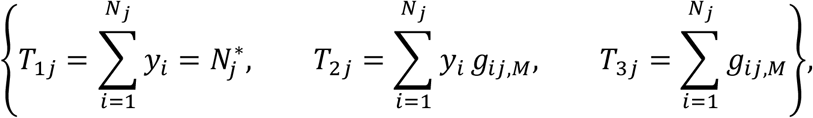

where *T*_1*j*_ is the total number of cases for variant *j, T*_2*j*_ is the covariance between the outcome *y* and genotype *g* for variant *j* under model *M*, and *T*_3*j*_ is the minor allele count for variant *j* under model *M*. For each variant, data can be provided as either: the sufficient statistic triple {*T*_1*j*_, *T*_2*j*_, *T*_3*j*_} plus sample size *N*_*j*_ (necessary for computing standard errors) or separately, on assuming Hardy-Weinberg equilibrium (HWE), 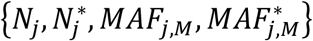. Default GWAS analyses assume HWE, making input data widely available[13]–[15], [19]–[21] for researchers to perform isGWAS, replicate or further expand on classical GWAS analyses (**Methods**). If MAFs for cases and controls are supplied, isGWAS will automatically convert to the pair 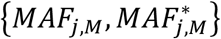 (**Methods**). After convergence, which is guaranteed for most scenarios by a re-initialization approach (empirically all scenarios converged using isGWAS-Firth), the estimated genetic effect parameter 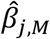 and standard error 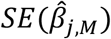 are used to construct Wald *p*-values (**Methods**). Additional options include a sample-level likelihood ratio-test or *p*-values computed using sandwich-robust standard errors (**Supplementary Methods**). A simplified illustration highlighting differences and computational advantages of isGWAS against ILD-based genetic association analyses is summarised in **<u>Figure 1</u>**.

### isGWAS reliably identifies genetic associations across cohorts and diseases

We benchmarked isGWAS in real-data settings and performed simulation studies to compare isGWAS performance and results relative to several existing individual-level data (ILD)-based approaches. Our assessments broadly fall into two categories: (1) methods which require ILD, i.e., REGENIE[8], logistic and Firth corrected regression[16], and (2) approaches which do not require ILD directly, i.e., the logistic ad-hoc estimator[17] and Fisher’s Exact Test (FET)[22]. We note FET was successfully leveraged for efficient large-scale GWAS analyses recently[23]. Using data from UK Biobank (UKB), we first assessed isGWAS performance against the popular ILD based regression approach REGENIE[8] by deploying both methods for analyses of seven diseases some of which were previously used for establishing GWAS methodology[5], [7], [8]. Second, we evaluated isGWAS’s ability to replicate 309 significantly associated variants from the Biobank Japan (BBJ) meta-analysis of 30 diseases[24] using only the published sample-summaries, i.e., without access to ILD, per variant and disease pair. Lastly, by considering multiple iterations of nested schizophrenia meta-analysis GWAS[25]–[27] we assessed isGWAS’s ability to accurately predict genomic regions and significant novel associations. The isGWAS sample size of the meta-analysis from 2014 was virtually expanded to match the numbers from the larger 2022 GWAS whilst holding constant the allele frequency and prevalence information in 2014 cohort.

#### Performance against ILD based regression GWAS in UK Biobank

We compared isGWAS results and computational performance against the state-of-the-art ILD-based method REGENIE for seven diseases in UKB: asthma (IC10:J45); atherosclerosis (IC10:I25); colon cancer (IC10:C18); hypertension (IC10:I10); glaucoma (IC10:H40); stroke (IC10:I63); and thyroid gland cancer (IC10:C73). Case-control ratios varied from 1:2 in hypertension to 1:669 in thyroid gland cancer across the diseases (**Supplementary Table 1a**), allowing for the review of performance in near balanced to highly imbalanced case-control settings. To help attenuate the possible influence of confounders when deploying isGWAS, particularly sample relatedness and population structure, we describe and apply additional data quality control (QC) steps before computing the required sample-level sufficient statistics (**Methods**). After additional QC, the total sample size analysed was ∼335,000 individuals per disease (**Supplementary Table 1**). This sample was used to perform and contrast analyses in both isGWAS and REGENIE. We review approaches which leverage additional insight from any removed samples in the **Discussion**. For each of the seven diseases, we applied isGWAS (no covariates) and two-step REGENIE with Firth correction (including the covariates age, sex and ethnicity principal components) to ∼11 million autosomal variants. Results are presented in **Table 1, Figures 2-3, Supplementary Figures 1-15, Supplementary Tables 1-9 and Supplementary Files 1-2**.

**Table 1.** Accuracy, true positive rate, false positive rate and false discovery rate of isGWAS using REGENIE results as gold-standard and a threshold of p = 5e − 08 as classification rule. Results are obtained on all 11,079,229 variants used for the analysis of seven diseases in UK Biobank without clumping/finemapping.

| **Disease (ICD code)** | **Case:control ratio** | **TPR** | **FPR** | **Acc** | **FDR** |
| --- | --- | --- | --- | --- | --- |
| **Hypertension (I10)** | 1:2 | 0.625 | 0.000026 | 0.99961 | 0.041 |
| **Asthma (J45)** | 1:6 | 0.982 | 0.000028 | 0.99994 | 0.016 |
| **Atherosclerosis (I25)** | 1:9 | 0.886 | 0.000011 | 0.99995 | 0.043 |
| **Glaucoma (H40)** | 1:26 | 0.891 | 0 | 0.99997 | 0.036 |
| **Stroke (I63)** | 1:56 | NA | 0 | 1 | NA |
| **Colon Cancer (C18)** | 1:94 | 0.944 | 0 | 0.99999 | 0.037 |
| **Thyroid Gland Cancer (C73)** | 1:669 | 1 | 0 | 0.99999 | 0.051 |

**Figure 1.**
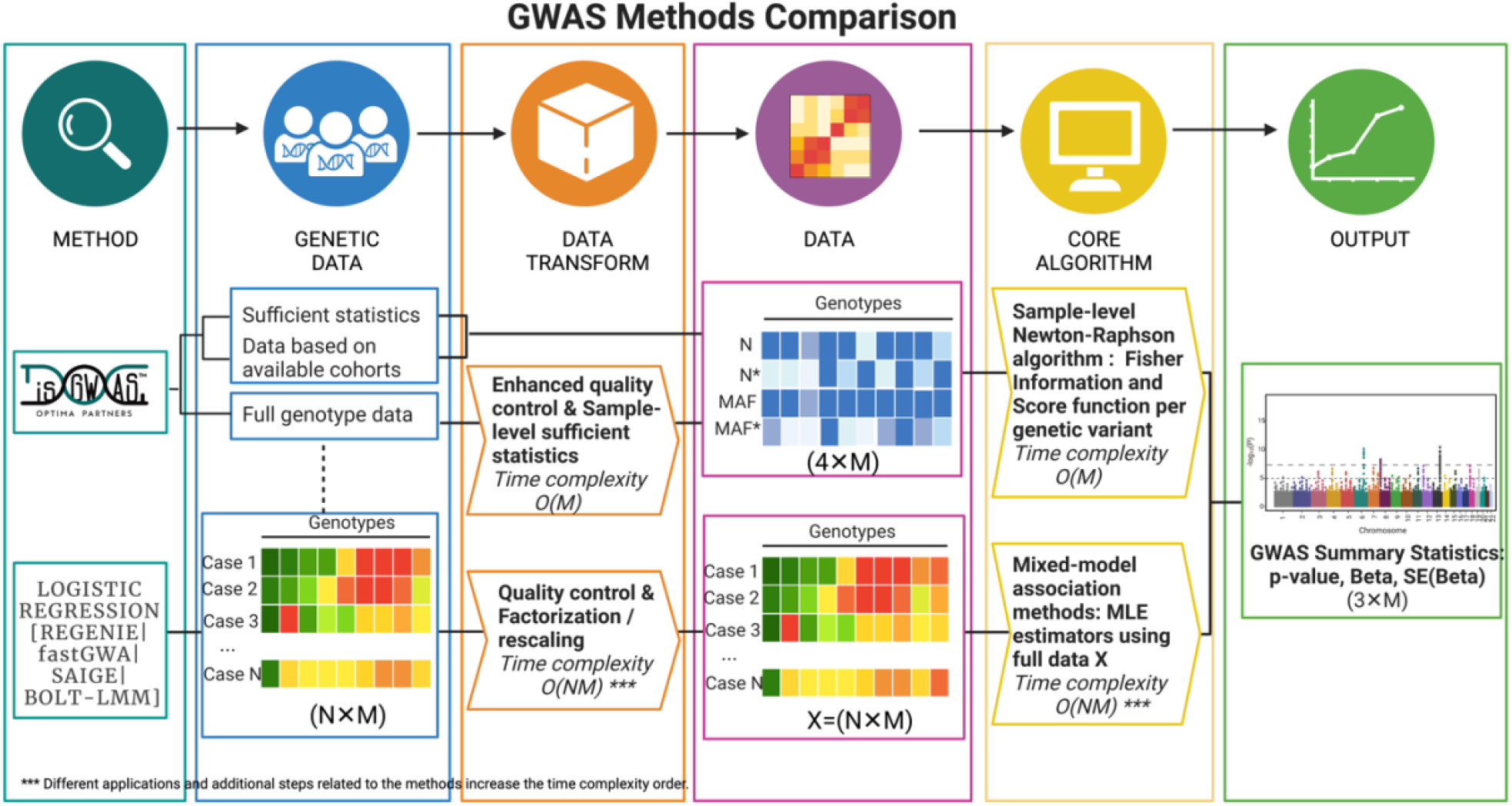
Diagram highlighting main differences between isGWAS and other GWAS approaches.

**Figure 2.**
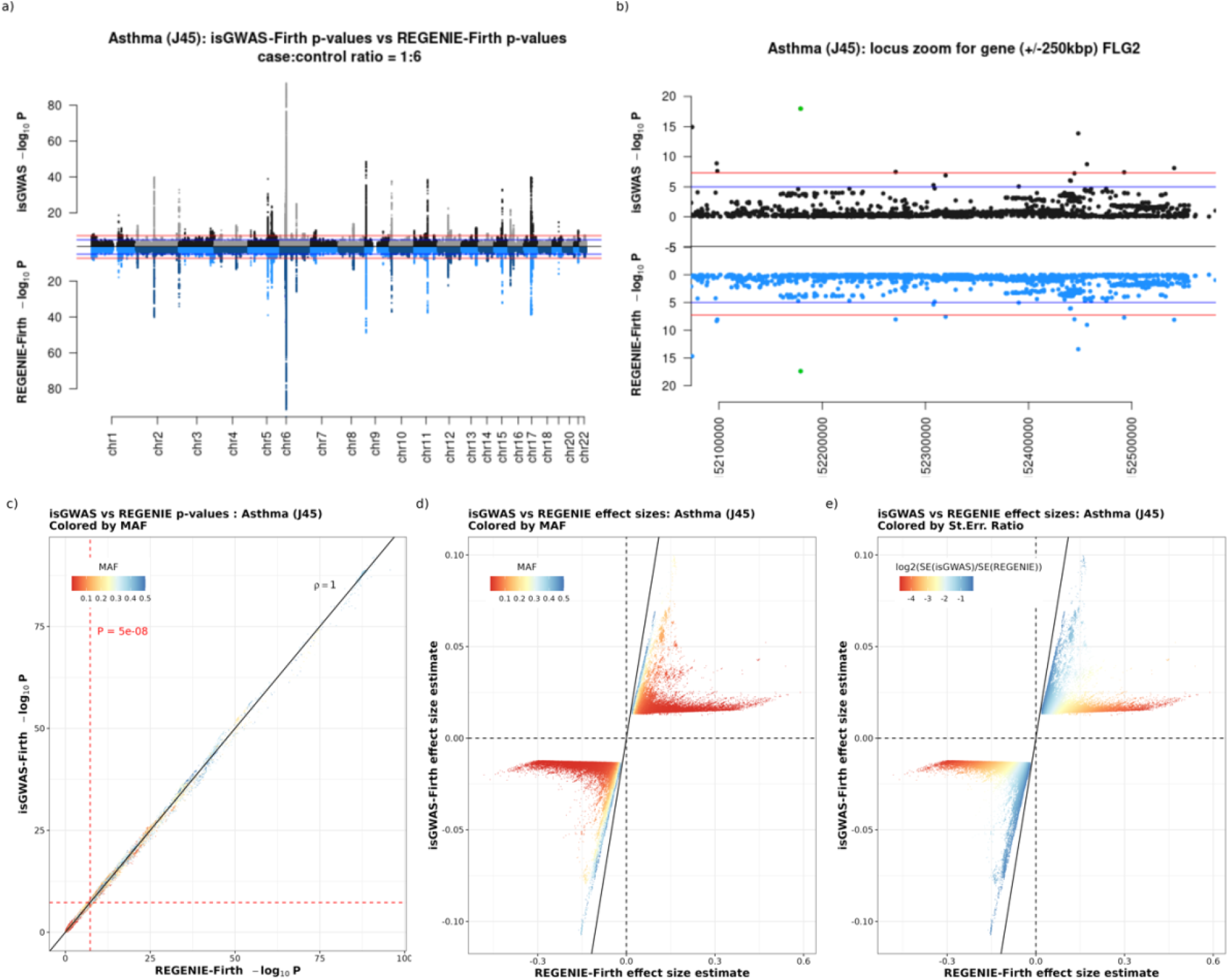
Comparative results for Asthma (IC10:J45) from UK Biobank. Subplot (a) is a mirror Manhattan plot comparing −log10 P values for isGWAS and REGENIE-Firth and subplot (b) is a locus zoom of the gene FLG2 region +/-250kbp on chromosome 7. Subplot (c) plots −log10 pvalues for isGWAS and REGNIE-Firth with the standard threshold P-value indicated colored by population-level MAF. Subplots (d) and (e) showcase β− βeffect size estimates for variants with p-values<0.05 and are coloured by population-level MAF and 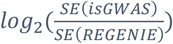.

Across all seven traits tested we observed close to perfect consistency between REGENIE and isGWAS association results, as illustrated in the mirrored Manhattan and *p-p* plots for asthma (**Figure 2a**) and other diseases (**Supplementary Figures 1-6**). Concordance between REGENIE and isGWAS is further validated by benchmarking accuracy (**Table 1**) and Pearson correlations between estimated *p*-values (*cor(p*_*isG*_, *p*_*R*_)) > 0.94 at *log*_10_ scale (**Supplementary Tables 2-3**). The results are consistent for varying prevalence levels (**Supplementary Figures 1-6**) and are not affected by covariate ad*j*ustment (**Supplementary Tables 2-3**). The consistency translated to the regional locus level. This is exemplified by a locus zoom plot of the *FLG2* gene region for asthma (**Figure 2b**) where isGWAS not only nominated the identical GWAS lead variants but also largely recapitulated the overall association pattern identified through REGENIE. This observation is consistent across the lead independent loci from the asthma GWAS (**Supplementary Figure 7**) and translates to all other diseases studied (**Supplementary Figure 8**). For a comprehensive numerical comparison of association results, we took REGENIE derived *p*-values as the ground truth, retaining all SNPs with *p* < 0.01 and setting the true positive threshold as *p* < 5 × 10^−8^ (excluding stroke (**Supplementary Figure 4**) which did not yield any significant associations). We computed the accuracy, false positive (FPR), true positive (TPR) and false discovery rates (FDR) of isGWAS (**Figure 3a** and **Table 1**). Accuracy of isGWAS was ≥ 99.98% for each disease, highlighting excellent overall correspondence between methods. The FPR was low, i.e., *FPR* ≲ 10^−5^, and TPR was generally good at > 88% - excluding hypertension which had a *TPR* = 0.63. The FDR was below ≤ 5% for each disease, revealing that the positive predictive value of isGWAS was greater than 95%.

**Figure 3.**
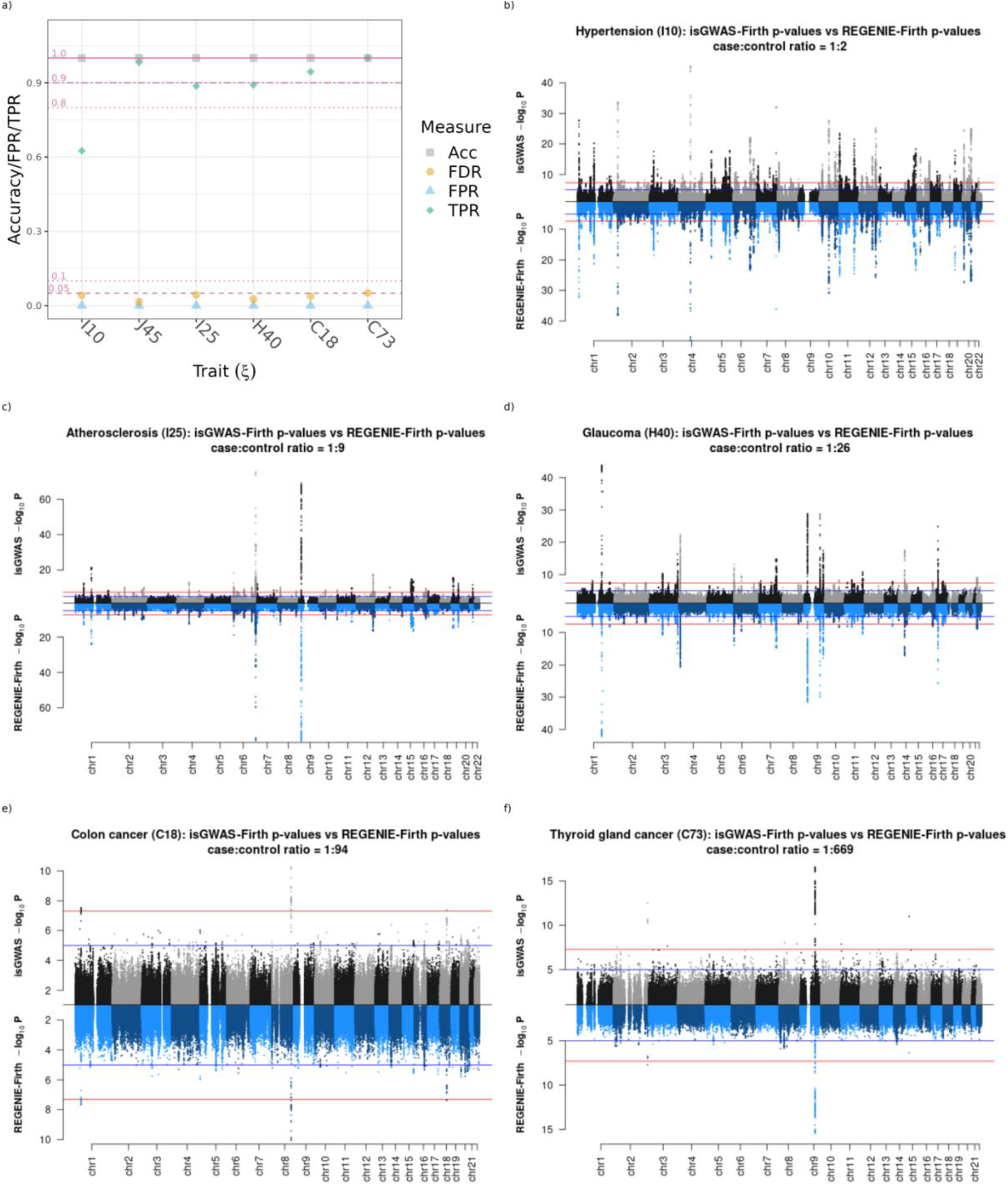
a) Accuracy/TPR/FPR comparing REGENIE-Firth and isGWAS results, where p = 5e − 08 threshold has been used as indicator for correct classification accuracy. Results are obtained on all 11,079,229 variants used for the analysis without clumping/finemapping. See Supplementary Table 1 for full results. Manhattan plots b), c), d), e) and f): Comparative results for five diseases from UK Biobank. Mirror Manhattan plots comparing −log_10_ P values for isGWAS and REGENIE-Firth for six different diseases obtained from UK Biobank. Stroke was excluded from the analysis due to no variants passing significance threshold.

**Figure 4.**
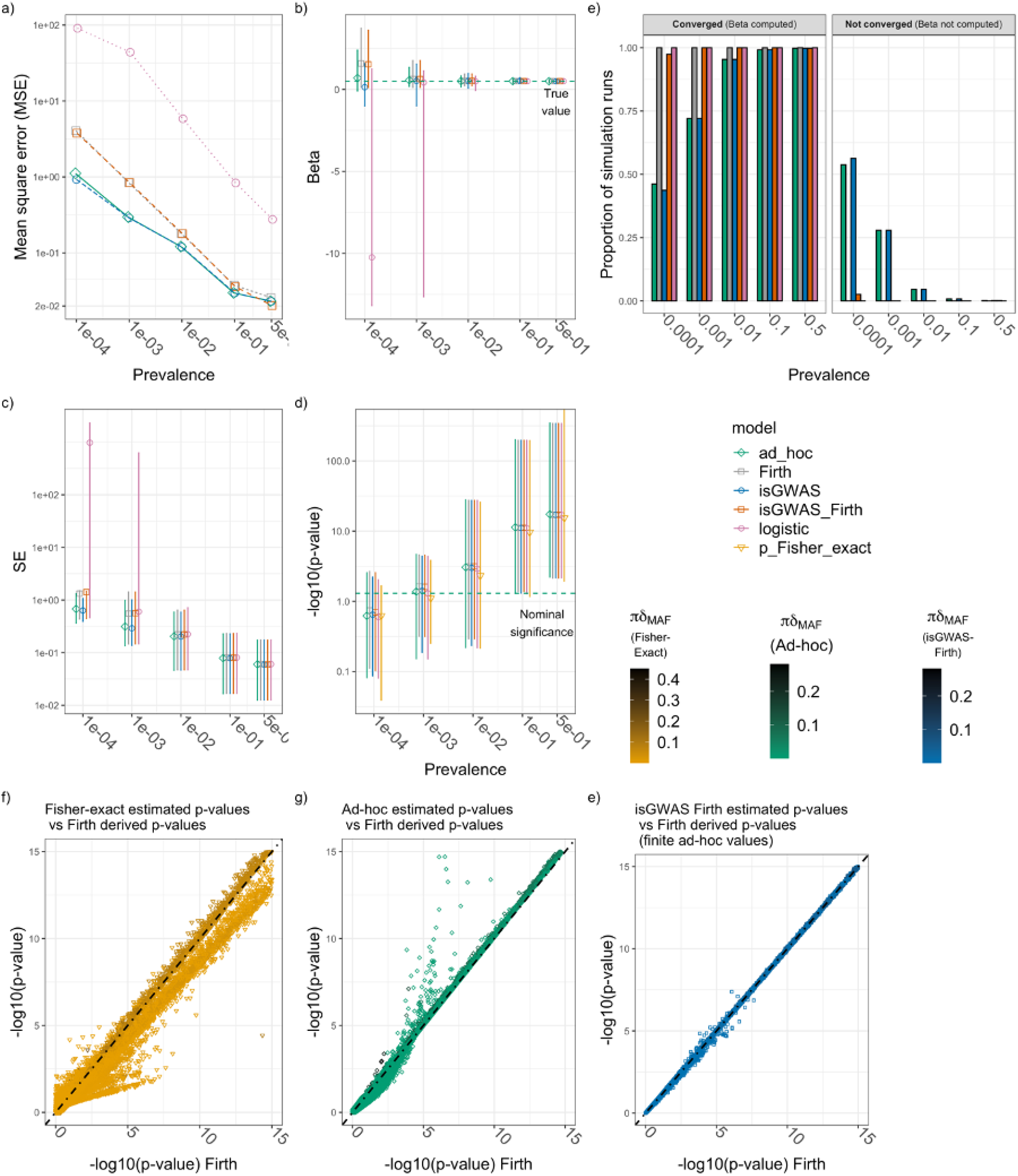
Simulation I results. Clockwise from top: a) Mean square error; b) distribution of estimated beta values; c) distribution of associated standard errors and d) distribution of -log10(p-value), for each model - logistic regression, firth regression, ‘isGWAS’, ‘isGWAS_Firth’ and, for p-values only, Fisher’s Exact Test - and specification of disease prevalence. Panel e) presents the relationship between Firth regression derived -log_10_(p-value), along the horizontal axis, and the corresponding ‘isGWAS_Firth’ and Fisher’s Exact Test (FET) computed values, on the vertical axis. Results are presented in the range [0,15] as FET regularly failed to converge for very small p-values and coloured according to value of πδ_MAF_, where π denotes prevalence and 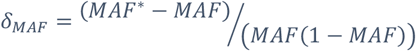. In panels b)-d) a point denotes the median value and error-bars the first and ninth deciles of the range.

Importantly, isGWAS and REGENIE results differed for two broad categories: (i) the estimation of genetic effect sizes; and (ii) computational performance. When non-confounding covariates are excluded (*β*_no_cov_) or included (*β*_cov_) in a model, previous and extensive investigations of effect size estimates in logistic regression deduce that |*β*_no_cov_| ≤ |*β*_cov_|, i.e., regression estimates are smaller in magnitude when excluding covariates but the null-hypothesis of no association is maintained[28], [29]. Overall, we replicate these results in our analyses. isGWAS computed effect sizes are smaller in absolute value, but largely concordant with covariate-ad*j*usted REGENIE. Moreover, we fail to re*j*ect the null hypothesis for the same variants almost always between methods - suggesting that the isGWAS QC helps attenuate issues of population confounding (**Figures 2d-e, Supplementary Figures 1d-e – 6d-e, 9, 10 and 11**). An investigation of the performance of isGWAS without removing related individuals highlights potential expansion of isGWAS beyond the recommended QC (**Methods, Supplementary Information**), but further investigation – possibly leveraging the re-sampling potential of isGWAS - is required on the reliability of isGWAS in family-based cohorts and ethnically diverse populations (**Supplementary Figures 12-13, Supplementary Tables 5-7)**. In our full-QC analyses, all estimated effects between isGWAS and REGENIE were observed to be in the same direction, and the correlation between estimates was on average *cor*(*β*_*isG*_, *β*_*REGENIE*_) ≈ 0.7 (**Supplementary Tables 1-3**). The relative drop in the correlation between effect estimates (≈ 0.7) and p-values (≳ 0.94) is anticipated[28] and can be explained on noting that, across all diseases more precise effect estimates (i.e., those with smaller standard errors) have stronger concordance between approaches (**Figure 2e and Supplementary Figures 1d-6d**). Overall, we found that at least 98% of isGWAS and REGENIE confidence intervals (CI) overlap, (**Supplementary Table 4**). When effect estimates are viewed as a function of MAF, the absolute value of REGENIE-derived estimates seemingly increase (along with standard errors) as MAF decreases across all scenarios. This contrasts with isGWAS where the relationship between MAF and effect size is less clear: fewer variants with low MAF are associated with relatively larger effect sizes. However, the correspondingly narrower standard errors guarantee the same significance p-values as REGENIE. The isGWAS derived distribution of effect sizes is consistent with the hypothesis of a flattened heritability distribution under negative selection[30]. Genomic inflation computed from isGWAS results across all analyses was on average ≈ 1.07 and ranged between (0.94, 1.26) which was similar to REGENIE with average ≈ 1.1 and range (1.01,1.3) (**Supplementary Figure 14** and **Supplementary Table 8**). We deploy isGWAS with genotype imputation in our primary analyses and, as secondary sensitivity analyses, without imputation. Our investigation reveals some surprising results. Imputation occasionally led to changes in MAF between cases and controls such that estimated genetic effects switched sign (i.e., effect direction) relative to results computed from non-imputed data (**Supplementary Figure 15, Supplementary Table 3, Supplementary Files 2-3**). The approach might be used to efficiently flag ambiguous significant results in analyses that are the result of the missing values imputation strategy (mean-imputed in the case of REGENIE).

Finally, the computational gains of isGWAS relative to REGENIE Step 2 are striking: a full genome-wide association assessment for each disease took approximately 4 minutes using isGWAS and, on average over different prevalence, this is around 1,300 times faster than a like-for-like assessment using REGENIE Step-2 (**Figure 7, Supplementary Table 9, and Supplementary File 4**).

#### Replicating significant associations in Biobank Japan analyses

Using only publicly available summary information from Biobank Japan (BBJ), i.e., without access to ILD, we looked to compare and replicate BBJ GWAS results across 42 diseases[24]. We considered 309 variants that were identified in [24] as genome-wide significant (*p* < 5 × 10^−8^) across 30 of the 42 diseases. Our results reveal very close alignment between isGWAS computed associations and those of [24] - correlation between p-values at *log*_10_ scale was *cor*(*p*_*isG*_, *p*_*BBJ*_) = 0.98 with 92.2% of isGWAS computed genetic effects within the 95% CI of the original study (**Supplementary Figure 16)**. Using the published study-level results[24] as the ground truth, isGWAS demonstrated good sensitivity and specificity (**Supplementary Figure 17**). We alternatively assessed performance when setting more stringent significance thresholds - returning near identical conclusions when classifying variants at *p* < 9.58 × 10^−9^ (used in the original publication). Results for X-chromosome variants in males and females were similarly concordant (results not presented).

### isGWAS model validation using simulations

We generated simulated datasets to assess performance of isGWAS - with and without Firth correction - against a variety of classical methods which either: (a) do not require ILD, the logistic ad-hoc estimator[17] and Fisher’s Exact Test[22]; or (b) require ILD, logistic and Firth corrected regression[7], [8], [16]. We perform two simulation studies (**Figure 4, Supplementary Figure 18, Supplementary Information)**. isGWAS-Firth outperformed all other approaches in terms of either computational cost or robustness of results over the range of scenarios considered. It is well documented that computational performance is reduced when using Firth’s bias correction in ILD regression analyses[7], [8], [16], we discover, however, that no-ILD isGWAS-Firth regression has significantly improved performance relative to uncorrected isGWAS (**Supplementary Table 21**). As anticipated[23], when disease prevalence is rare (i.e., π ≤ 0.01) parameter estimates computed using non-Firth corrected ILD regression were unreliable. The MSE and distribution of parameters estimated via ILD logistic regression were often orders of magnitude poorer than other methods (**Figure 4a-c**). **Figure 4f-h** highlights the chronological evolution of no-ILD *p*-value estimates, from Sasieni’s logistic ad-hoc estimator (1997)[17], Fisher’s Exact Test (1922)[22] to isGWAS-Firth, illustrating improvements in estimation via successive approaches. See **Supplementary Information** for detailed results review.

### Leapfrog re-sampling: using isGWAS to extrapolate variant association results to larger sample sizes

When ILD are available, the computational benefits of isGWAS make it possible to deploy resampling approaches to estimate empirical effect sizes, *p*-values and corresponding confidence intervals, previously considered computationally daunting in GWAS[31], [32]. We extend the idea by introducing a heuristic leapfrog re-sampling (LRS) algorithm to help forecast future results in larger hypothetical GWAS sample sizes (**Methods**). The LRS is summarised in three key steps: (1) specify a target sample size along with the number and size of sub-samples to be generated; (2) (leapfrog-step) compute sufficient statistics in the sub-samples and re-scale the estimated number of cases and controls to match the larger target sample size; and deploy isGWAS in each leapfrog sample to recover a distribution of association *p*-values over the collection of sub-samples. In our testing of the LRS, we use the median *p*-value as a generally robust estimate of a target p-value (weighted or distribution-based summaries can alternatively be considered). Thus, the LRS leverages variation in both genotype and disease status between individuals in the current sample to help predict updates of parameters after adding new samples. Despite perceived similarities, traditional GWA power calculators[33] and the isGWAS-LRS are different. isGWAS-LRS does not require input of case-control ratios, heritability (i.e., beta estimates) or type-I error rates. Instead, multiple regression analyses are combined to forecast and test parameter estimates in expanding sample sizes.

We run the leapfrog re-sampler in both simulation and real-data settings, informed by the seven studied diseases in UKB (**Methods and Supplementary Information**). We evaluate performance over a range of initialisations, starting from a 10% increase to a maximum of 100% (i.e., 2-fold) increase in GWAS sample size relative to the current actual UKB sample size. Results are presented in **Figure 5, Supplementary Tables 10-11**. As is standard, we assume a true positive association of p<5×10−8 in the target sample. Our results in simulated scenarios (**Figure 5f**) reveal that: when doubling sample size from *N*_*current*_ = 276,204 to a maximum *N*_*target*_ = 552,408, the accuracy and TPR progressively dropped for subsequent increases in the target sample size, but values for each measurement were typically ≥80% across the range. Our real world LRS analyses of UK Biobank data replicate and further elucidate performance across the six of the seven diseases (**Figure 5a-e**). Using a subsample size of *N*_*current*_ ≈ 135,000 we increased target sample size up to *N*_*target*_ ≈ 270,000, taken as the maximum observed sample size we could benchmark against. For all choices of target sample size, and across each disease, we observe high accuracy rates (≥95%). However, the TPR was sensitive to disease prevalence, reducing monotonically as the target sample size increased. Broadly, TPR remained reasonable (≳ 60%) up to a 2-fold increase in sample size, except for the very rare (case-control ratio of 1:669) thyroid gland cancer. This is due to fewer significant variants being included in the assessment as a result of lower percentage of heritability explained, which can artificially reduce the TPR for each new locus with relatively high odds ratios. Naturally, TPR reduces as a function of decreasing disease prevalence, as re-sampling from fewer cases can increase the variability in MAFs and thus isGWAS forecasting. We note that our theoretical sub-sampling approach had better predictive capabilities, owing to the prevalence preserving sampling strategy taken (**Methods and Supplementary Figure 19**).

**Figure 5.**
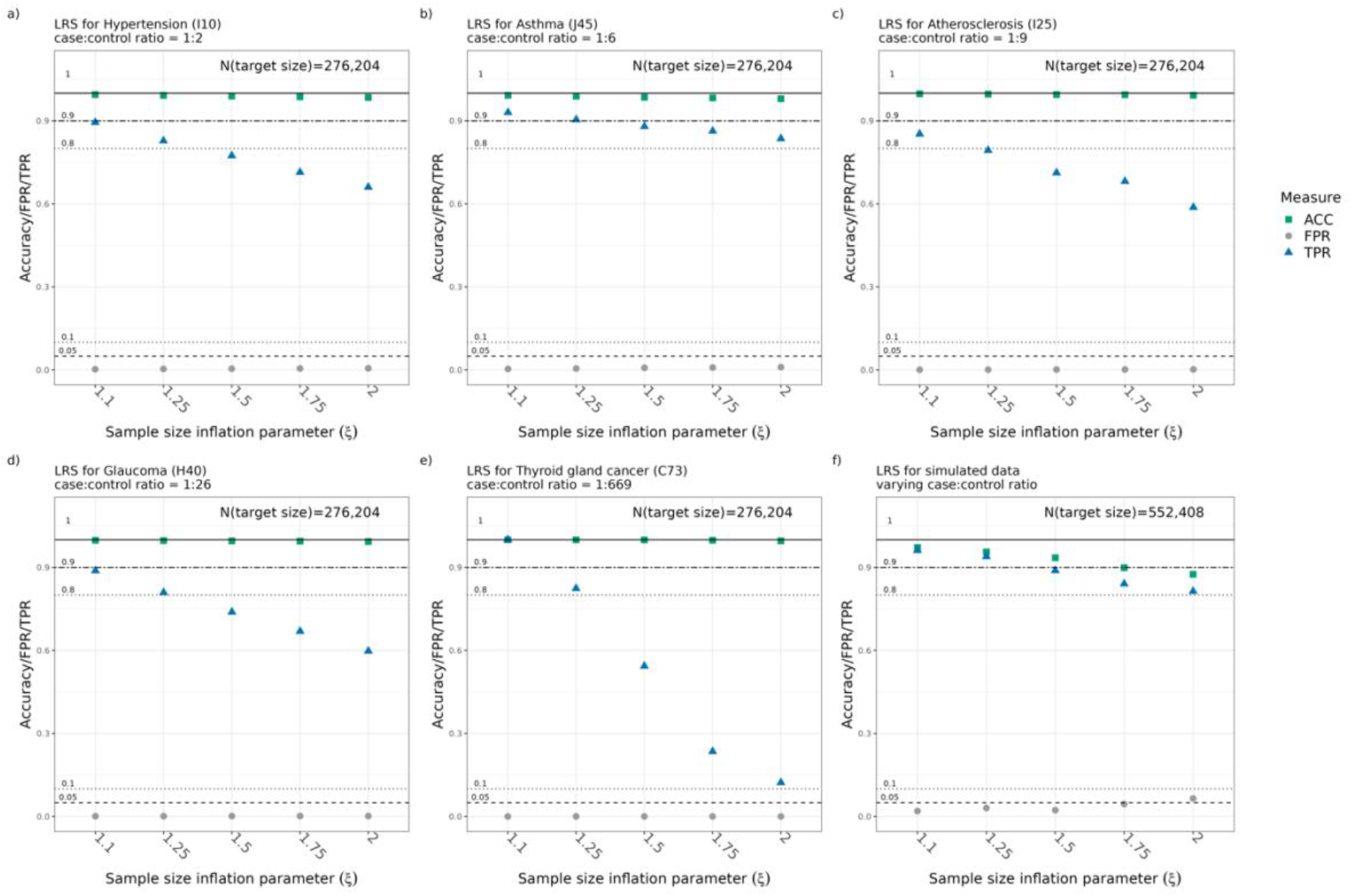
Performance of Leapfrog re-sampler (LRS) benchmarked against results derived from a)-e) the sample of ∼276,204 individuals from UK Biobank with five different diseases and b) a simulated sample of 552,408 individuals (i.e., double UK Biobank sample size). For each value of ξ, we subset the target sample size down to 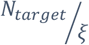 individuals and deploy the LRS to compute predictions for the target sample N_target_. As the maximum number of UK Biobank samples was 276,204, this was taken as the target. For example, when ξ = 2, we subset the full sample to 138,102 individuals and run the LRS to compute predictions of the larger 276,204 sample. We use results from the disease analysis, benchmarking LRS predictions against those computed on the pruned genome sampling uniformly across significance associations, resulting in ∼3500 variants per studied disease. Colon cancer and Stroke are excluded from this figure as they don’t have significant variants or a very low number of such after pruning. In the right panel we generated 1000 simulated datasets under the null of no genetic association or the alternative (see simulation protocol for details).

We also assessed isGWAS’s ability to extrapolate results when ILD were not available, using a highly constrained version of the leapfrog re-sampler (**Supplementary Information**). In this scenario, MAF and disease prevalence per variant are fixed, computed from the maximum current sample (i.e., without sub-sampling), and the number of cases and controls are proportionately increased to match the target sample size. We did this for two GWAS of schizophrenia: (a) 2014 analyses with up to *N* = 77,096 (cases = 33,640, controls = 43456) European ancestry individuals[25]; and (b) the larger (and future) 2022 analyses with up to *N* = 130,644 (cases = 53386, controls = 77258) European ancestry individuals[27]. We treat the 2014 study as the current sample size and the 2022 sample size as the target future state, which we benchmark predictive performance against. The studies were selected because, for each variant *j*, necessary data to run isGWAS, i.e., 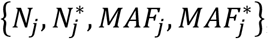, were made publicly available. Note these data are pooled estimates, computed across all European cohorts. Despite not accessing ILD, our results reveal reasonable concordance between isGWAS 2014 extrapolated results and the published analyses of 2022 (**Figure 6, Supplementary Figures 20-21, Supplementary Tables 13-16**). Like our Biobank Japan analyses, we also used a more stringent significance threshold (*p* < 10^−10^) to help attenuate false positives, observing improved overall performance by recovering a good TPR ≥ 70% (**Supplementary Table 13**). We do not report FDR as these cannot be accurately computed when filtering results based on a p-value inclusion/exclusion threshold. Of the overlapping 608 clumped variants considered, isGWAS-LRS identified 136 associations that were not yet deemed GWAS significant (i.e,. *p* > 5*e* − 8) in the 2014 study but later identified as significant in the 2022 study. Moreover, of the 436 significant associations predicted by isGWAS, 75% overlap with observed significant associations in 2022. isGWAS predicted an additional 74 associations as significant that were not significant in 2022 – of those 52 were near the significance threshold with *p* < 9*e* − 07. There were 121 variants not correctly predicted by the 2014 cohort. This could be due to increased ethnical and relatedness heterogeneity in the 2022 cohort that was not present in the 2014 analysis.

**Figure 6.**
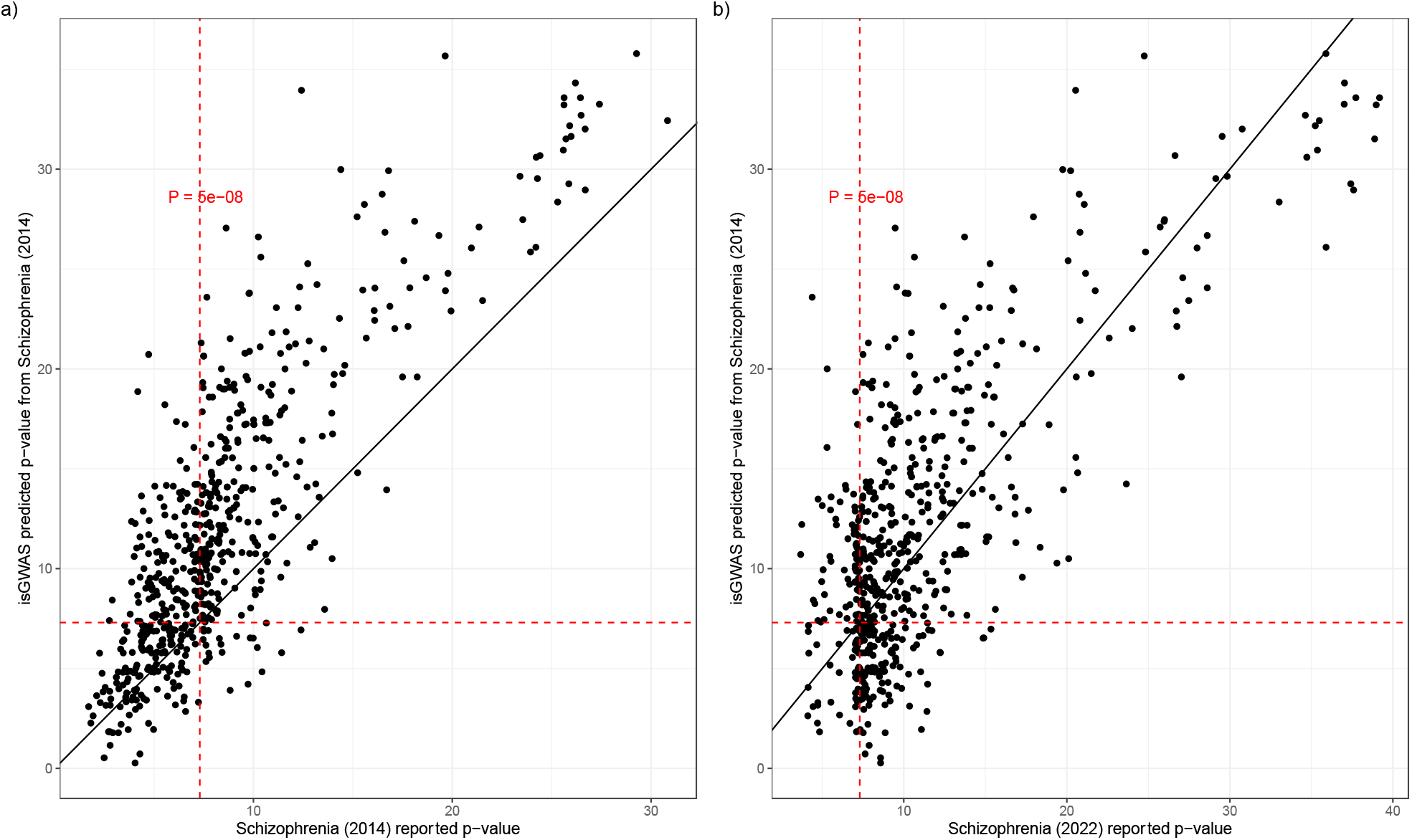
Prediction results for Schizophrenia (2022) using data from Schizophrenia (2014): population-level information for 608 significantly associated loci (P<1e-07) obtained from clumping with parameters (R^2^ = 0.2, p_1_ = 1e − 7, p_2_ = 1e − 7) has been used to infer p-values. a) The figure compares reported GWAS Schizophrenia (2014) p-values and isGWAS predicted p-values using population-level information from Schizophrenia (2014) matching for the larger 2022 cohort size. b) The figure compares reported GWAS Schizophrenia (2022) p-values and isGWAS predicted p-values using population-level information from Schizophrenia (2014) matching for the larger 2022 cohort size. The dashed red line represents the threshold P=5e-08.

**Figure 7.**
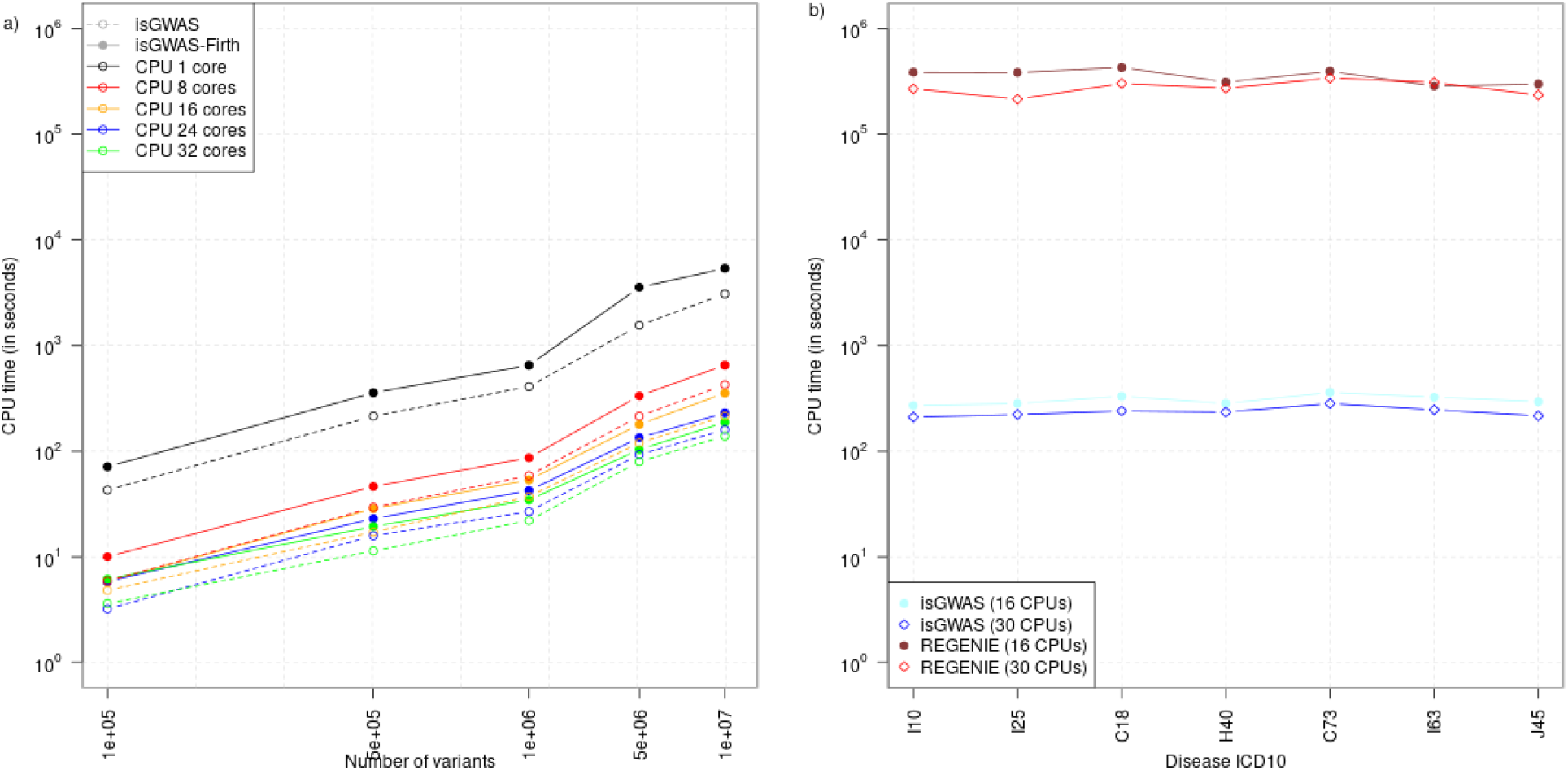
a) Computational CPU time (in seconds) for an increasing number of variants. The results compare performance of isGWAS with and without Firth distributed over different number of CPU cores. The data was obtained from UK Biobank ICD10:C73 disease with low disease prevalence (case-control ratio = 1:669). The x- and y-axis are on log_10_ scale. b) Computational CPU time (in seconds) for seven UK Biobank diseases on 11,079,229 variants for ∼335,000 individuals. We compare the performance of isGWAS running on 16 and 30 CPUs vs the performance of REGENIE Step 2 running on 16 CPUs (with 16 threads) and 30 CPUs (with 30 threads). The computation is performed on the same machine. The y-axis is on log_10_ scale.

### Computational performance and convergence details

isGWAS is an iterative algorithm whose convergence (i.e., ability to estimate model parameters) depends on several tuning parameters (**Methods**). Using default parameter settings, isGWAS-Firth converged in all real-data and simulated scenarios tested (**Figure 4e and Supplementary Tables 17, 19-20**). Convergence was achieved in around 0.001 seconds per variant (**Supplementary Table 21**) on a 2.4 GHz 8-Core Intel Core i9 processor. The non-Firth corrected isGWAS algorithm may require more iterations, particularly for diseases with lower prevalence (e.g., case:control ratio of 1:94 and lower) which included scenarios where convergence was not achieved (**Figure 4e, Supplementary Tables 17-20, Supplementary Information**).

When distributed over 32 CPU cores on a high-performance cluster, Firth-corrected isGWAS analysed a single disease from UK Biobank across ∼11 million SNPs and for ∼335,000 individuals in ∼4 minutes (**Figure 7, Supplementary Tables 9 and 18**). This means that isGWAS-Firth can perform around 1,500 disease GWAS for every one GWAS performed using an alternative methodology. The same analysis with a small number of CPU cores was completed in tens of minutes using isGWAS-Firth (**Figure 7**). Further computational gains at larger sample sizes will likely be achieved as ILD methods can scale poorly with sample size, whereas isGWAS has near fixed computational cost at any size. As isGWAS currently computes associations for each variant independently, additional improvements such as parallelisation are possible. Full details are available in **Supplementary File 4**.

## Discussion

In this study, we developed isGWAS, an efficient, biobank-scalable method for genetic association testing which can: (a) compute regression parameters and test for a variant-disease association in real-time (i.e., approximately one millisecond) for any sample size; (b) bypass the need to run large-scale GWAS using high-performance computing facilities owing to ultra-low system resource demands (i.e., runtime and memory); and (c) infer GWAS results from virtually enlarged sample sizes using a novel re-sampling procedure. The isGWAS algorithm design allows analyses to be run without the need to hold or access individual-level data (ILD) directly, thereby providing a single methodological framework to utilise a wide range of data sources such as published summary-level data from biobanks and repositories.

isGWAS draws inspiration from classical methodologies to overcome significant computational bottlenecks associated with massive-scale analyses. The practical simplicity and quick runtime of classical approaches have seen them deployed in a recent large-scale analysis[23]. Rather than using ILD, as contemporary GWAS regression analyses do, isGWAS distils the required input data down to sufficient statistics – a low-dimensional summary of ILD that captures all necessary information required to compute a genetic-disease association model parameter. In combination with modifications to the Newton-Raphson procedure, used to estimate model parameters in a logistic regression, our use of sufficient statistics dramatically reduces the computational time for disease association testing relative to existing methods. Achieving up to a 1,500-fold improvement in computational runtime, when benchmarked against a state-of-the-art GWAS tool, isGWAS reduced time to genome-wide insight from several days down to ∼4 minutes. Thereby unlocking potential for massive scale exploration of genetic-disease associations in real-time and making feasible the routine assessment of thousands of disease endpoints and studies. Computational bottlenecks associated with existing GWAS methodologies are fast approaching. Analyses of resources such as UK Biobank WGS data, the emerging massive cohorts of the Global Biobank Initiative[1], and Our Future Health[34] are expected to push current GWAS tools to their system resource limits with significant associated time-to-insight penalties. Conversely, with no computational sensitivity to sample size, as the number of variants assessed and sample sizes continue to increase, the relative savings and benefits of isGWAS can be expected to grow.

To attenuate possible issues of confounding and population stratification, we propose that additional QC-steps are performed before computing sufficient statistics for isGWAS. In our analyses, these steps reduced our UK Biobank sample size from ∼408k individuals (used in the original testing of REGENIE[8]) down to a more homogeneous sample of ∼335k individuals used to generate and compare results from isGWAS and REGENIE. Our results reveal an often-striking concordance between approaches genome-wide as well as at the regional locus level. The reduction in sample size was compensated by the isGWAS leapfrog re-sampler (LRS), which we demonstrate efficiently helped extrapolate GWA results onto larger sample sizes (up to 2-times). While we note sensitivity of an LRS extrapolation to disease prevalence, across the range considered the TPR and FPR were well calibrated to at least a 1.5-fold increase in sample size. The LRS might therefore be leveraged to aid GWAS cohort design, for example to quantify the potential benefit of sampling more participants with a disease of interest against cost. In our analyses of the PGC Schizophrenia cohort, we deployed a highly restricted (i.e., no ILD) version of the LRS: forecasting results from a smaller 2014 cohort[25] onto a sample size that matched a future 2022 study[27]. Despite no guarantees of sufficiency, isGWAS LRS identified 75% of significant variants that were later identified in the larger 2022 cohort (almost double the size) while maintaining a low FDR. Unlike extrapolation via the ILD leapfrog re-sampler, this naïve extrapolation does not account for differences in the MAF of cases and controls between 2014 and 2022 data. Regardless, the above findings highlight potential for isGWAS to furnish reasonable forecasts of future results without accessing ILD directly. Our recent predictions from FinnGen consortium data[18], [35] provide confidence that the isGWAS algorithm is applicable also to multi-ethnic GWAS through analyzing each ethnicity separately and combining results in a meta-analysis, as it is common practice[36].

Beyond, isGWAS can be applied to help address routinely asked questions about future scenarios and evaluate enrichment contribution of biobanks to disease-specific associations[35] or to protein-specific variant associations[18], particularly in the rare spectrum. isGWAS-Firth provides a timely, rapid regression-based analysis of common, rare and ultra-rare variants. Unlike ILD-based analyses, where Firth’s correction significantly increases computational time[7], [8], there is no computational penalty when using Firth’s correction in the isGWAS framework - in fact, we observe improved computational performance. The advantage of considerable improvements in computational runtime is that it allows for the introduction of forecasting, re-sampling and other non-parametric techniques - the LRS being one example. These might widen robust association testing strategies as well as provide new avenues to tackle confounding or population sub-structure. For now, we envisage the possibility that the wider human genetics community routinely compute and make available the sufficient statistics, i.e. MAF in the cases and the cohort, and the corresponding sample sizes per variant, toward a publicly available, privacy compliant, data asset. In addition to avoiding the need for expensive high-performance computing facilities and memory intensive data storage, the data asset might enhance meta-analyses and biological insight, improve equitable access, and enable faster collaborations between teams and help bridge financial and resource gaps between institutions and research groups internationally.

## Methods

### Disease SNP association model

Let *S*_*M*_∈ {1,2} be the maximum number of copies of the effect allele for an individual under model *M*∈ {*A, D, R*}, where *A* denotes an additive model, *R* a recessive model and *D* a dominant model, i.e.,

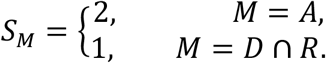

Furthermore let,

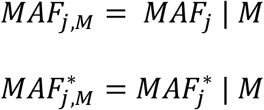

where, for a given model *M, M*AF_*j,M*_ is the minor allele frequency for variant *j* in the sample, ancestry, or population and 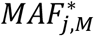 the minor allele frequency in the cases. We let *Y* denote disease status and *G*_*j*_ the *j*th genotype in the sample. For convenience we write *G*_*j,M*_= *G*_*j*_|*M*. It is assumed that the outcome model for *Y*, conditional on *G*_*j*_, is given by:

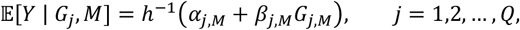

where, conditional on model *M*, the pair {*α*_*j,M*_*β*_*j,M*_} denote the intercept and genotype effect and *h* is a function linking the outcome to genotype *G*_*j,M*_ for all *j* = 1,2, …, *Q* genetic variants considered. In deriving the isGWAS estimation procedure we assume that *h* is the logit function, i.e.,

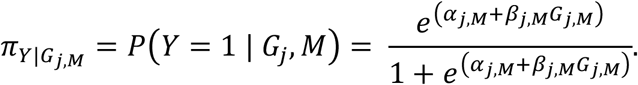

The isGWAS methodology can, however, be broadened to other link functions and outcome types. We take 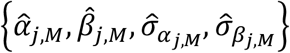 to be sample based estimates of the intercept *α*_*j,M*_ and coefficient *β*_*j,M*_, and their associated standard errors 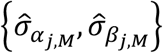. We allow the genetic effect *β*_*j,M*_ and genotype *G*_*j*_ (or an observation thereof *g*_*j*_) to be analyzed on either (i) the standardized scale or (ii) non standardized scale. To accommodate this, we introduce the variable 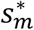, so that:

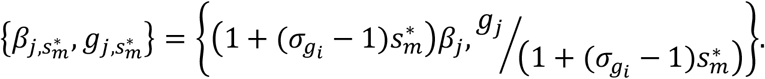

Hence, when 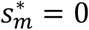 analyses are performed on the non-standardized scale and 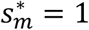 on the standardized scale. Default analyses assume the genetic effect is assessed on the non-standardized scale, i.e., 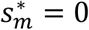. Note that, while p-values are generally invariant to the choice of effect scale 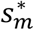, betas and standard errors are dependent on the specification of 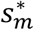.

### Sample-Level Newton-Raphson (SaLN-R) algorithm

Here we detail the isGWAS procedure for computing summary statistics 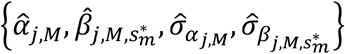 using only four data points,

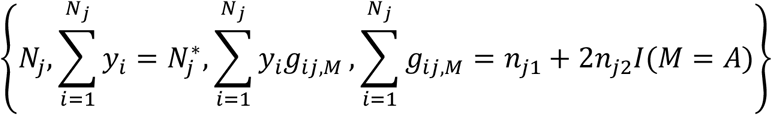

or, as we show, the quadruple

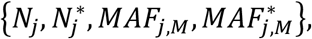

where *N*_*j*_ denotes the study or population sample size and 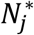 the number of cases in the sample, see ad-hoc estimator (Supplementary Information) for definitions of *n*_*j*·_. Note that we have allowed the sample size *N* and number of cases *N*^∗^ to vary by genotype *j*, this is useful when emulating results from GWAS. This is because the number of individuals analyzed in GWAS can vary by genotype owing to (e.g.,) quality of imputation or available data per variant and participant in a study. Ideally the sample size and number of cases would not vary by genotype and when using isGWAS to forecast GWAS results, users do not need not vary *N*_*j*_, i.e,.

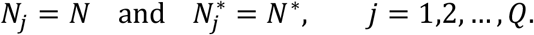

Given a vector of observed data 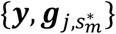, where 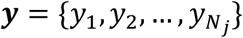 and 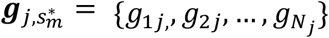, estimates of model parameters are typically derived by maximizing the log-likelihood function

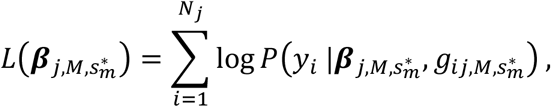

which is equivalent to identifying parameter values 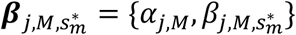 which satisfy:

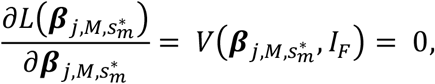

where 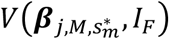 denotes the logistic score function, i.e.,

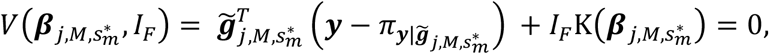

with *I*_*F*_ denoting an indicator function used to highlight that a Firth modified version of the score function has been used. For ease of mathematical presentation initially, we detail the Firth ad*j*usted SaLN-R algorithm later, i.e., we set *I*_*F*_ = 0 in this section. Additionally, to improve succinctness of notation, we drop the use of the parameter 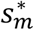 - reintroducing where necessary - and set *β*_*j,M*_= {*α*_*j,M*_, *β*_*j,M*_} and 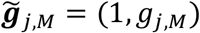 above, so that 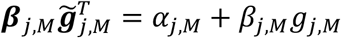. We compute candidate solutions to by expanding *V*(*β*_*j,M*_) as a Taylor series about a value *β*_*j*0,*M*_ and up to second order, i.e., using the Newton-Raphson (N-R) method:

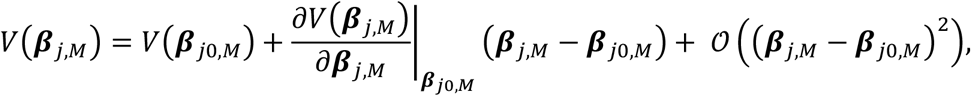

Which is re-written as

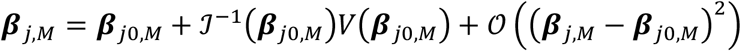

and generalized into an N-R iterative algorithm:

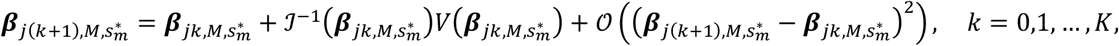

where we have re-introduced 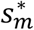 to highlight that the algorithm is dependent on the choice of effect scale. The variable 𝒥^−1^ denotes the inverse Fisher Information matrix, where

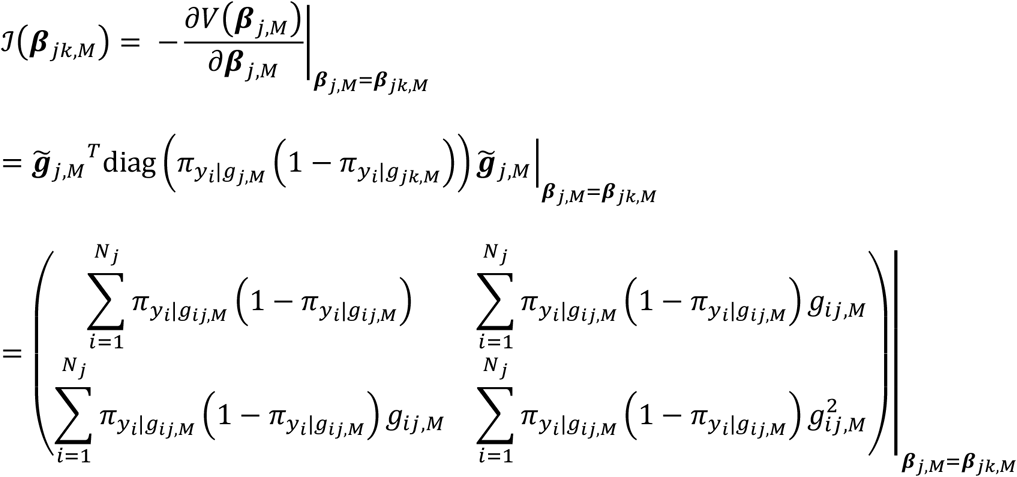

and the score function is given by

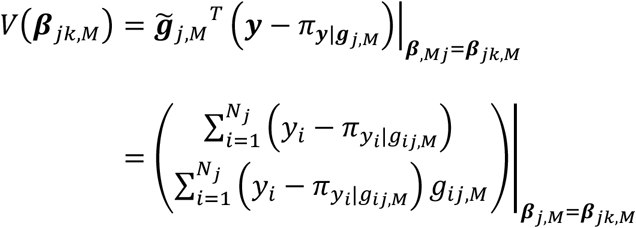

Both 𝒥(*β*_*jk,M*_) and *V*(*β*_*jk,M*_) above require individual-level data to compute their values. isGWAS aims to estimate values for these variables using sample-level information only, thereby avoiding the immediate need for individual data. To achieve this, we approximate both the Fisher Information matrix and the Score function via the pair {𝒥_𝕖_(*β*_*j*k,*M*_), *V*_𝕖_(*β*_*jk,M*_)}, where:

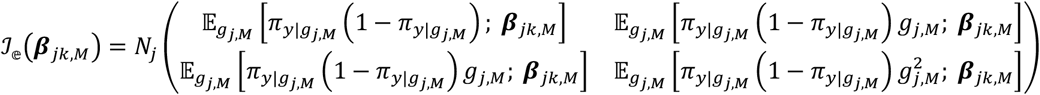

and

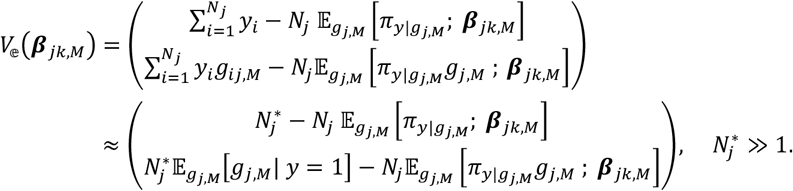

with 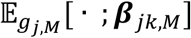 denoting that expectation is taken with respect to *g*_*j,M*_ and evaluated at *β*_*j,M*_= *β*_*jk,M*_. Note that {𝒥_𝕖_(*β*_*jk,M*_), *V*_𝕖_(*β*_*jk,M*_)} are motivated by switching from empirical, i.e., sample-based, estimates in {𝒥(*β*_*jk,M*_), *V*(*β*_*jk,M*_)} to their expected value analogues, which reverses the usual mode of estimation. Sample size *N*_*j*_ is presumed large and thus switching from sample-based to expected values in the N-R algorithm is well motivated. However, when the number of cases 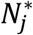 is ‘small’, an approximation of 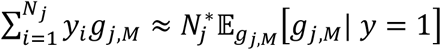 becomes weaker and we recommend using the statistic 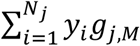. Values for the elements in {𝒥_𝕖_(*β*_*jk,M*_), *V*_𝕖_(*β*_*jk,M*_)} are computed via:

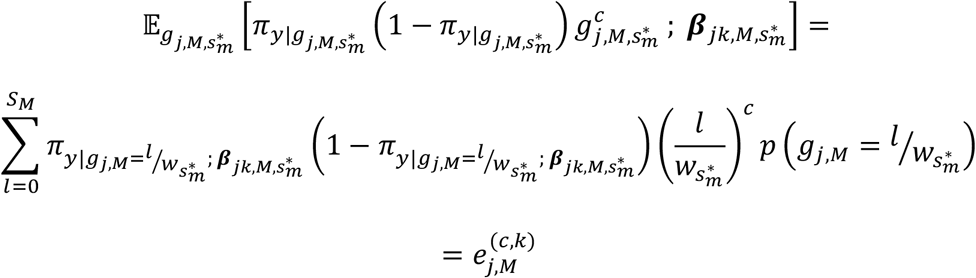

and

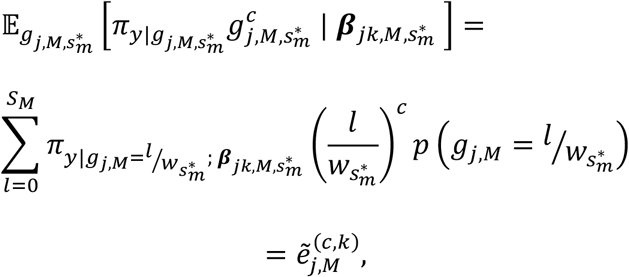

where 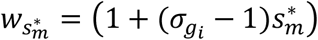 and the superscript and subscript in 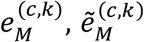 are used to highlight that expectation has been taken conditional on k-th iteration *β*_*jk,M*_ and under modelling assumption *M*(and implicitly effect scale 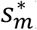). Probability mass 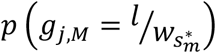 is either defined a-priori or can be approximated empirically, which we detail later. In combination, therefore, it follows that:

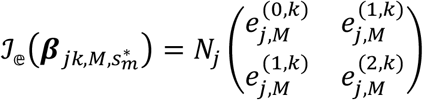

and

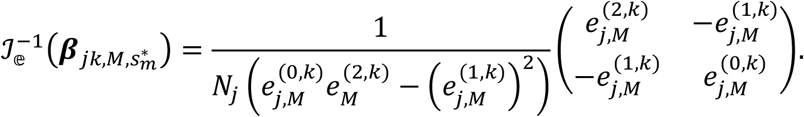

Following the same process that led to the above, we re-write the sample-level Score function 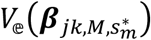 as:

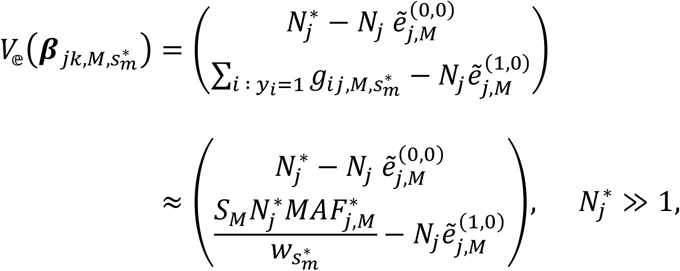

where we have used the following approximation:

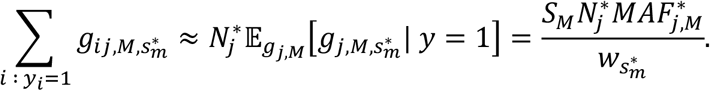

Hence, replacing the pair 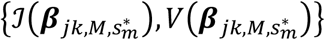 with the sample-level approximations 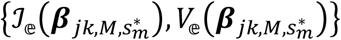 we furnish the SaLN-R algorithm:

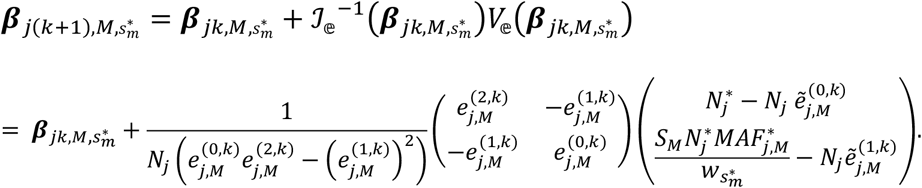

The standard error of the updates, 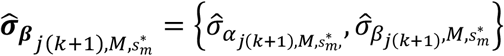, are given by the diagonal of the inverse Fisher information matrix, i.e.,

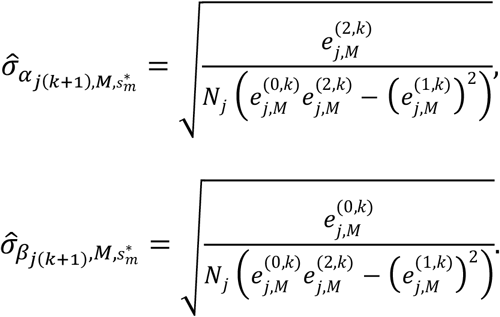

It can be shown from the above that:

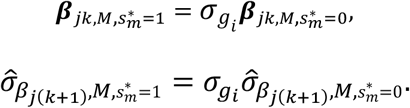

We set 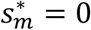 to compute values for the pair 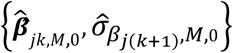 and use the above identities to return parameter estimates on the standardized scale 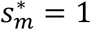. The data required to run the SaLN-R algorithm are:

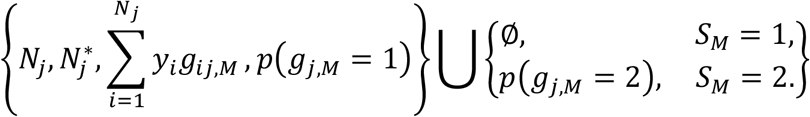

We use the approximations

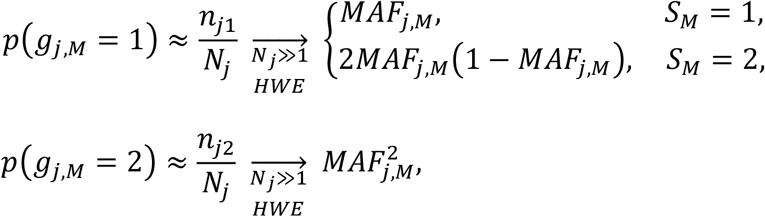

where 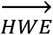 is used to denote under Hardy-Weinberg equilibrium.

The SaLN-R algorithm is extended to include Firth’s penalty function (see **Supplementary Information for more details**):

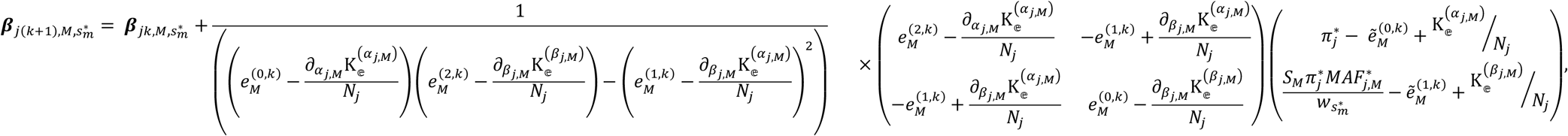

where

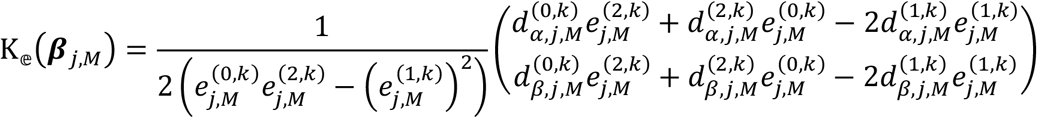

and

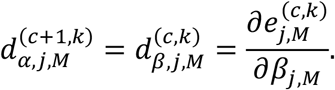

### isGWAS is computed using sufficient statistics

Under Hardy-Weinberg equilibrium, the quadruple 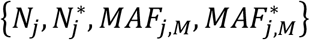 are combined to form the global and local (under a wide radius of convergence) sufficient statistics from the logistic model. Consequently, they hold all necessary information to compute regression parameter estimates 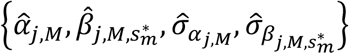 over a broad range of scenarios. Regardless of Hardy-Weinberg being valid or not, we show that the triple {*T*_1*j*_, *T*_2*j*_, *T*_3*j*_},

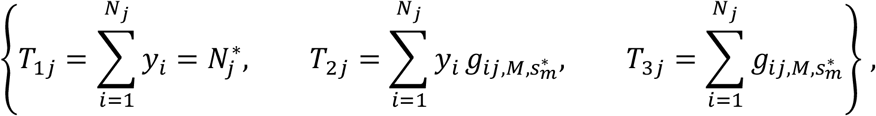

are the two global and one local sufficient statistics and these can alternatively be used as input variables in isGWAS. To show this, we write:

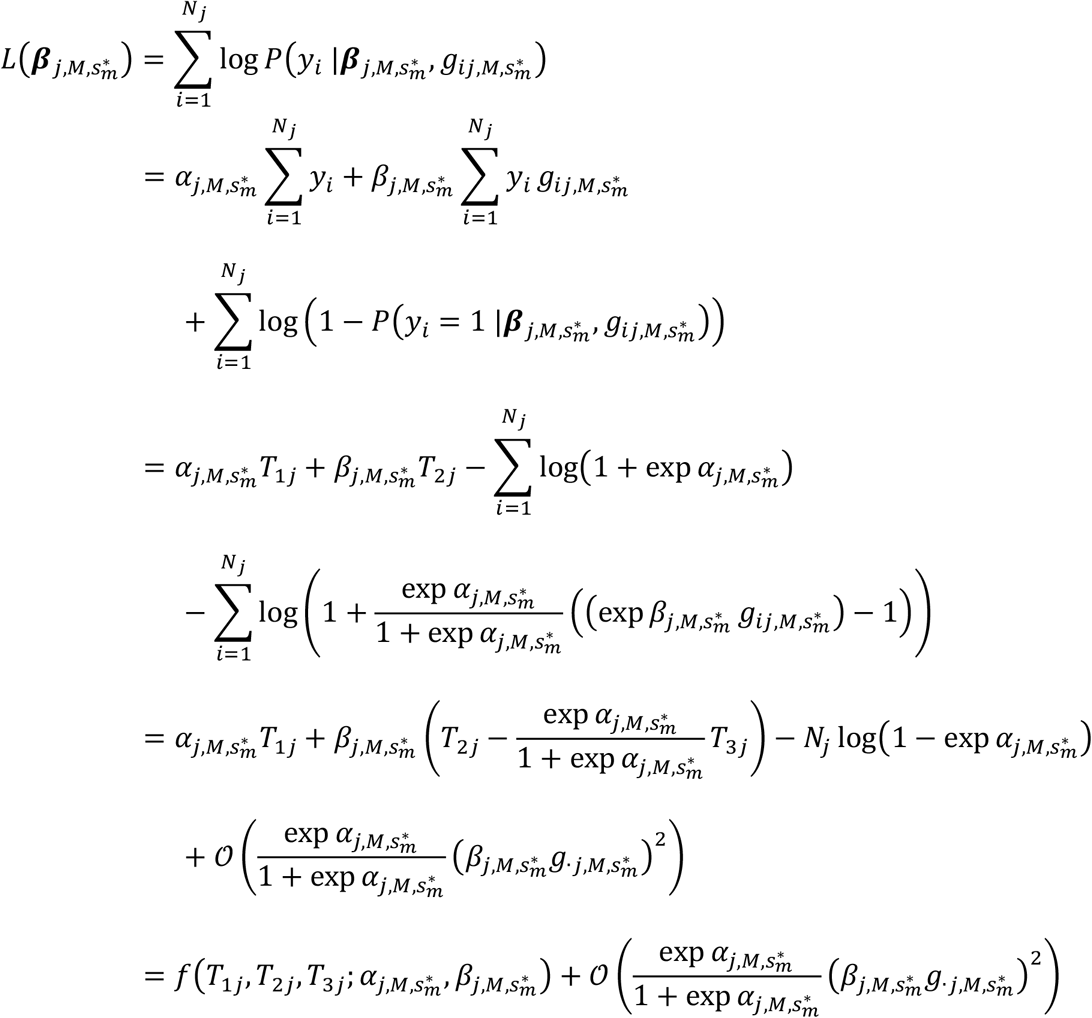

and valid when

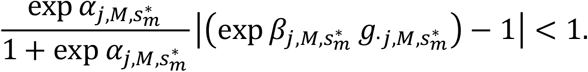

Hence, the global sufficient statistics are {*T*_1*j*_, *T*_2*j*_} and (on assuming random 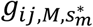 as in the SaLN-R algorithm) the locally sufficient statistic is {*T*_3*j*_}, where:

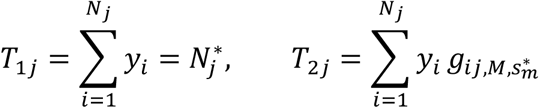

and

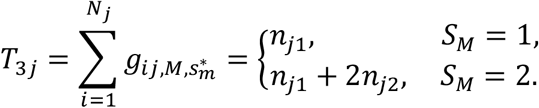

Under Hardy-Weinberg equilibrium, we can write

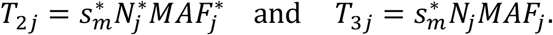

### Leapfrog re-sampler: forecasting results in target sample sizes

To estimate regression parameters 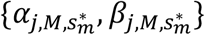 in larger target sample sizes, i.e., 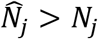, we propose the following strategy:

1. **Specify number** *K*, **sub-sample** *γ*_1_ **and target sample** *γ*_2_ **parameters**, where *K* ≥ 1, 0 < *γ*_1_ < 1 and *γ*_2_ > 1.
2. **Generate random sub-samples of individuals of size** 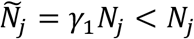. For each of *k* = 1,2, …, *K*, generate a random sub-sample 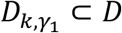, where 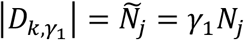.
3. **(Leapfrog-step) Compute subsample quadruple and pro*j*ect to target sample size** 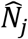. For each subsample 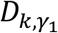, compute values 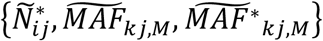 and pro*j*ect these on to the target sample size, i.e., 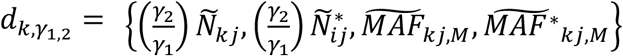 for sample 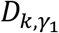
  - Note that 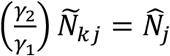, which is the target ‘future’ sample size.
4. **Deploy isGWAS across all (pro*j*ected) quadruples** 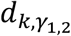 and record each estimate of the genetic effects, standard error and p-value 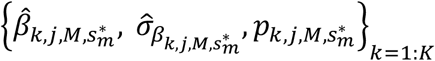
5. **Estimate p-value in target sample size** as a summary point estimate (e.g., median) or range across all *K* sub-samples,

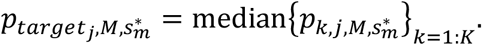

#### Data Quality Control: preparation of sufficient statistics for isGWAS

In order to deploy isGWAS successfully, the sufficient statistics are required to be prepared in a sample where only a single individual (preferably case) from pairs or n-tuples of 3^rd^, 2^nd^ and 1^st^ degree relatives is retained. Additionally, ethnical outliers must also be removed. In summary, to deploy isGWAS successfully we require either: (a) access to the sufficient statistics computed after duplications of related n-tuples and ethnical outliers are removed; or (b) access to the individual level data, whereupon the sufficient statistics can be prepared as described in (a). We provide a detailed outline of recommended Quality Control for genetic Individual Level Data (ILD) to running successfully isGWAS in **Supplementary Information**.

#### Application to Biobank data

The GWAS results used in the assessment of isGWAS were taken from large-scale analyses of UK Biobank[13], Biobank Japan[14] and the Psychiatric Genomics Consortium[15]. The UK Biobank[13] is a large-scale biomedical database and research resource containing in-depth genetic and health information from half a million UK participants. From the full available UK Biobank cohort, we obtain phenotypes for seven different diseases with varying levels of prevalence. These are Hypertension (IC10:I10), Asthma (IC10:J45), Atherosclerosis (IC10:I25), Glaucoma (IC10:H40), Stroke (IC10:I63), Colon Cancer (IC10:C18) and Thyroid Gland Cancer (IC10:C73) patients. From a total cohort of 502,422 participants, we used the following inclusion criteria: white British (Field 22006), non-related (>3rd degree), no patients with difference in reported (Field 31) and genetic (Field 22001) sex, no patients with aneuploidy (Field 22019), no patients with unusual heterozygosity and high missing rates (Field 22027). The ethnicity component is obtained from samples who self-identified as ‘White British’ according to Field 21000 and have very similar genetic ancestry based on a principal components analysis of the genotypes. Retaining one related individual (where we favour the retention of cases) we obtain a working sample size of ∼335,000 individuals; the approximate value is owing to small differences in the number of cases between disease phenotypes (**Supplementary Information**). Comparative analysis for these varying populations is reported in the main text. The prevalence ratios and exact number of cases and controls are provided in **Supplementary Table 1**. The variant based statistics needed for isGWAS were obtained from the imputed UK Biobank dataset. A quality info score>0.9 is applied to the data, and the number of cases and controls per variant and the MAF for variant in cases and controls is based on patients with non-missing genotypes for the variant using software PLINK[4]. Sample-level MAF>0.001 is used as inclusion criteria for the variants to analyse. For each disease, we run isGWAS analysis using default settings under the ‘additive’ genetic model. In addition, we also perform GWAS analysis using two-step REGENIE[8] applied to all variants with a MAF>0.001 and Genotype Score>0.99. Firth correction was enabled and performed on variants with p-value<0.1. REGENIE was also ad*j*usted for covariate information (age, sex, ancestry). For each disease we provide the following diagnostic plots: 1) mirrored Manhattan plot comparing directly p-values for isGWAS and REGENIE, 2) p-value – p-value plots comparing REGENIE and isGWAS, 3) *β* − *β* plots comparing REGENIE and isGWAS where we have colored the values by a) MAF and b) ratios of computed standard error (SE) between methods, i.e., 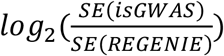. Across all diseases and variants considered, we compare performance of isGWAS and isGWAS-Firth to REGENIE-Firth.

Schizophrenia data from the Psychiatric Genomics Consortium[15] was used to conduct two different large-scale GWAS analysis. The first GWAS analysis was executed with data from 77,096 European individuals (33,640 cases, 43,456 controls)[25]. The second GWAS analysis was executed with data from the larger 130,644 European individuals (53,386 cases, 77,258 controls)[27]. We used the 2014 dataset to infer the 2022 results. To do this, we refine the significant results from both imputed 2014 and 2022 summary statistics using clumping with PLINK. The European 1000 Genomes Pro*j*ect v3[19] dataset was used as a reference population for the clumping procedure. Twelve strategies for clumping were explored: three were LD *R*^2^-based only, the other nine were a combination of clumping by LD block information and p-value thresholding. The refined variants are used to assess the inference capabilities of isGWAS both within each of the two datasets and the enrichment capabilities of isGWAS to infer p-values of the 2022 dataset using the 2014 dataset. For the 2014 dataset, 225 variants were remaining after the clumping. For the 2022 dataset, 451 variants were remaining after the clumping. From those, 54 are overlapping and 608 is the unique set between the two datasets.

The Biobank Japan data was used to conduct a large-scale GWAS with 212,453 Japanese individuals across 42 different diseases[24]. We obtained the published significantly associated loci (P < 5e-08) in autosomes from the GWAS findings which amounted to 309 variants across 30 different diseases. Similarly, we used the significantly associated X chromosome findings for males and females that amounted to a total of nine significantly associated loci across five diseases, although results are omitted from text. We applied isGWAS to the three different sets of variants using default parameters to assess the performance of isGWAS. To aid association interpretation, we use the following additional statistical tests to assess the accuracy and sensitivity of the isGWAS calculator for the Biobank Japan data. First, a classical ROC curve was produced where the true/false actual value was determined by various p-value thresholds (benchmarked against published Biobank Japan results). The isGWAS calculator is an inferential tool thus this usage of the ROC curve is unconventional, however, it provides us with the opportunity to assess the sensitivity to the choice of thresholds used to correct for multiple testing. These are 10^−10^, 10^−8^, 5 × 10^−8^, 10^−7^, where we have also used 9.58 × 10^−9^ for Biobank Japan as recommended by the authors[24]. AUC values were not obtained as this is not a standard classification problem and they are not interpretable in this context. Second, an adapted ROC curve was produced which accounts for two different thresholds – one more stringent one to determine the true positive rate and one less stringent one to determine the true negative rate. **Supplementary Figure 22** showcases this scenario and highlights the importance of a threshold choice and its impact on a sensitivity analysis. The main aim of isGWAS calculator is to be used as an inferential tool for truly significant or truly non-significant genetic signals. Thus, using two thresholds – one for truly significant and one for truly non-significant – provides us the assess the sensitivity of isGWAS to this scientific question. Third, the obtained *β* values were compared to the true ones by obtaining the percentage of 1) predicted *β* values in the 95% C.I.s of the true *β* values and 2) 95% C.I.s of the predicted *β* values in the 95% C.I.s of the true *β* values.

### Simulation scenarios

In the first scenario, for each individual *i* and iteration index *k*, we randomly generate disease status via *y*_*ik*_ ∼ *Ber* (π_*ik*_ *α*_*k*_, *β*) with probability of disease π_*ik*_ = *expit*(*α*_*k*_ + *βg*_*ik*_) and *g*_*ik*_ *Bin*(2, *MAF*_*k*_). Minor allele frequency is randomly selected from the set *MAF*_*k*_ ∈ {10^−4^, 5 × 10^−4^, 0.01} ⋃ { 0.025, 0.05, …, 0.5} and the genetic effect on disease risk is fixed as *β* = 0.5. In the second study, we allow the genetic effect to vary, i.e., *β* ≡ *β*_k_, by fixing disease status per individual and generating genotype data in controls *g*_*ik*_|*y*_*ik*_ = 0 ∼ *Bin*(2, *MAF*_*k*_) or cases 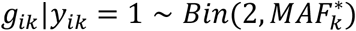, where minor allele frequency in cases is taken as the outer product with the sample minor allele frequency, with a random increase or decrease in frequency (which controls the magnitude and direction of genetic effect), i.e., we introduce the set 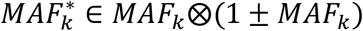. The parameter *β*_*k*_ is then estimated via each of the 5 estimators using the vector of simulated data {***y***_*k*_, ***g***_*k*_}.

We compare isGWAS and isGWAS-Firth against classical logistic and Firth corrected regression[16], [37], [38]. Details for the second scenario are provided alongside full description of the simulation protocol in the **Supplementary Information**.

#### Leapfrog re-sampler: simulation and real-data analyses

The parameters *{K, γ*_1_, *γ*_*2*_} in the leapfrog re-ampler are assessed over a variety of values. To attenuate the computational burden of a 3-dimensional grid search, we considered scenarios in which: K = 100, 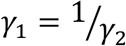 and a *γ*_*2*_-fold increase in sample size of *γ*_*2*_ ∈ {1.1,1.*2*5, 1.5, …, *2*.5}, i.e., a 10% to 150% increase in sample size. Furthermore, we take our working sample size to be 276,204 individuals, which matches the number of all unrelated individuals in our UKB sample (i.e., on not retaining any member of a related pair – which is therefore fixed between diseases). We used our simulation protocol (**Supplementary Information**) to generate synthetic samples and additionally assessed performance across all seven disease datasets in UK Biobank. Variants for assessment were selected after pruning in PLINK[4] was applied to the ∼11 million variants with the following parameters: genotype quality>0.99, MAF>0.01, HWE p < 10*e* − 15, 1000 bp windows, 100 variant increments, *R*^*2*^ > *0*.9. From the pruned variants, 5% were selected uniformly from variants with p > 10^−6^ and all variants with p *≤* 10−^6^ were retained. Final number of variants progressed for LRS for the seven diseases are provided in **Supplementary Table 10**. For simulated data, data for smaller sub-samples were simulated using full cohort and empirical distributions for MAF and disease prevalence. In our tests of the LRS, we assess the predictive properties of isGWAS on real-life data where the ground truth is either computed from the entire sample or provided in the literature. Predictions from X-fold increases in sample size are compared using standard accuracy, FDR, FPR and TPR measures based on a putative true significance threshold of *5e* − *08*.

### Computational resources

Real-life analyses were performed using up to 48 virtual CPU cores of a 2.5 GHz Intel Xeon Gold 6240R processor with 64 GB of memory. Simulation analyses were performed using up to 8 virtual CPU cores of a 2.4 GHz Intel Core i9 processor.

#### Computational comparison protocol

We contrast the computational performance of isGWAS and REGENIE (Step-2 only). For clarity, REGENIE Step-1 simplifies the outcome and model by projecting out covariate information, before variant-disease association analyses are performed in Step-2. To directly compare both methods, we performed individual GWA analyses of each of the seven diseases considered in UK Biobank across ∼11m variants for ∼335,000 individuals. Owing to computational cost of the ILD method, we summarise results from a single GWA analysis per trait. Performance of isGWAS across repeated runs, for varying numbers of SNPs and available CPUs, up to a maximum of 10m variants, is also performed.

## Data availability

The genotype data, phenotype status and allele counts were extracted from UK Biobank[13] to support the findings of this study. The genome-wide association summary data with available allele frequencies and cohort counts that was used to support the findings of this study are available from: Psychiatric Genomics Consortium[15] and Biobank Japan[14].

## Code availability

The tool is available for use on the webportal www.optima-isgwas.com. The isGWAS algorithm is also available on github (https://github.com/cnfoley/isgwas/).

## Acknowledgements

The authors would like to thank Prof Peter Visscher for his insightful comments and suggestions on the methodology. The authors are additionally grateful to Manju Dissanayake, Chris Marshall, Michael Mulholland, Andrew Donald and the Optima Partners Data Science and Engineering software development team.

## Author contributions

CNF developed the mathematical and statistical methodologies, developed the statistical software and conceptualized the design. ZK developed the statistical software and webtool, designed the methodological analysis pipeline and conducted the real-life analyses. REM contributed to the interpretation of results. HR conceptualized and supervised the study. BBS conceptualized, designed the study, contributed to the method, application, and contextualization of the study. All authors contributed to the writing of the manuscript.

