## Supplementary Information and Supplementary Figures 1-22 for "isGWAS: ultra-high-throughput, scalable and equitable inference of genetic associations with disease"

#### 1. The isGWAS Method: Firth adjustment

##### 1.1 Firth modified SaLN-R algorithm

Firth[1] modified the score function $V(\boldsymbol{\beta}_{j,M})$ using Jeffrey's prior as a bias reducing penalty function, removing the leading order asymptotic bias of the maximum likelihood estimate of the parameters $\boldsymbol{\beta}_{j,M}$. The objective is to find values $\boldsymbol{\beta}_{j,M}$ that satisfy the modified score function $V_F(\boldsymbol{\beta}_{j,M})$:

$$V_F(\boldsymbol{\beta}_{j,M}) = \tilde{\boldsymbol{g}}_{j,M}^T \left(\boldsymbol{y} - \pi_{\boldsymbol{y}|\tilde{\boldsymbol{g}}_{j,M}}\right) + \mathrm{K}(\boldsymbol{\beta}_{j,M}) = 0$$

where

$$\mathrm{K}(\boldsymbol{\beta}_{j,M}) = \begin{pmatrix} \mathrm{K}_{\mathbb{e}}^{(\alpha_{j,M})} \\ \mathrm{K}_{\mathbb{e}}^{(\beta_{j,M})} \end{pmatrix}.$$

Following the process that led to the standard isGWAS algorithm, we set

$$\mathrm{K}_{\mathbb{e}}^{(\alpha_{j,M})} = \frac{1}{2} tr\left[ I_{\mathbb{e}}^{-1}(\boldsymbol{\beta}_{j,M}) \frac{\partial I_{\mathbb{e}}(\boldsymbol{\beta}_{j,M})}{\partial \alpha_{j,M}} \right]$$

$$\mathrm{K}_{\mathbb{e}}^{(\beta_{j,M})} = \frac{1}{2} tr\left[ I_{\mathbb{e}}^{-1}(\boldsymbol{\beta}_{j,M}) \frac{\partial I_{\mathbb{e}}(\boldsymbol{\beta}_{j,M})}{\partial \beta_{j,M}} \right]$$

which can be written as:

$$\mathrm{K}_{\mathbb{e}}(\boldsymbol{\beta}_{j,M}) = \frac{1}{2\left(e_{j,M}^{(0,k)} e_{j,M}^{(2,k)} - \left(e_{j,M}^{(1,k)}\right)^2\right)} \begin{pmatrix} d_{\alpha,j,M}^{(0,k)} e_{j,M}^{(2,k)} + d_{\alpha,j,M}^{(2,k)} e_{j,M}^{(0,k)} - 2 d_{\alpha,j,M}^{(1,k)} e_{j,M}^{(1,k)} \\ d_{\beta,j,M}^{(0,k)} e_{j,M}^{(2,k)} + d_{\beta,j,M}^{(2,k)} e_{j,M}^{(0,k)} - 2 d_{\beta,j,M}^{(1,k)} e_{j,M}^{(1,k)} \end{pmatrix}$$

with

$$d_{\alpha,j,M}^{(c+1,k)} = d_{\beta,j,M}^{(c,k)} = \frac{\partial e_{j,M}^{(c,k)}}{\partial \beta_{j,M}}$$

$$= \frac{\partial \mathbb{E}_{g_{j,M,s_m^*}}\left[ \pi_{y|g_{j,M,s_m^*}} \left(1 - \pi_{y|g_{j,M,s_m^*}}\right) g_{j,M,s_m^*}^c \,;\, \boldsymbol{\beta}_{jk,M,s_m^*} \right]}{\partial \beta_{j,M,s_m^*}}$$

$$= \sum_{l=0}^{s_M} \pi_{y|g_{j,M}=l/w_{s_m^*};\boldsymbol{\beta}_{jk,M,s_m^*}} \left(1 - \pi_{y|g_{j,M}=l/w_{s_m^*};\boldsymbol{\beta}_{jk,M,s_m^*}}\right)$$

$$\times \left(1 - 2\pi_{y|g_{j,M}=l/w_{s_m^*};\boldsymbol{\beta}_{jk,M,s_m^*}}\right) \left(l/w_{s_m^*}\right)^{c+1} p\left(g_{j,M} = l/w_{s_m^*}\right).$$

Following the same approach, but now for the Fisher information matrix:

$$\mathcal{I}\left(\boldsymbol{\beta}_{jk,M}, I_F = 1\right) = -\frac{dV_F\left(\boldsymbol{\beta}_{j,M}\right)}{d\boldsymbol{\beta}_{j,M}}$$

$$= \widetilde{\boldsymbol{g}}_{j,M}{}^T \operatorname{diag}\left(\pi_{y_i|g_{j,M}}\left(1 - \pi_{y_i|g_{jk,M}}\right)\right) \widetilde{\boldsymbol{g}}_{j,M} - \frac{d\mathrm{K}\left(\boldsymbol{\beta}_{j,M}\right)}{d\boldsymbol{\beta}_{j,M}},$$

it follows that the Firth modified SaLN-R algorithm is given by

$$\boldsymbol{\beta}_{j(k+1),M,s_m^*} = \boldsymbol{\beta}_{jk,M,s_m^*} + \frac{1}{\left(\left(e_M^{(0,k)} - \frac{\partial_{\alpha_{j,M}}\mathrm{K}_{\mathbb{e}}^{(\alpha_{j,M})}}{N_j}\right)\left(e_M^{(2,k)} - \frac{\partial_{\beta_{j,M}}\mathrm{K}_{\mathbb{e}}^{(\beta_{j,M})}}{N_j}\right) - \left(e_M^{(1,k)} - \frac{\partial_{\beta_{j,M}}\mathrm{K}_{\mathbb{e}}^{(\alpha_{j,M})}}{N_j}\right)^2\right)}$$

$$\times \begin{pmatrix} e_M^{(2,k)} - \frac{\partial_{\alpha_{j,M}}\mathrm{K}_{\mathbb{e}}^{(\alpha_{j,M})}}{N_j} & -e_M^{(1,k)} + \frac{\partial_{\beta_{j,M}}\mathrm{K}_{\mathbb{e}}^{(\alpha_{j,M})}}{N_j} \\ -e_M^{(1,k)} + \frac{\partial_{\beta_{j,M}}\mathrm{K}_{\mathbb{e}}^{(\alpha_{j,M})}}{N_j} & e_M^{(0,k)} - \frac{\partial_{\beta_{j,M}}\mathrm{K}_{\mathbb{e}}^{(\beta_{j,M})}}{N_j} \end{pmatrix} \begin{pmatrix} \pi_j^* - \tilde{e}_M^{(0,k)} + \mathrm{K}_{\mathbb{e}}^{(\alpha_{j,M})}/N_j \\ \frac{S_M \pi_j^* MAF_{j,M}^*}{w_{s_m^*}} - \tilde{e}_M^{(1,k)} + \mathrm{K}_{\mathbb{e}}^{(\beta_{j,M})}/N_j \end{pmatrix}$$

##### *1.1.1 SaLN-R: without sample-size*

The SaLN-R algorithm can be re-written to avoid the need for a user to specify a sample size $N_j$, i.e.,

$$\boldsymbol{\beta}_{j(k+1),M} = \boldsymbol{\beta}_{jk,M} + \frac{1}{N_j\left(e_M^{(0,k)} e_M^{(2,k)} - \left(e_M^{(1,k)}\right)^2\right)} \begin{pmatrix} e_M^{(2,k)} & -e_M^{(1,k)} \\ -e_M^{(1,k)} & e_M^{(0,k)} \end{pmatrix} \begin{pmatrix} N_j^* - N_j \tilde{e}_M^{(0,k)} \\ S_M N_j^*\left(MAF_{j,M}^* - MAF_{j,M}\right) - N_j \tilde{e}_M^{(1,k)} \end{pmatrix}$$

$$= \boldsymbol{\beta}_{jk,M} + \frac{1}{\left(e_M^{(0,k)} e_M^{(2,k)} - \left(e_M^{(1,k)}\right)^2\right)} \begin{pmatrix} e_M^{(2,k)} & -e_M^{(1,k)} \\ -e_M^{(1,k)} & e_M^{(0,k)} \end{pmatrix} \begin{pmatrix} \pi_j^* - \tilde{e}_M^{(0,k)} \\ \frac{S_M \pi_j^* MAF_{j,M}^*}{w_{s_m^*}} - \tilde{e}_M^{(1,k)} \end{pmatrix}$$

where we have let

$$\pi_j^* = \frac{N_j^*}{N_j}.$$

In this 'sample-size free' use of SaLN-R, users need only provide three variables:

$$\{\pi_j^*, MAF_{j,M}, MAF_{j,M}^*\}$$

The above result can be motivated (computed) following the approach that led to (eqn#) but now in the pursuit of finding solutions to the population score function, $V_{pop}$:

$$V_{pop}(\boldsymbol{\beta}_{j,M}) = \begin{pmatrix} Y - \pi_{Y|g_{j,M},\boldsymbol{\beta}_{j,M}} \\ \left(Y - \pi_{Y|g_{j,M},\boldsymbol{\beta}_{j,M}}\right) g_{j,M} \end{pmatrix} = \mathbf{0}.$$

Thus, the parameter $\pi_j^*$ can be considered the population disease prevalence. In this set-up, there are no standard errors for the resulting main effect parameters $\boldsymbol{\beta}_{j(k+1),M}$ and naturally a sample-based Firth adjustment is not appropriate.

1.2 SaLN-R Likelihood Ratio Test

Under the null of no effect of genotype on disease, estimates of the parameter $\alpha_{j,M}$ are obtained by solving:

$$\hat{\alpha}_{j,M|\beta_{j,M}=0} : \sum_{i=1}^{N_j} y_i - \pi_i = 0.$$

This can be optimized explicitly, i.e.,

$$N_j^* - N_j \frac{e^{\hat{\alpha}_{j,M|\beta_{j,M}=0}}}{1 + e^{\hat{\alpha}_{j,M|\beta_{j,M}=0}}} = 0$$

$$\hat{\alpha}^{(0)}{}_{j,M} = \hat{\alpha}_{j,M|\beta_{j,M}=0} = \log\left|\frac{N_j^*}{N_j - N_j^*}\right|.$$

Conditional on $\hat{\alpha}^{(0)}{}_{j,M}$, the log-likelihood under the null hypothesis is given by

$$L(\hat{\alpha}^{(0)}{}_{j,M}, 0) = \sum_{i=1}^{N_j} \log P(y_i \,|\hat{\alpha}^{(0)}{}_{j,M}, g_{ij,M})$$

$$= \sum_{i=1}^{N_j} \hat{\alpha}^{(0)}{}_{j,M} y_i + \ln\left(1 - \pi\left(\tilde{g}_{ij} | \hat{\alpha}^{(0)}{}_{j,M}\right)\right)$$

$$= \hat{\alpha}^{(0)}{}_{j,M} N_j^* + N_j \log\left(1 - \pi_{y|\,\hat{\alpha}^{(0)}{}_{j,M}}\right).$$

The log-likelihood under the alternative hypothesis is written:

$$L\left(\hat{\alpha}_{j,M}, \hat{\beta}_{j,M,s_m^*}\right) = \sum_{i=1}^{N_j} \log P\left(y_i \,|\hat{\beta}_{j,M,s_m^*}, g_{ij,M,s_m^*}\right)$$

$$= \sum_{i=1}^{N_j} y_i\left(\hat{\alpha}_{j,M} + \hat{\beta}_{j,M,s_m^*} \tilde{g}_{ij,s_m^*}\right) + \ln\left(1 - \pi\left(\tilde{g}_{ij,s_m^*} | \hat{\beta}_{j,M,s_m^*}\right)\right)$$

$$\approx \left(\hat{\alpha}_{j,M} + \frac{\hat{\beta}_{j,M,s_m^*} S_M MAF_{j,M}^*}{w_{s_m^*}}\right) N_j^*$$

$$+ N_j \sum_{l=0}^{S_M} \ln\left(1 - \pi_{y|g_{j,M}={}^{l}/_{w_{s_m^*}};\, \hat{\beta}_{jk,M,s_m^*}}\right) p\left(g_{j,M} = {}^{l}/_{w_{s_m^*}}\right).$$

Thus, writing the likelihood-ratio (LTR) as

$$2\left(L\left(\hat{\alpha}_{j,M}, \hat{\beta}_{j,M,s_m^*}\right) - L\left(\hat{\alpha}_{j,M}, 0\right)\right)$$

the SaLN-R LTR statistic is given by

$$\chi^2_{LRT_{isGWAS}} = 2\left(N_j^*\left(\hat{\alpha}_{j,M} - \hat{\alpha}^{(0)}{}_{j,M} + \frac{\hat{\beta}_{j,M,s_m^*} S_M MAF_{j,M}^*}{w_{s_m^*}}\right) + \right.$$

$$\left. N_j\left(\left(\sum_{l=0}^{S_M} \log\left(1 - \pi_{y|g_{j,M}={}^{l}/_{w_{s_m^*}};\, \hat{\beta}_{jk,M,s_m^*}}\right) p\left(g_{j,M} = {}^{l}/_{w_{s_m^*}}\right)\right) - \log\left(1 - \pi_{y|\,\hat{\alpha}^{(0)}{}_{j,M}}\right)\right)\right),$$

which we assess the probability of under $\chi_1^2$ distribution.

*1.2.1 Robust standard errors: sandwich estimation*

The so-called sandwich estimator is given by:

$$\operatorname{diag}\left[\left(\mathcal{J}_{\mathrm{F}}^{-1}\left(\boldsymbol{\beta}_{jk,M,s_m^*}\right)\left(\sum_{i=1}^{N_j} V_i\left(\boldsymbol{\beta}_{j,M}\right)\left(V_i\left(\boldsymbol{\beta}_{j,M}\right)\right)^T\right)\mathcal{J}_{\mathrm{F}}^{-1}\left(\boldsymbol{\beta}_{jk,M,s_m^*}\right)\right)^{-\frac{1}{2}}\right]$$

where $\mathcal{J}_{.}^{-1}\left(\boldsymbol{\beta}_{jk,M,s_m^*}\right)$ was defined in section 1.1 and

$$V_i\left(\boldsymbol{\beta}_{j,M}\right)\left(V_i\left(\boldsymbol{\beta}_{j,M}\right)\right)^T = \begin{pmatrix} v_{i11} & v_{i12} \\ v_{i21} & v_{i22} \end{pmatrix}$$

with

$$v_{i11} = (y_i - \pi_i)^2 + 2\mathrm{K}_{\mathbb{e},i}^{(\alpha_{j,M})}(y_i - \pi_i) + \left(\mathrm{K}_{\mathbb{e},i}^{(\alpha_{j,M})}\right)^2,$$

$$v_{i12} = v_{i21} = (y_i - \pi_i)^2 g_{ij,M} + \left(\mathrm{K}_{\mathbb{e},i}^{(\beta_{j,M})} + \mathrm{K}_{\mathbb{e},i}^{(\alpha_{j,M})} g_{ij,M}\right)(y_i - \pi_i) + \mathrm{K}_{\mathbb{e},i}^{(\beta_{j,M})}\mathrm{K}_{\mathbb{e},i}^{(\alpha_{j,M})},$$

$$v_{22} = (y_i - \pi_i)^2 g_{ij,M}^2 + 2\mathrm{K}_{\mathbb{e},i}^{(\beta_{j,M})}(y_i - \pi_i) g_{ij,M} + \left(\mathrm{K}_{\mathbb{e},i}^{(\beta_{j,M})}\right)^2.$$

We compute estimates for $\{v_{i11}, v_{i12}, v_{i22}\}$ following the same approach that led to (#eqn).

#### 1.3 Converting between Additive and Recessive or Dominant models

The probabilities for each possible combination of reference and effect allele in $e_M^{(c,k)}$ and $\tilde{e}_M^{(c,k)}$ under the additive model are

$$p\left(g_j = l - S_M MAF_{j,M} \mid M = A\right) = \begin{cases} \left(1 - MAF_{j,A}\right)^2, & l = 0, \\ 2MAF_{j,A}\left(1 - MAF_{j,A}\right), & l = 1, \\ MAF_{j,A}^2, & l = 2, \end{cases}$$

while under dominant or recessive models,

$$p\left(g_j = l - S_M MAF_{j,M} \mid M \in \{D,R\}\right) \begin{cases} 1 - MAF_{j,M} & l = 0, M \in \{D,R\}, \\ MAF_{j,M} & l = 1, M \in \{D,R\}. \end{cases}$$

Taking the reference model to be $M = A$, i.e., the additive model, we can translate minor allele frequencies between model choices by matching the identities above:

$$MAF_{j,D} = MAF_{j,A}\left(2 - MAF_{j,A}\right),$$

$$MAF_{j,R} = MAF_{j,A}^2.$$

Thus, a user need only supply the readily available information $MAF_{j,A}$, and analogously $MAF_{j,A}^*$, and isGWAS computes $\left\{\hat{\alpha}_{j,M}, \hat{\beta}_{j,M}, \hat{\sigma}_{\alpha_{j,M}}, \hat{\sigma}_{\beta_{j,M}}\right\}$ under alternative model assumptions $M \in \{D, R\}$.

#### 1.4 Setting initial values for the SaLN-R algorithm

To improve computational efficiency and stability, initial values $\boldsymbol{\beta}_{j0,M} = \{\alpha_{j0,M}, \beta_{j0.M}\}$ for the SaLN-R algorithm are deduced by writing the probability of disease $Y$ conditional on genotype $g_j$ and model $M$, as a power series in $\beta_{j,M} g_{j,M}$ and retaining leading order terms. That is,

$$\pi_{Y|g_{j},M} = P(Y = 1 \mid g_j, M) = \frac{e^{(\alpha_{j,M} + \beta_{j,M} g_{j,M})}}{1 + e^{(\alpha_{j,M} + \beta_{j,M} g_{j,M})}}$$

$$= \frac{e^{\alpha_{j,M}}}{(1 + e^{\alpha_{j,M}})} \frac{e^{\beta_{j,M} g_{j,M}}}{1 + \frac{e^{\alpha_{j,M}}}{(1 + e^{\alpha_{j,M}})}\left(e^{\beta_{j,M} g_{j,M}} - 1\right)}$$

$$= \frac{e^{\alpha_{j,M}}}{(1 + e^{\alpha_{j,M}})} + \left(\frac{e^{\alpha_{j,M}}}{(1 + e^{\alpha_{j,M}})^2}\right)\beta_{j,M} g_{j,M} + \left(\frac{e^{\alpha_{j,M}}(1 - e^{\alpha_{j,M}})}{2(1 + e^{\alpha_{j,M}})^3}\right)\beta_{j,M}^2 g_{j,M}^2 + \cdots$$

$$a_{j,M} + b_{j,M} g_{j,M} + \mathcal{O}\left(\beta_j^2 g_j^2\right), \qquad \frac{e^{\alpha_{j,M}}}{(1 + e^{\alpha_{j,M}})}\left|e^{\beta_{j,M} g_{j,M}} - 1\right| < 1,$$

where

$$\alpha_{j,M} = \log\left(\frac{a_{j,M}}{1 - a_{j,M}}\right) \quad \text{and} \quad \beta_{j,M} = \frac{b_{j,M}}{a_{j,M}(1 - a_{j,M})}.$$

Following the approach of[2], estimates $\{\hat{a}_{j,M}, \hat{b}_{j,M}\}$ can be derived by solving the linearized approximation to the Score function, and re-writing the solution in terms of the four sample-level values $\{N_j, N_j^*, MAF_{j,M}, MAF_{j,M}^*\}$, i.e.,

$$\hat{a}_{j,M} = \frac{\sum_{i=1}^{N_j} y_{ij}}{N_j} = \frac{N_j^*}{N_j},$$

$$\hat{b}_{j,M} = \frac{MAF_{j,M}^{*} - MAF_{j,M}}{MAF_{j,M}\left(1 - MAF_{j,M}\right)} \frac{N_j^{*}}{\left(N_j - 1\right)} \quad : \quad \frac{e^{\alpha_{j,M}}}{\left(1 + e^{\alpha_{j,M}}\right)} \left| e^{\beta_{j,M} g_{j,M}} - 1 \right| < 1 \text{ and } N_j \gg 1.$$

Hence, we take as initial values $\boldsymbol{\beta}_{j0,M} = \{\alpha_{j0,M}, \beta_{j0.M}\}$ in the SaLN-R algorithm:

$$\alpha_{j0,M} = \log\left(\frac{N_j^{*}}{N_j - N_j^{*}}\right) \quad \text{and} \quad \beta_{j0,M} = \frac{MAF_{j,M}^{*} - MAF_{j,M}}{MAF_{j,M}\left(1 - MAF_{j,M}\right)} \frac{N_j}{\left(N_j - N_j^{*}\right)} \left(1 + O\left(N_j^{-1}\right)\right).$$

$$\beta_{j0,M} = \frac{\left(\frac{MAF_{j,M}^{*}}{MAF_{j,M}} - 1\right)}{\left(1 - MAF_{j,M}\right)} \frac{N_j}{\left(N_j - N_j^{*}\right)} \approx \frac{\left(\frac{MAF_{j,M}^{*}}{MAF_{j,M}} - 1\right)}{\left(1 - \pi_M\right)}, \qquad MAF_{j,M} \ll 0.5.$$

We also allow the option of initializing the isGWAS algorithm using the ad-hoc estimator[3] which, when valid (typically for common variants and disease prevalence > 0.01), was shown to increase time to convergence.

#### 1.5 Convergence diagnostic for initialization isGWAS

The initial values $\boldsymbol{\beta}_{j0,M} = \{\alpha_{j0,M}, \beta_{j0.M}\}$ in (eqn#) are sensible when

$$\frac{e^{\alpha_{j0,M}}}{\left(1 + e^{\alpha_{j0,M}}\right)} \left| e^{\beta_{j0,M} g_{j,M}} - 1 \right| < 1.$$

During extensive testing, we observed that the proposed isGWAS initial values occasionally violate the inequality above and the SaLN-R algorithm can either: (a) fail to converge; or (b) diverge. Further investigation of these scenarios revealed that scenarios (a) and (b) arose due to a poor choice in the initial specification of the log-odds $\beta_{j0.M}$ - differences between initial and final estimates of the intercept in the linear predictor were typically very small, i.e., $\left|\alpha_{j0,M} - \alpha_{jk,M}\right| \ll 1$. To help overcome potential initialization issues, we developed the following heuristic, which repeatedly resets the initial values $\{\alpha_{j0,M}, \beta_{j0.M}\}$ until SaLN-R converges or reaches a user defined timeout. Let

$$R\left(\boldsymbol{\beta}_{j0,M}, g_j\right) = \frac{e^{\alpha_{j0,M}}}{\left(1 + e^{\alpha_{j0,M}}\right)} \left| e^{\beta_{j0,M} g_{j,M}} - 1 \right|.$$

isGWAS deploys the follow (re)initialization procedure:

**Step 1:** initialize the SaLN-R algorithm

$$\alpha_{j0,M} = \log\left(\frac{N_j^*}{N_j - N_j^*}\right),$$

$$\beta_{j0,M} = \begin{cases} \dfrac{MAF_{j,M}^* - MAF_{j,M}}{MAF_{j,M}\left(1 - MAF_{j,M}\right)} \dfrac{N_j}{\left(N_j - N_j^*\right)}, & R\left(\boldsymbol{\beta}_{j0,M}, g_j = 1\right) \leq 1, \\ \ln\left(2 + e^{-\hat{a}_{i0}}\right), & R\left(\boldsymbol{\beta}_{j0,M}, g_j = 1\right) > 1. \end{cases}$$

**Step 2**: deploy the 'Repeated initialization algorithm' below:

**Algorithm II:** Repeated initialization algorithm

**Require:** initial values $\{\alpha_{j0,M}, \beta_{j0,M}\}$, data $\{N_j, N_j^*, MAF_{j,M}, MAF_{j,M}^*\}$, error tolerance $\epsilon_{max}$, step-error $\Delta\epsilon_0 > 0$ and max iterations $\tau > 0$.

    **if** $\beta_{j0,M} > 0$ and $\beta_{j0,M} > \ln(2 + e^{-\alpha_{j0,M}})$ **then**

    $\beta_{j0,M} = \ln(2 + e^{-\alpha_{j0,M}})$

    **end if**

**Set** $i = 0, k = 0$

**While** $k \leq \tau$ and $\Delta\epsilon_0 > \epsilon_{max}$

    **Update** $\{\alpha_{j(k+1),M}, \beta_{j(k+1),M}\} = \text{SaLNR}(\alpha_{jk,M}, \beta_{jk,M})$

        **if** $\{\alpha_{j(k+1),M}, \beta_{j(k+1),M}\} \in \{\text{NaN}, \pm\infty\}$ **then**

        $i \rightarrow i + 1$

        $\{\alpha_{j(k+1),M}, \beta_{j(k+1),M}\} = \left\{\frac{\alpha_{j0,M}}{2^i}, \frac{\ln(2+e^{-\alpha_{j0,M}})}{2^i}\right\}$

        **end if**

    $\Delta\epsilon_0 = \max\{|\alpha_{j(k+1),M} - \alpha_{jk,M}|, |\beta_{j(k+1),M} - \beta_{jk,M}|\}$

    $k \rightarrow k + 1$

**End while**

**Return** $\{\alpha_{j(k+1),M}, \beta_{j(k+1),M}\}$

#### 1.6 Automatic data quality check: successfully running isGWAS

For the $j$th variant, the sample-based estimate of the minor allele frequency ($MAF_{j,M}$) satisfies the following relationship:

$$MAF_{j,M} = \frac{N_j^*}{N_j} MAF_{j,M}^* + \left(1 - \frac{N_j^*}{N_j}\right) MAF_{j,M_{control}},$$

where $MAF_{j,M} \geq 0,\ MAF_{j,M}^* \geq 0,\ N_j \geq N_j^* > 0$ and $MAF_{j,M_{control}}$ denotes the minor allele frequency in the control-group. When deploying isGWAS, users should make sure that their choice of input variables,

$$\{N_j, N_j^*, MAF_{j,M}, MAF_{j,M}^*\}$$

return a valid value for $MAF_{j,M_{control}}$, which is equivalent to satisfying the following inequality:

$$MAF_{j,M_{control}} = \left(\frac{N_j}{N_j - N_j^*}\right)\left(MAF_{j,M} - \frac{N_j^*}{N_j} MAF_{j,M}^*\right) \geq 0$$

$$\Rightarrow \left(MAF_{j,M} - \frac{N_j^*}{N_j} MAF_{j,M}^*\right) \geq 0.$$

To avoid issues of numerical instability, i.e., when values of $MAF_{j,M_{control}}$ approach numerical internal precision boundaries, and to improve overall numerical stability of the SaLN-R algorithm, the isGWAS software automatically detects and avoid scenarios in which:

$$\left(\frac{N_j}{N_j - N_j^*}\right)\left(MAF_{j,M} - \frac{N_j^*}{N_j} MAF_{j,M}^*\right) = MAF_{j,M_{control}} < \epsilon_{fp},$$

where $\epsilon_{fp}$ is taken to be $\epsilon_{fp} = 10^{-10}$. This straightforward numerical check demonstrably improved results of SaLN-R vs classical individual-level data regression analyses (**Main text Figure 4**, **Supplementary Figure 18**). This is because, scenarios in which $MAF_{j,M_{control}} < \epsilon_{fp}$ result in highly variable regression estimates of betas and associated standard errors, whereas the SaLN-R algorithm.

#### 1.7 Ad-hoc estimator

The ad-hoc estimator for logistic regression was proposed by[3] uses contingency table information:

| | **Number of effect alleles** | | | |
|---|---|---|---|---|
| | 0 | 1 | 2 | Total |
| **Case** | $r_{j0}$ | $r_{j1}$ | $r_{j2}$ | $R_j$ |
| **Control** | $s_{j0}$ | $s_{j1}$ | $s_{j2}$ | $S_j$ |
| **Total** | $n_{j0}$ | $n_{j1}$ | $n_{j2}$ | $N_j$ |

$$\hat{\beta}_{ad-hoc} = \frac{{r_1 s_0}/{N_{01}} + {r_2 s_1}/{N_{12}} + 4\left({r_2 s_0}/{N_{02}}\right)}{{r_0 s_1}/{N_{01}} + {r_1 s_2}/{N_{12}} + 4\left({\sqrt{r_2 s_0 r_0 s_2}}/{N_{02}}\right)}$$

with

$$N_{jlm} = n_{jl} + n_{jm}.$$

Corresponding p-values can be approximated by computing standard error estimates via the linked $\chi^2_{G_j}$ statistic, i.e.

$$\hat{\sigma}_{\beta_{ad-hoc}} = \frac{\left|\hat{\beta}_{ad-hoc}\right|}{\sqrt{\chi^2_{G_j}}},$$

where

$$\chi^2_{G_j} = \frac{2N_j\left(N_j\left(r_{j1} + 2r_{j2}\right) - R_j\left(n_{j1} + 2n_{j2}\right)\right)^2}{R_j(N_j - R_j)\left(N_j\left(n_{j1} + 4n_{j2}\right) - \left(n_{j1} + 2n_{j2}\right)^2\right)}.$$

We use these to form an ad-hoc $Z$-score in our simulation studies, i.e.,

$$Z_{ad-hoc} = \frac{\hat{\beta}_{ad-hoc}}{\hat{\sigma}_{\beta_{ad-hoc}}}.$$

#### 1.8 Population substructure, cryptic confounding: genomic inflation adjustment via effective sample-size

For genome-wide uses of isGWAS we assess values for the genomic inflation factor $\lambda_{GC}$ via:

$$\lambda_{gc} = \frac{\text{median}(\chi^2_{isGWAS})}{0.454}$$

Noting that, in the computation of the regression parameters $\{\alpha_{j,M}, \beta_{j,M}\}$, the standard isGWAS algorithm is invariant to the choice of sample size, we can therefore efficiently adjust the isGWAS standard error and Z-score using the following identity:

$$Z_{j,M,\lambda_{GC}} = \frac{\hat{\beta}_{j,M}}{\sqrt{\lambda_{GC,M}}\hat{\sigma}_{j,M}} = \frac{Z_{j,M}}{\sqrt{\lambda_{GC,M}}}$$

which is equivalent to running the original analyses with the 'effective sample-size' $N_j^{(eff)}$, i.e.,

$$N_j^{(eff)} = \frac{N_j}{\lambda_{gc}},$$

$$\hat{\sigma}^{(eff)}{}_{\beta_{j(k+1)},M} = \sqrt{\frac{e_{j,M}^{(0,k)}}{N_j^{(eff)}\left(e_{j,M}^{(0,k)} e_{j,M}^{(2,k)} - \left(e_{j,M}^{(1,k)}\right)^2\right)}}$$

$$= \hat{\sigma}_{\beta_{j(k+1)},M}\sqrt{\lambda_{gc}}.$$

P-values computed using $Z_{j,M,\lambda_{GC}}$ might account for some cryptic relatedness and population substructure that can arise in the user defined data $\{N_j, N_j^*, MAF_{j,M}, MAF_{j,M}^*\}$, for each variant $j = 1,2,\ldots,Q$. Note that our use of the genomic inflation adjusted sample size, i.e., the effective sample-size, is similar to the adjusted sample used in the meta-analysis of Z-scores in LD-Score Regression[4]. In our real-data analyses, we instead perform robust QC

(**Methods**) to attenuate the need to deploy a genomic inflation adjustment. Nevertheless, genomic inflation modified Z-score and corresponding p-values might be considered as part of routine sensitivity analyses.

##### 1.9 Recommended Quality Control for genetic Individual Level Data (ILD) to running successfully isGWAS

Variable selection:

1. MAF=0.001 threshold
2. Hardy-Weinberg equilibrium test threshold $p = 10^{-15}$
3. Genotyping quality rate above 99%
4. Genotype missingness under 10%
5. Minor allele frequency count above 10

Cohort selection:

1. Use principal components of ancestral analysis to select a homogeneous population using a clustering procedure.
2. Drop any subjects with mismatch between reported and genetic sex, or with evidence for sex chromosome aneuploidy.
3. Drop any subjects with missingness above 10%.
4. Obtain kinship information using KING software[5] and select maximum number of unrelated cohort using network-driven strategy[6] while favouring cases over controls.

#### 2 Assessing predictive capabilities of isGWAS

#### 2.1 Leapfrog re-sampling: using isGWAS to extrapolate variant association results to future sample sizes

Recall that $N_j$ denotes the current sample size for the $j$th variant. Our approach leverages variation in both genotype and disease status between individuals (which might resemble variation present in new additional samples) and is summarised in 4-steps: (1) Specify number $K$, sub-sample $0 < \gamma_1 < 1$ and target sample $\gamma_2 > 1$ parameters; (2) generate $K$ random sub-samples of individuals of size $\gamma_1 N_j$; (3) (leapfrog-step) compute sufficient statistics in the sub-sample and re-scale the estimated number of cases and controls to match the larger virtual target sample size (i.e., multiply case and control counts by ${}^{\gamma_2}/_{\gamma_1}$); and (4) deploy isGWAS in each leapfrog sub-sample and return a summary of the association $p$-values across all $K$ samples. As a generally robust point estimate, we compute the median $p$-value in testing (ranges of values can alternatively be returned).

We run the leapfrog re-sampler in both simulation and real-data settings, informed by the 7 tested diseases in UK Biobank. As parameters $\{K, \gamma_1, \gamma_2\}$ are user defined, we evaluate performance over a range of values and consider a maximum $\gamma_2$-fold increase of $\gamma_2 = 2.5$, equating to a 150% increase in sample size relative to the current size. Results are presented in **Main text Figure 5 and Supplementary Table 11**. On assuming that a group of additional participants are reasonably well approximated by one or more sub-groups of individuals in the current sample, and taking a true positive association as $p < 5 \times 10^{-8}$ in the target sample, our results reveal that: when doubling sample size from $\mathrm{N} = 276{,}204$ to $\gamma_2\mathrm{N} = 552{,}408$, the FDR was well controlled, i.e., $FDR \lesssim 10\%$, and while accuracy and TPR dropped when increasing the target sample size (i.e., on increasing $\gamma_2$) values for each measurement were typically $\gtrsim 65\%$ across the range (**Main text Figure 5**).

#### 2.2 Theoretic sub-sampling

We perturb the original prevalence value of the sub-sample by randomly drawing it from a normal distribution with mean $\pi$ and variance $\frac{\pi(1-\pi)}{N}$ . The $MAF^*$ and $MAF - MAF^*$ values are perturbed by sampling from a normal distribution with mean $MAF$ and variance $\frac{MAF(1-MAF)}{2N} + \frac{P_{AA}-MAF^2}{2N}$ where $P_{AA}$ is the frequency of genotype $AA$ where $A$ is the minor allele. The new $MAF,\ MAF^*$ values are used to recalculate isGWAS for the full sample size and compared to the isGWAS. We also compare the results from this theoretic based prediction scenario to obtaining subsamples from the population, calculating their $MAF$ values and using the results to predict the associations in the full dataset. For each of the seven UK Biobank analysed diseases, we progress this theoretic sub-sampling approach for all variants with $p < 10^{-6}$.

The over-all performance is better than the LRS one since in all cases the introduced variability is closer to the truth, which is not guaranteed with a random sampling strategy in the LRS. See **Supplementary Figure 19**.

#### 2.3 Using real schizophrenia data for predictive purposes

In our analyses of variants in the Schizophrenia 2022 dataset[7], we refined variants by clumping via PLINK[8]. Note that, not all clumped variants from the 2022 dataset (451 in total) were present in the clumped variant set from the 2014 dataset (225 in total) – there were intersecting overlapping variants and a total of 608 variants present across both datasets (regardless of clumping strategy). Using the 225 variants available in the 2014 dataset[9], from clumping parameters ($R^2 = 0.2, p_1 = 1e-07, p_2 = 1e-07$), an application to the 2014 dataset shows inference results for p-values for autosomal significant loci (**Supplementary Figure 20**) show strong prediction results as 92.91% of estimates lie inside the original 95% C.I. of the effect (100% of estimates C.I.s lie inside the original 95% C.I.s). There is no linear association between the accuracy of the inferred p-values and any MAF driven values. Using the 451 variants with clumping parameters set ($R^2 = 0.2, p_1 = 1e-07, p_2 = 1e-07$), an application to the 2022 dataset shows inference results for p-values for autosomal significant loci (**Supplementary Figure 21**) show lower prediction results as 82.48% of estimates lie inside the original 95% C.I. of the effect (100% of estimates C.I.s lie inside the original 95% C.I.s). We use the population-level MAF information from 2014 Schizophrenia dataset for the 608 overlapping variants (present in both sets regardless of clumping strategy) and match them to the 2022 Schizoprenia dataset to obtain inferred effect and significance values. We observe good inference (**Main text, Figure 6**), supported further by the fact that 84.12% of estimates lie inside the original 95% C.I. of the effect (100% of C.I. of estimates lie inside the original 95% C.I.).

### 3 UK Biobank Comparative Results

#### 3.1 isGWAS genomic inflation

To understand the impact of stringent QC on the isGWAS results, we compare the genomic inflation in the UK Biobank dataset per chromosome (**Supplementary Figure 14**) and at the genome level (**Supplementary Table 8**).

For genome-wide uses of isGWAS we assess values for the genomic inflation factor $\lambda_{GC}$ via:

$$\lambda_{gc} = \frac{\text{median}(\chi^2_{isGWAS})}{0.454}.$$

We compare isGWAS genomic inflation to that of REGENIE with and without covariate adjustment and to a simple logistic regression model without covariate adjustment. We note that isGWAS has lower genomic inflation to that of the other models implying that the QC procedure in combination with the isGWAS approach which operates at the population-level, surpass the effects of genomic inflation posed by working with individual level data.

#### 3.2 Performance of isGWAS compared to REGENIE without isGWAS enhanced QC

For all subjects in UK Biobank with ethnical status ‘white British’ and no reported discrepancy between genetic and reported sex, we select the full set of subjects regardless of relatedness status, which is a total of 408,110 subjects. In addition, we produce three more cohorts: a maximum unrelated cohort per disease where we favour cases over controls to keep in the cohort, a cohort of subjects with up to and including third degree relatedness, a cohort of subjects with up to and including second degree relatedness. To obtain these three cohorts, we have used the kinship information as provided by the UK Biobank[10], recommended kinship thresholds[5], and a network inspired cohort selection strategy[6] that we adapted to favour cases over controls for different degrees of relatedness. Number of cases and controls analysed per disease for each of the four cohorts is provided in **Supplementary Table 5**. For each of the seven diseases and cohorts, we run a one-to-one comparison between isGWAS and REGENIE-Firth. The set of variables used was 1,333,940 – this is the set of pruned variables using PLINK[8] with the following parameters: genotype quality>0.99, MAF>0.01, HWE $p < 10e-15$, 1000 bp windows, 100 variant increments, $R^2 > 0.9$. For all added degrees of relatedness, REGENIE and isGWAS (both with Firth correction) perform almost equally well as observed from the p-p value plots and the corresponding correlation value (**Supplementary Figure 12**). This is further supported by the accuracy/TPR/FPR rates (**Supplementary Table 6**).

#### 3.3 Assessing sensitivity to imputation of genotypes

There is a small additional real-terms computational cost of running isGWAS with and without imputed values for missing genotypes. We therefore assessed sensitivity of results to imputation for each of the 7 diseases considered In UK Biobank, by additionally performing analyses using the raw genotypic data (, i.e., for each variant we removed any participants with missing values under a missing completely at random assumption) and compare to REGENIE imputed values. isGWAS uses MAFs for cases and controls obtained from the non-missing values for each variant, whereas REGENIE and other classical methods impute those missing values to prevent dropping case. On contrasting results with our main analyses (**Figures 2-3, Main text**), which used REGENIE imputed genotypes, we observe some surprising results (**Supplementary Figure 15**). Imputation occasionally led to modified changes in the MAF between cases and controls such that estimated genetic effects switched sign (i.e., effect direction) relative to results computed from non-imputed data (**Supplementary Figure 15**). This seems to occur when the raw MAF is close to 0.5 (see **Supplementary File 3** for a full list of variants and their characteristics). We question the suitability of interpreting results from these variants and their use in subsequent downstream investigations (e.g.,) meta or colocalization analyses, where effect direction can be important. Hence, isGWAS can be routinely deployed to assess sensitivity to imputation and consequently the reliability of results for variants in analyses. The correlations for the p-values and effect size values between isGWAS-Firth and REGENIE-Firth are provided in **Supplementary Table 3**. The full results are included in **Supplementary File 2**. The specific results for opposite  results are in **Supplementary File 3**. From all significant variants ($p<0.05$) identified by REGENIE and isGWAS (imputed MAFs) we have identified a few of those, but due to the nature of the MAF change, the list is not exhaustive. These are: for I10 - 273 out of 270,365; for J45 - 154 out of 226445, for I25 - 94 out of 149674, for H40 - 246 out of 153465, for I63 - 36 out of 94498, for C18 - 28 out of 100608, for C73 - 178 out of 86144. As we see some of these are significant and could lead to false positives.

##### 3.4 Assessing isGWAS in the full UK Biobank cohort in the presence of relatedness and mixed ethnicity

We conduct an isGWAS comparison to REGENIE performance where all 486,378 UK Biobank subjects with available covariate information were included in the analysis to assess the performance of isGWAS when mixed ancestry is present in the cohort. The analysis was conducted on the same 1,333,940 pruned variants as the ones used in the relatedness investigation. As expected, performance in recovering the same set of significant variants is dropping (**Supplementary Table 7**) and from a p-p value plot exploration (**Supplementary Figure 13**) we note that isGWAS tends to underinflate p-values. This example highlights the need for a stringent QC procedure before using isGWAS.

#### 4 Simulation study

To assess the ability of isGWAS and isGWAS-Firth to emulate results from classical logistic and Firth corrected regression, using only sample-level summary data $\{N_j, N_j^*, MAF_{j,M}, MAF_{j,M}^*\}$, we performed two simulation studies: (I) disease status is simulated via a logistic link-function conditional on a non-zero association between variant and disease (**Supplementary Figure 18**); and (II) a 'model-free' simulation where disease status for each individual is manually assigned - in the absence of a statistical model - and genotype data are subsequently simulated conditional on disease status across a range of $MAF_{j,M}$ and $MAF_{j,M}^*$ values (**Main text Figure 4**). Briefly, simulation I assesses concordance of results when the 'ground-truth' variant-disease association is known, while simulation II bypasses the need to specify a disease model and corresponding genetic association parameter (e.g., a log-odds ratio), achieved instead by varying allelic frequency in both cases and controls, respectively. Simulation II allows for scenarios in which there is a single diseased individual in the sample and separately, a single individual with the effect allele (note these scenarios are allowed to co-occur). This is to evaluate whether the novel sample-level optimization procedure of isGWAS can return robust results in the rarer variant and prevalence spectrum, (e.g.,) helping to attenuate the so-called blow-up of log-odds ratios (betas) that is evident in individual-level data optimization procedures of highly sparse genotypic and disease data (i.e., rare disease and allelic frequencies).

In each simulation study we explore ultra-rare to common variant frequencies, as 100 random draws from the range $MAF, MAF^* \in [0.0001, 0.5]$, and 50 replicas for the rare disease prevalence $\pi \in [0.0001, 0.5]$. Note, disease prevalence of $\pi = 0.5$ is chosen to reflect a balanced case-control study. In both I and II, we randomly sampled sample sizes from $N \in \{10^4, 1.5 \times 10^4, 2 \times 10^4, \ldots, 10^5\}$ a total of 100 times. In total we generated 25,000 datasets for each study. For Simulation I we also compare results to the Ad-hoc estimator for logistic regression and the Fisher exact test to highlight various advantages and disadvantages of the different procedures.

We generated simulated datasets to assess performance of isGWAS - with and without Firth correction - against a variety of classical methods which either: (a) do not require ILD, the logistic ad-hoc estimator[3] and Fisher's Exact Test[11]; or (b) require ILD, logistic and Firth corrected regression[1]. We perform two simulation studies which, for each disease prevalence $\pi \in \{10^{-4}, 10^{-3}, 0.01, 0.1, 0.5\}$, we generate $k = 1,2, \ldots, 1000$ datasets with sample size $N_k$ randomly drawn from the set $N_k \in \{10^4, 2 \times 10^4, \ldots, 10^5\}$. In the first study, for each simulated individual we randomly generate disease status under a logit model and fix the genetic effect on disease so that the ground truth is known. In the second study, we do not specify a model for disease status and allow the genetic effect to vary between each dataset. In both scenarios, the range of disease prevalence and minor allele frequencies broadly match ranges computed in our UKB analyses. We also included more extreme values to help stress test isGWAS in ultra-rare disease and MAF settings. A total of 25,000 datasets were considered in each study and results are presented in **Main text Figure 4**, **Supplementary Figure 18** and **Supplementary Tables 19-21**.

isGWAS-Firth outperformed all other approaches in terms of either computational cost or robustness of results over the range of scenarios considered. For example, when compared to ILD Firth regression, isGWAS-Firth accurately matched estimates of: (a) genetic effects, median absolute relative error (MARE) of isGWAS-Firth relative to ILD Firth was 0.28%; (b) standard errors, MARE=0.25%; and (c) $p$-values, MARE= 0.4%. isGWAS-Firth computed parameter estimates around 800 times faster than the ILD Firth regression[12], i.e., median absolute relative difference in computational time between ILD Firth and isGWAS-Firth was 80,000% (see **Supplementary Table 19** for more information). As anticipated[13], when disease prevalence is rare (i.e., $\pi \leq 0.01$) parameter estimates computed using non-Firth corrected ILD regression were unreliable. The MSE and distribution of parameters estimated via ILD logistic regression were often orders of magnitude poorer than other methods (**Main text, Figure 4a-c**). In scenarios where ILD logistic regression performed poorly, the non-ILD contingency-table based analogues performed equally poorly – regularly failing to compute genetic effect estimates or $p$-values. Failure rates of the logistic ad-hoc estimator were 53.8% and 27.9% when disease prevalence was 1 in 10,000 or 1 in 1,000, respectively (**Main text, Figure 4e**). Similarly, Fisher's Exact Test failed to compute association $p$-values in 56.1% and 27.9% of scenarios. isGWAS without Firth correction mirrors logistic regression and was thus subject to comparable failure rates of 56.1% and 27.9% (**Main text, Figure 4e**). isGWAS-Firth, however, had a failure rate of 2.6% and 0% in these scenarios, and an ultra-low failure rate of 0.04% over all scenarios (**Main text, Figure 4e**). We note that MAF in cases was 0 in the very small number of scenarios in which isGWAS-Firth failed - excluding these scenarios isGWAS-Firth had a 0% failure rate. **Main text, Figure 4f-h** highlights the chronological evolution of non-ILD $p$-value estimates, from Fisher's Exact Test (1922)[11], Sasieni's logistic ad-hoc estimator (1997)[3] to isGWAS-Firth, illustrating improvements in estimation via successive approaches.

#### 5 Computational performance

isGWAS is an iterative algorithm whose convergence depends on several tuning parameters. The default parameter settings see the isGWAS-Firth algorithm converge always in real-data settings and almost always in our simulation scenarios (**Main text, Figure 4 and Supplementary Figures 18**). Convergence is achieved in around 0.001 seconds per variant (**Supplementary Table 21**). The uncorrected isGWAS algorithm can require more iterations, particularly for diseases with lower prevalence, and has overall weaker convergence rates (**Supplementary Tables 9 and 18**). We conclude that, unlike in ILD analyses, correcting for maximum likelihood bias via a Firth correction returns: (a) more reliable results and (b) in quicker computational time. As the isGWAS algorithm is performed one variant at a time, making isGWAS highly parallelisable. The calculations of isGWAS do not require any advance statistical packages and are based on basic algebraic functions making it easy to implement and execute. The method is coded in R 4.2.0 and takes advantage of the 'parallel' package 'mclapply' function to optimize computation time. The computation time and the computation cost depend entirely on the user machine available for computation. Extensive analysis was performed on a computing environment as follows: 48 virtual CPU cores of a 2.5 GHz Intel Xeon Gold 6240R processor, 64 GB of memory. Results of the analysis comparing isGWAS with and without Firth for a different number of variants distributed over different number of CPU cores are present in **Figure 4, Main text**. See **Supplementary Table 18** for full results. The results show that if distributed over 32 CPU cores, 10 million variants may be run with Firth-corrected isGWAS in 3.5 minutes. This is already a significant improvement over available state-of-the-art methods such as REGENIE[14] which for one binary trait requires a minimum average CPU Time of 5.4 hours to test 679,298 SNPs for association (training on 383,009 in the first step) for a UK Biobank population (~400K). We also observe that the time needed to run the analysis is affected by the prevalence – lower prevalence leads to longer convergence times in some situations. Full details are available in **Supplementary File 4**.

### Supplementary Figures

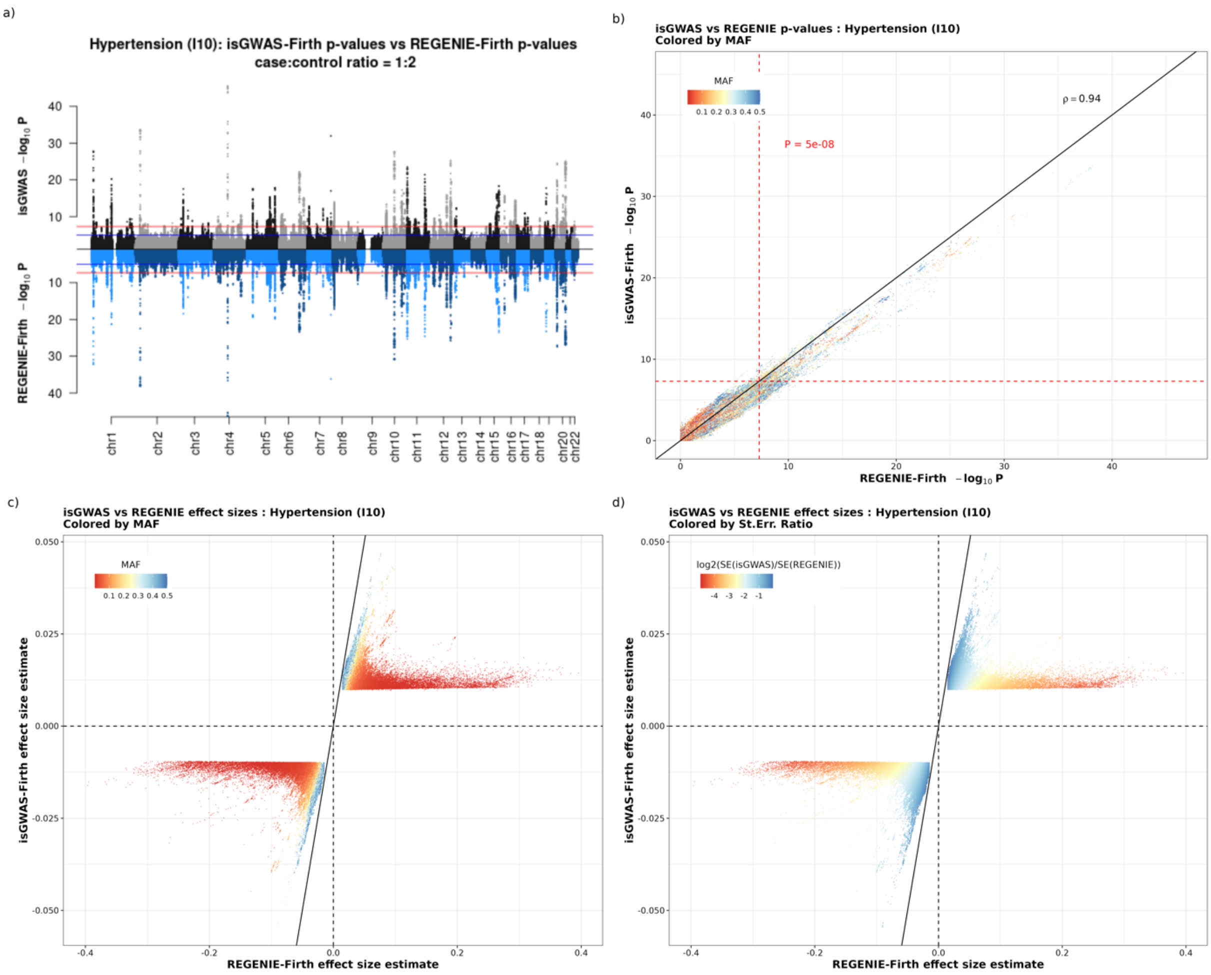

*Supplementary Figure 1. Comparative results for Hypertension (IC10:I10) from UK Biobank. Subplot (a) is a mirror Manhattan plot comparing $-\log_{10} P$ values for isGWAS and REGENIE-Firth, (b) plots $-\log_{10} P$ values for isGWAS and REGNIE-Firth with the standard threshold P-value indicated. Subplots (c) and (d) showcase $\beta - \beta$ effect size estimates for variants with p-values<0.05 and are coloured by population-level MAF and $\log_2(\frac{SE(isGWAS)}{SE(REGENIE)})$.*

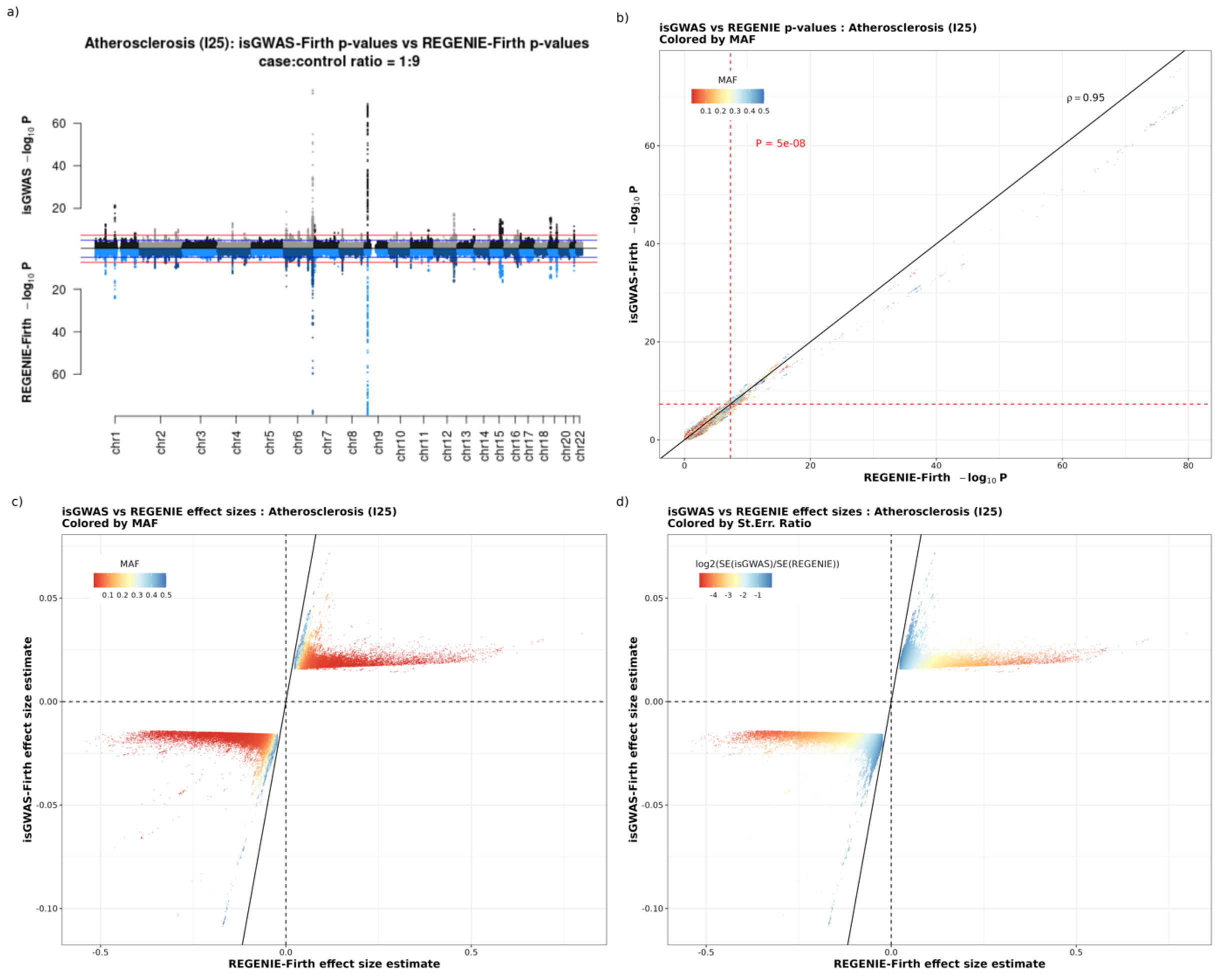

*Supplementary Figure 2. Comparative results for Atherosclerosis (IC10:I25) from UK Biobank. Subplot (a) is a mirror Manhattan plot comparing $-\log_{10} P$ values for isGWAS and REGENIE-Firth, (b) plots $-\log_{10} P$ values for isGWAS and REGNIE-Firth with the standard threshold P-value indicated. Subplots (c) and (d) showcase $\beta - \beta$ effect size estimates for variants with p-values<0.05 and are coloured by population-level MAF and $\log_2(\frac{SE(isGWAS)}{SE(REGENIE)})$.*

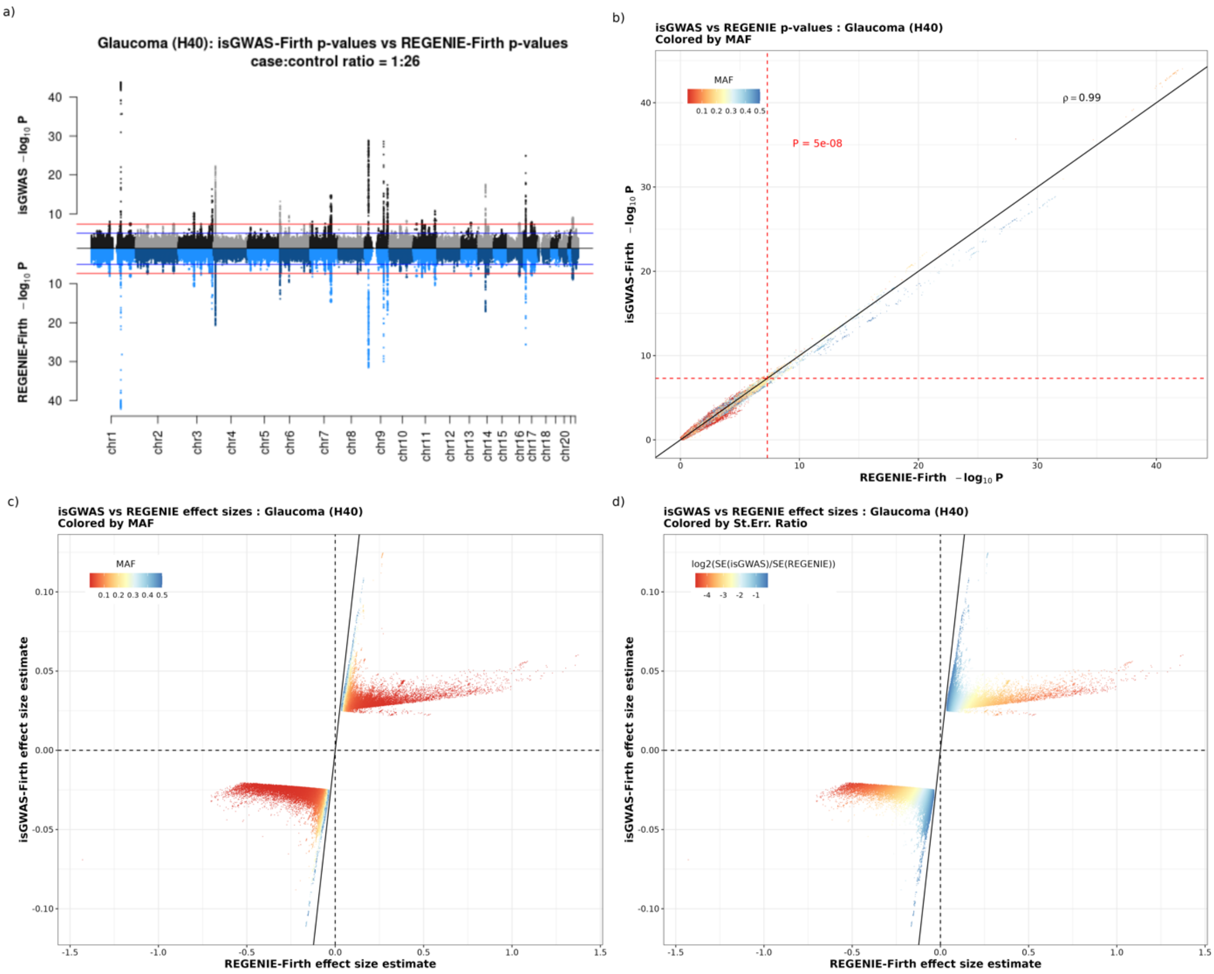

*Supplementary Figure 3. Comparative results for Glaucoma (IC10:H40) from UK Biobank. Subplot (a) is a mirror Manhattan plot comparing $-\log_{10} P$ values for isGWAS and REGENIE-Firth, (b) plots $-\log_{10} P$ values for isGWAS and REGNIE-Firth with the standard threshold P-value indicated. Subplots (c) and (d) showcase $\beta - \beta$ effect size estimates for variants with p-values<0.05 and are coloured by population-level MAF and $\log_2(\frac{SE(isGWAS)}{SE(REGENIE)})$.*

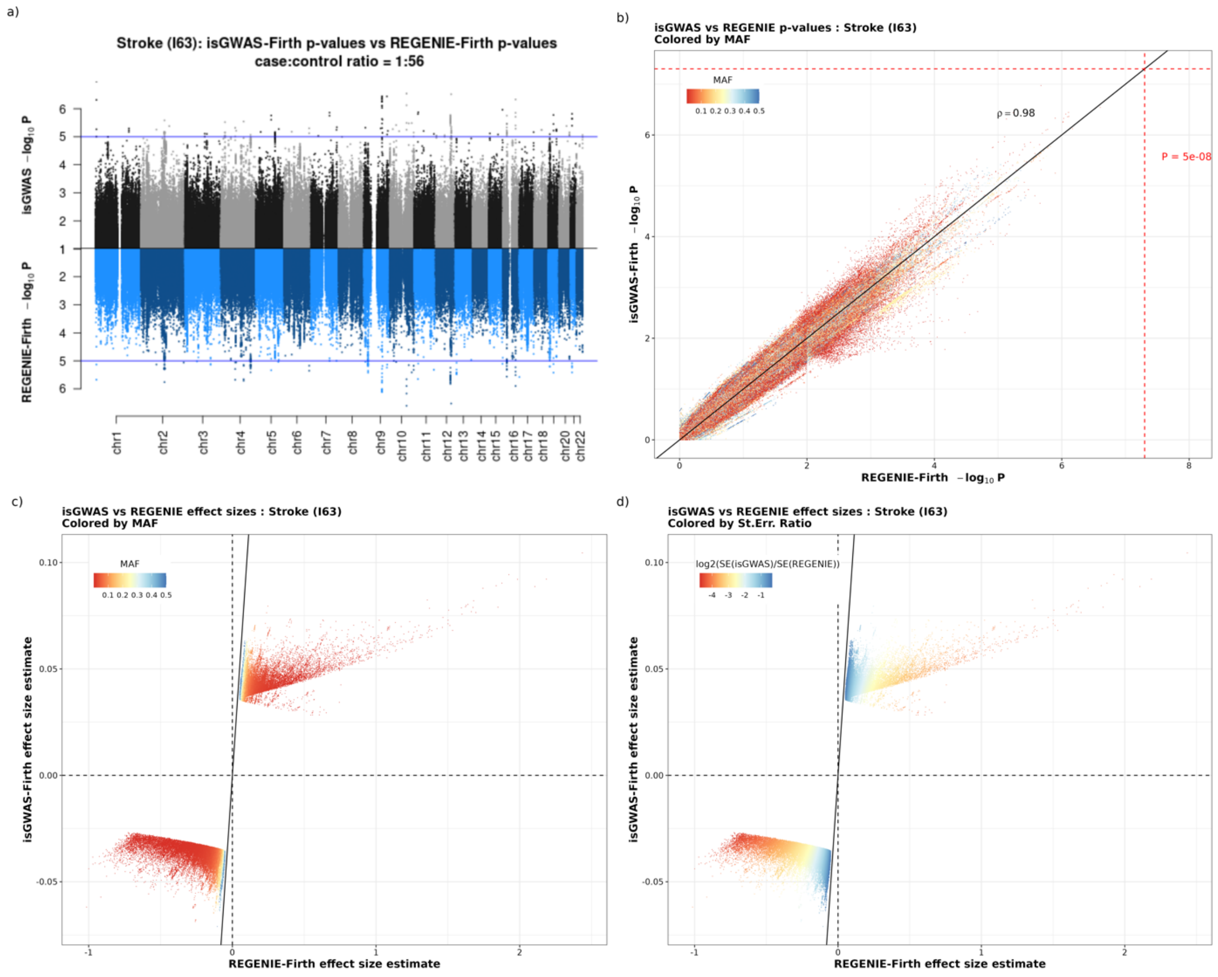

*Supplementary Figure 4. Comparative results for Stroke (IC10:I63) from UK Biobank. Subplot (a) is a mirror Manhattan plot comparing $-\log_{10} P$ values for isGWAS and REGENIE-Firth, (b) plots $-\log_{10} P$ values for isGWAS and REGNIE-Firth with the standard threshold P-value indicated. Subplots (c) and (d) showcase $\beta - \beta$ effect size estimates for variants with p-values<0.05 and are coloured by population-level MAF and $\log_2(\frac{SE(isGWAS)}{SE(REGENIE)})$.*

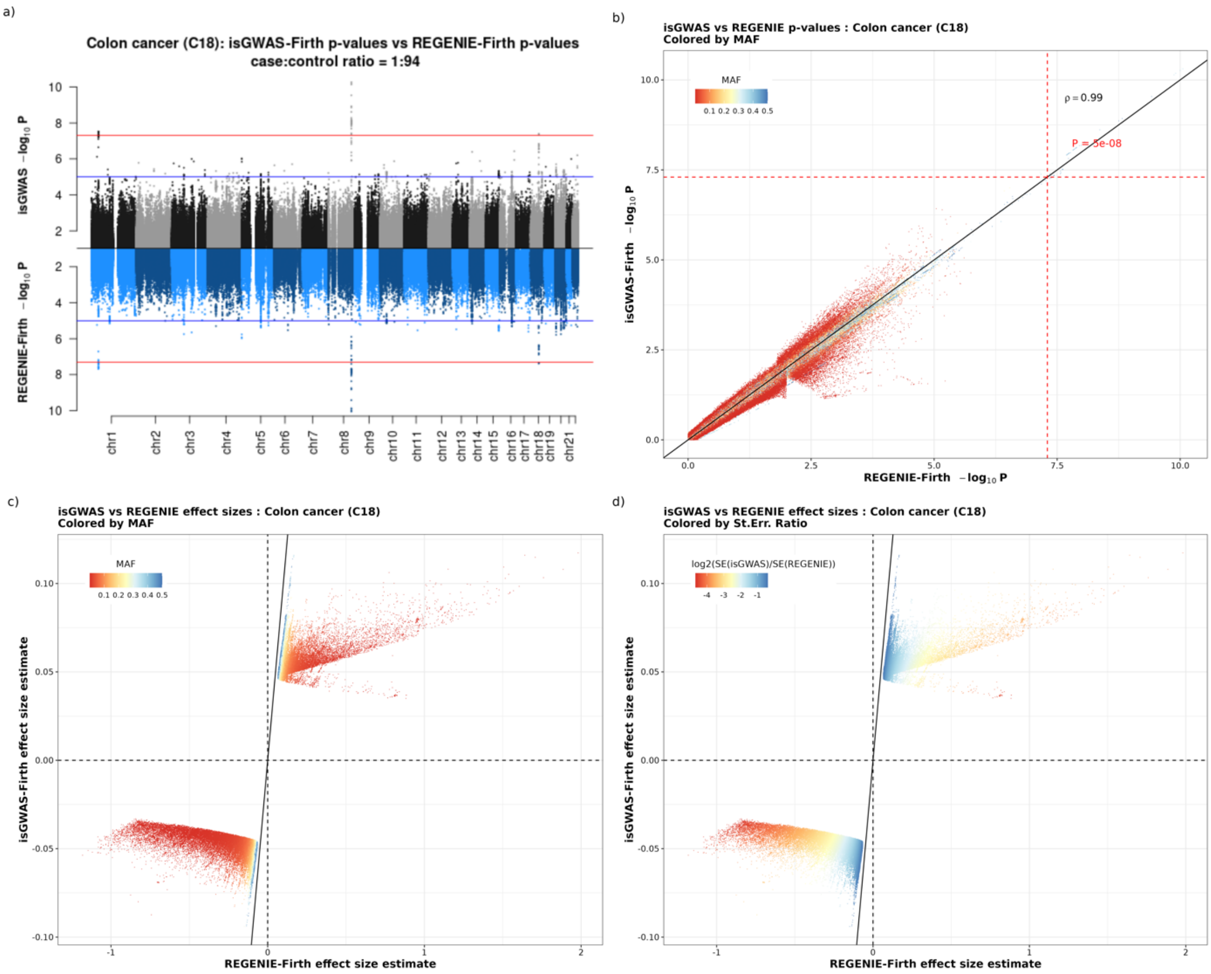

*Supplementary Figure 5. Comparative results for Colon cancer (IC10:C18) from UK Biobank. Subplot (a) is a mirror Manhattan plot comparing $-log_{10}P$ values for isGWAS and REGENIE-Firth, (b) plots $-log_{10}P$ values for isGWAS and REGNIE-Firth with the standard threshold P-value indicated. Subplots (c) and (d) showcase $\beta-\beta$ effect size estimates for variants with p-values<0.05 and are coloured by population-level MAF and $log_2(\frac{SE(isGWAS)}{SE(REGENIE)})$.*

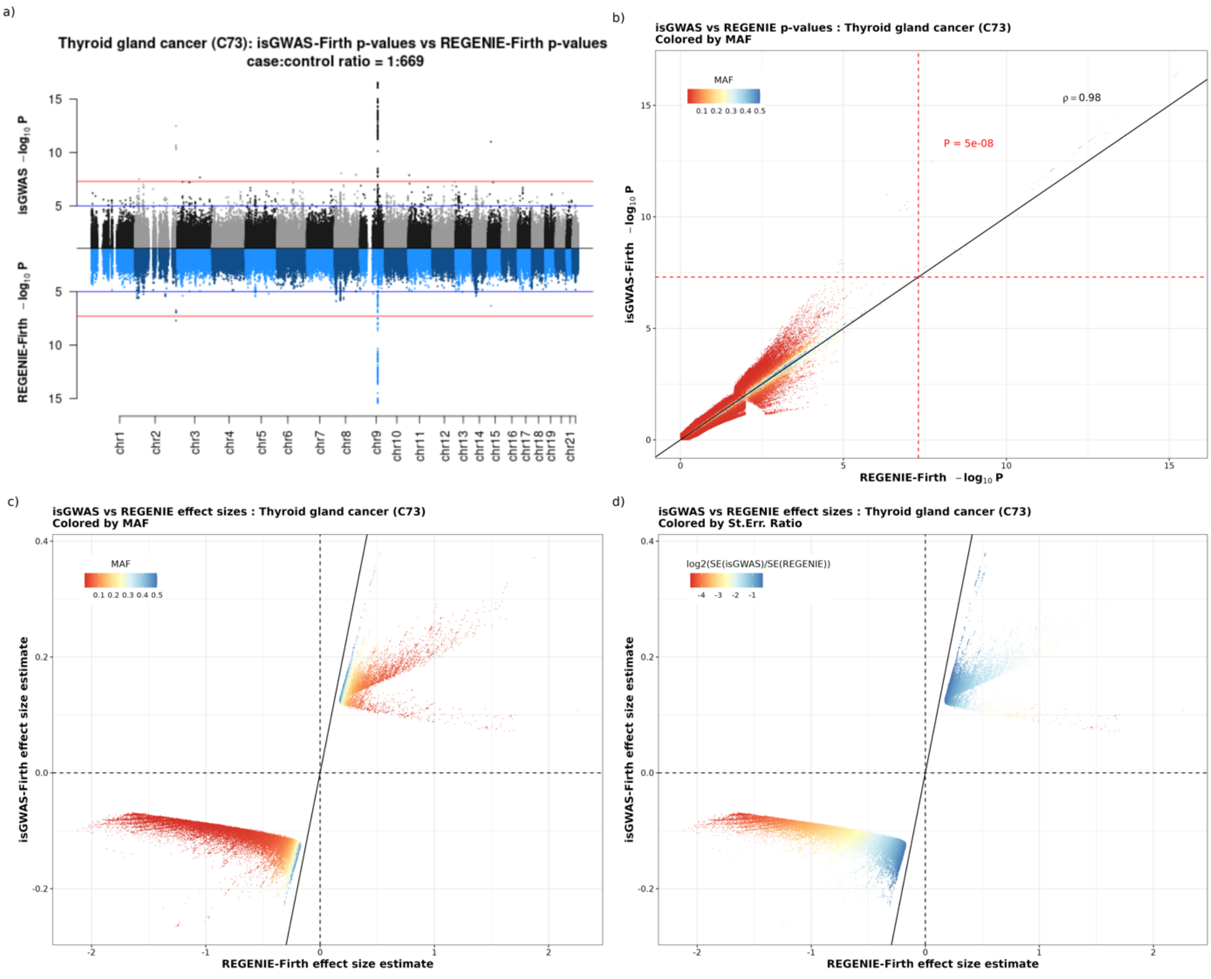

*Supplementary Figure 6. Comparative results for Thyroid gland cancer (IC10:C73) from UK Biobank. Subplot (a) is a mirror Manhattan plot comparing $-log_{10} P$ values for isGWAS and REGENIE-Firth, (b) plots $-log_{10} P$ values for isGWAS and REGNIE-Firth with the standard threshold P-value indicated. Subplots (c) and (d) showcase $\beta - \beta$ effect size estimates for variants with p-values<0.05 and are coloured by population-level MAF and $log_2(\frac{SE(isGWAS)}{SE(REGENIE)})$.*

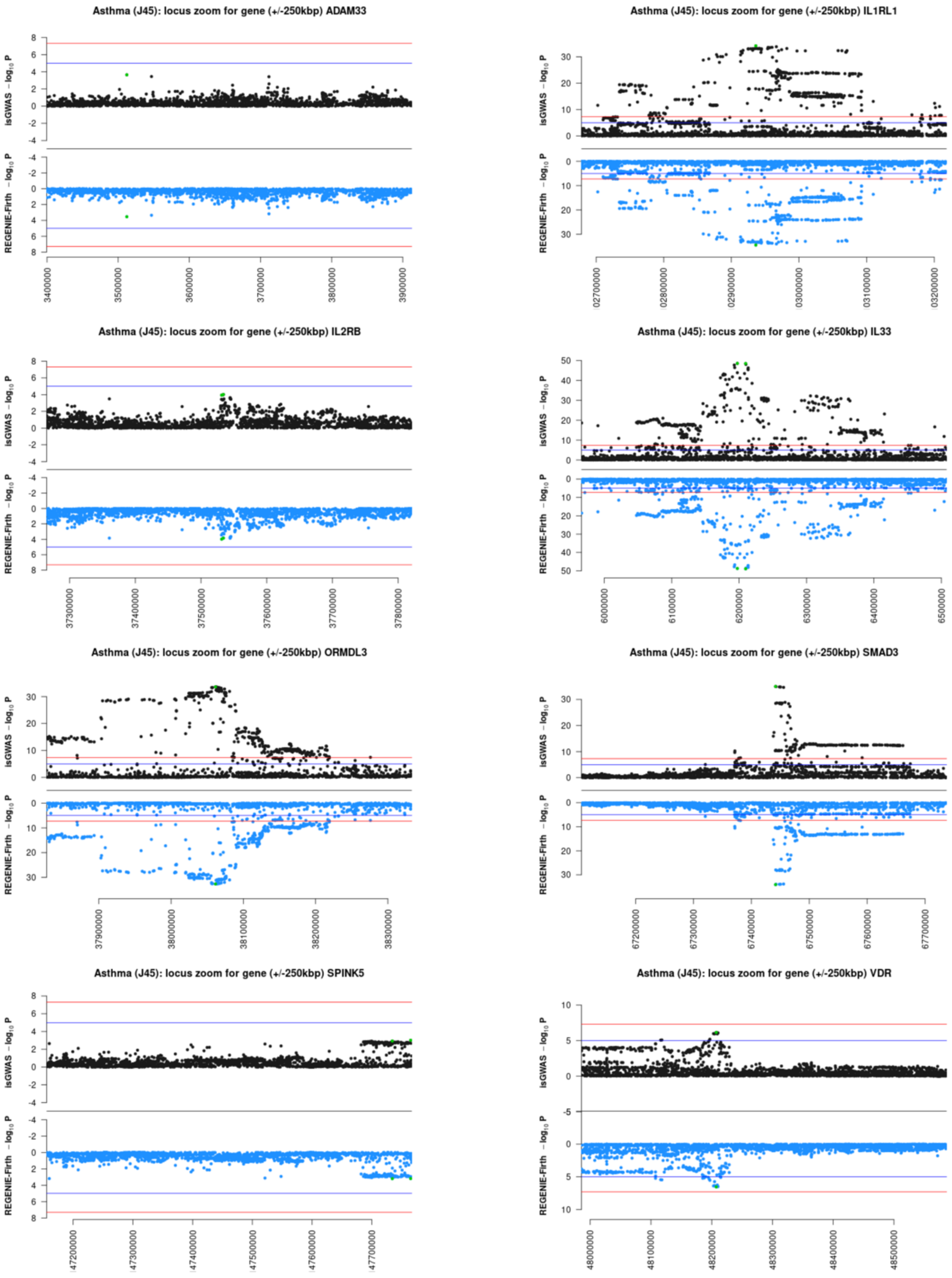

*Supplementary Figure 7. Locus zoom plots for the isGWAS and REGENIE results around eight genes known to have strong association to Asthma. Each figure is a locus zoom of the gene of interest +/-250kbp. The green variants on the plots correspond to those with lowest p-value in the plotted region.*

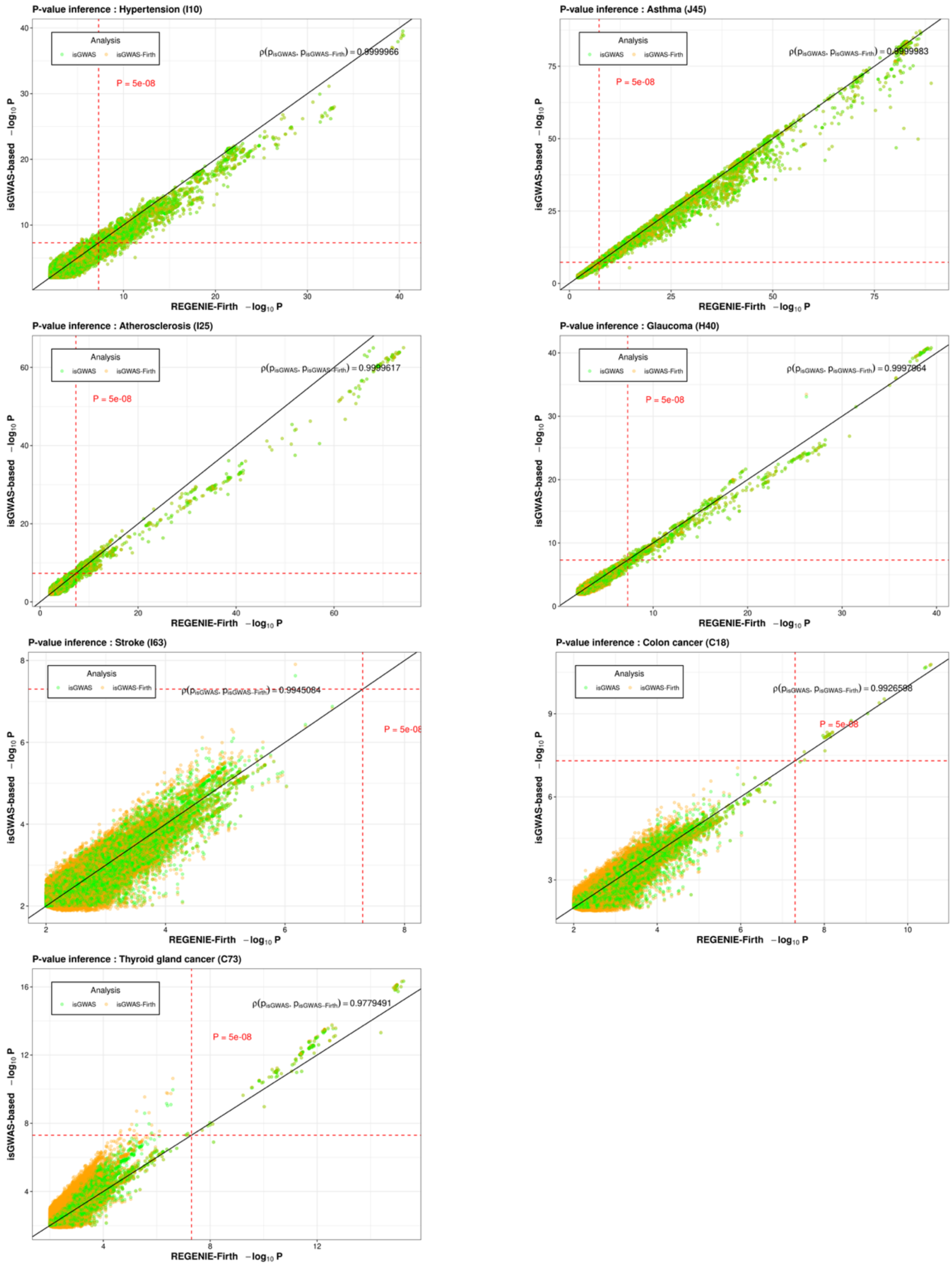

*Supplementary Figure 8. Performance of isGWAS with Firth correction. Representation of the values of* $-\log_{10} P$ *for isGWAS and REGENIE-Firth and the values for* $-\log_{10} P$ *for isGWAS-Firth and REGENIE-Firth. The classic threshold P-value is added for baseline.*

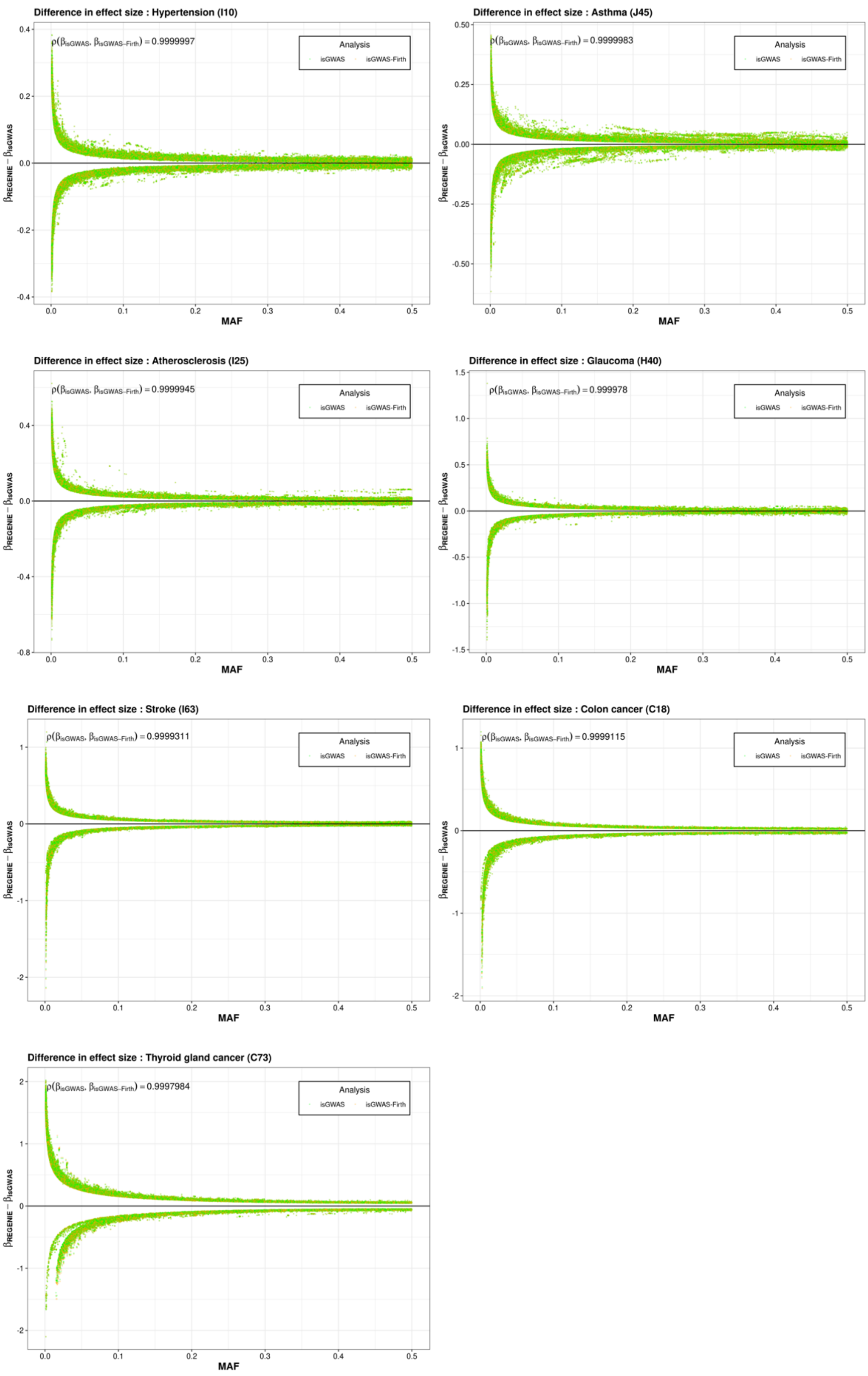

*Supplementary Figure 9. Performance of isGWAS with Firth correction. Difference between effect estimates for isGWAS and REGENIE-Firth and between effect estimates for isGWAS-Firth and REGENIE-Firth. The values $\beta_{REGENIE} - \beta_{isGWAS}$ are plotted against the population-level MAF of the variant.*

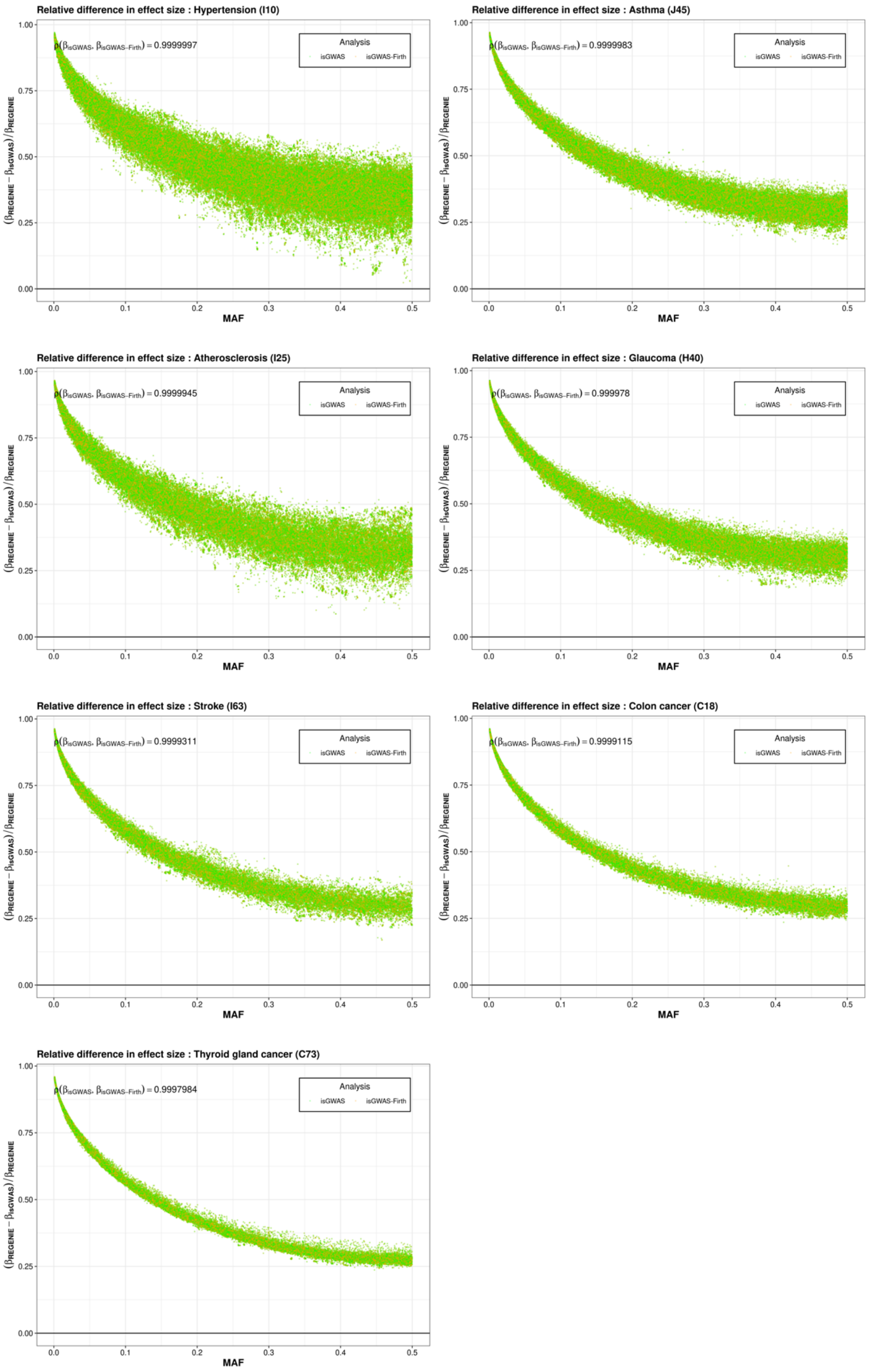

*Supplementary Figure 10. Performance of isGWAS with Firth correction. Difference between effect estimates for isGWAS and REGENIE-Firth and between effect estimates for isGWAS-Firth and REGENIE-Firth. The values $\beta_{REGENIE} - \beta_{isGWAS}$ are plotted against the population-level MAF of the variant.*

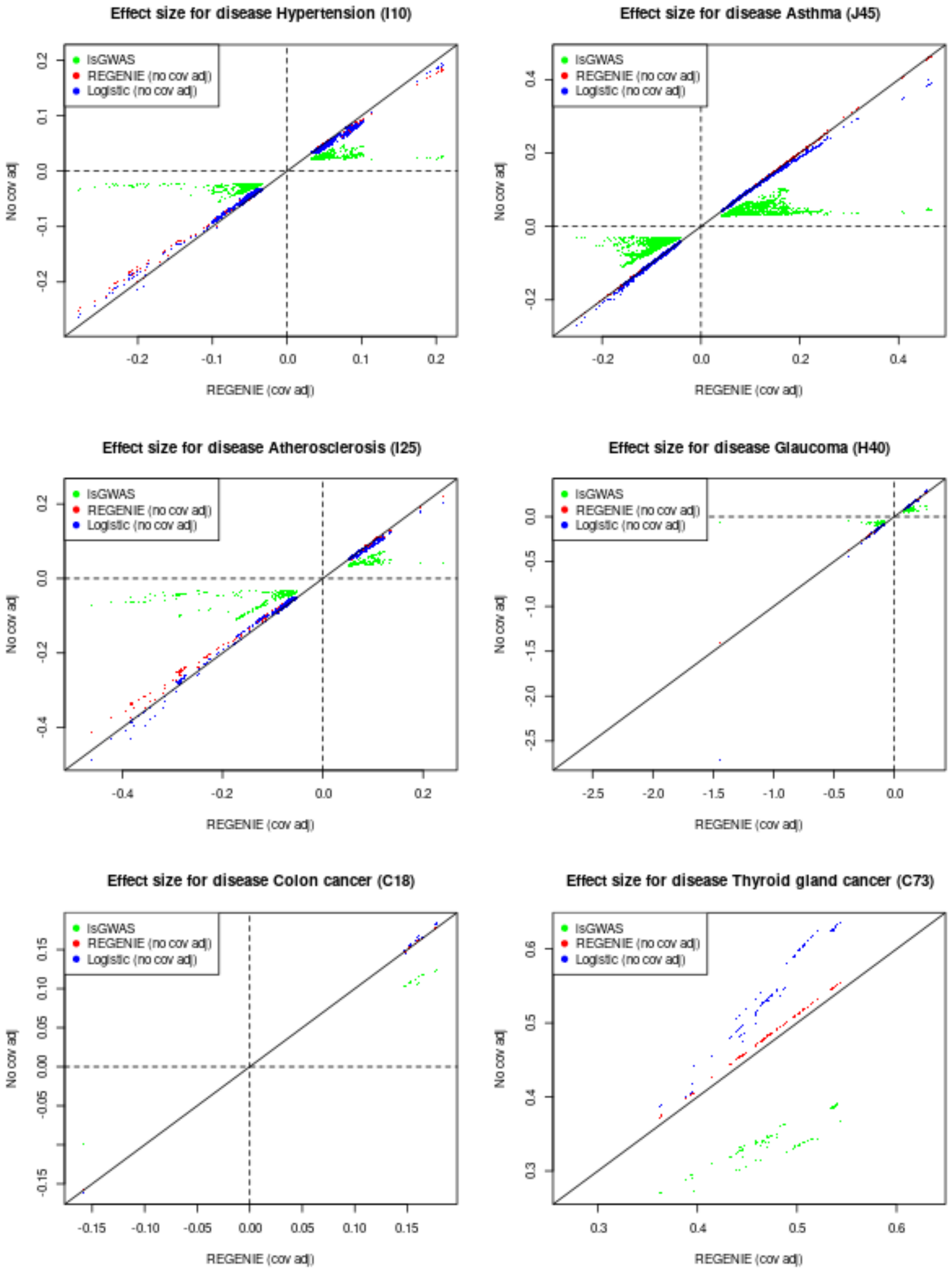

*Supplementary Figure 11. Effect size comparison for REGENIE with covariates to REGENIE without covaraites, isGWAS and a plain logistic model for UK Biobank diseases with different prevalences. Only effect sizes corresponding to variants with p<1e-07 have been plotted.*

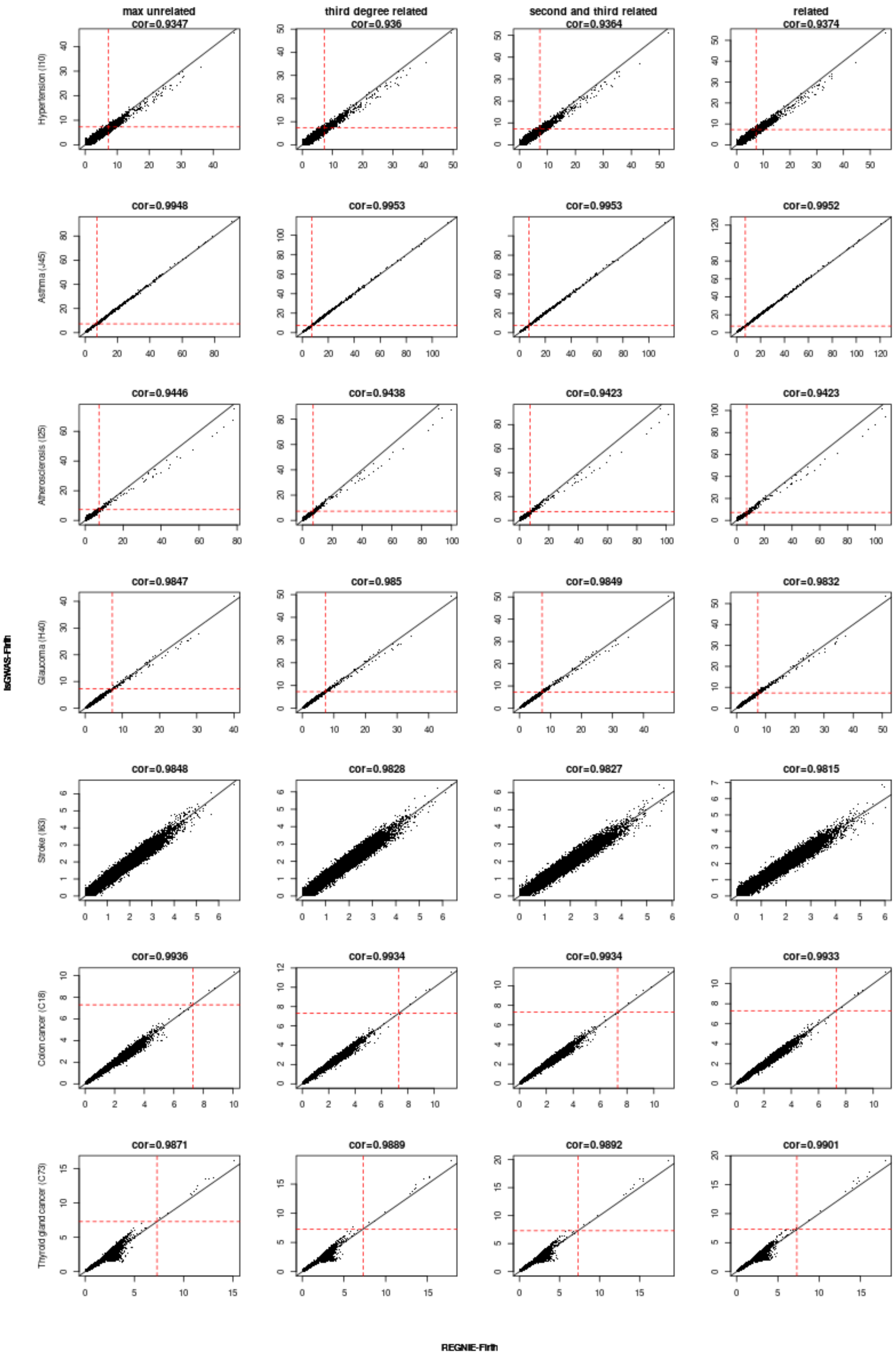
max unrelated
third degree related
second and third related
related
cor=0.9347
cor=0.936
cor=0.9364
cor=0.9374
cor=0.9948
cor=0.9953
cor=0.9953
cor=0.9952
cor=0.9446
cor=0.9438
cor=0.9423
cor=0.9423
cor=0.9847
cor=0.985
cor=0.9849
cor=0.9832
cor=0.9848
cor=0.9828
cor=0.9827
cor=0.9815
cor=0.9936
cor=0.9934
cor=0.9934
cor=0.9933
cor=0.9871
cor=0.9889
cor=0.9892
cor=0.9901
Hypertension (I10)
Asthma (J45)
Atherosclerosis (I25)
Glaucoma (H40)
Stroke (I63)
Colon cancer (C18)
Thyroid gland cancer (C73)
lsGWAS-Firth
REGNIE-Firth

Supplementary Figure 12. REGENIE-Firth vs isGWAS-Firth p-value results for seven UK Biobank diseases for 1,333,940 variants across different number of white British subjects determined by different degrees of relatedness in the following order: maximum number of unrelated subjects (where cases were retained), adding third degree relatedness, adding second and third degree relatedness, all white British subjects. Red dotted line represented  p-value threshold. Black solid line represents the 0-intercept, 1-slope line. Correlations of p-values between REGENIE and isGWAS are also provided.

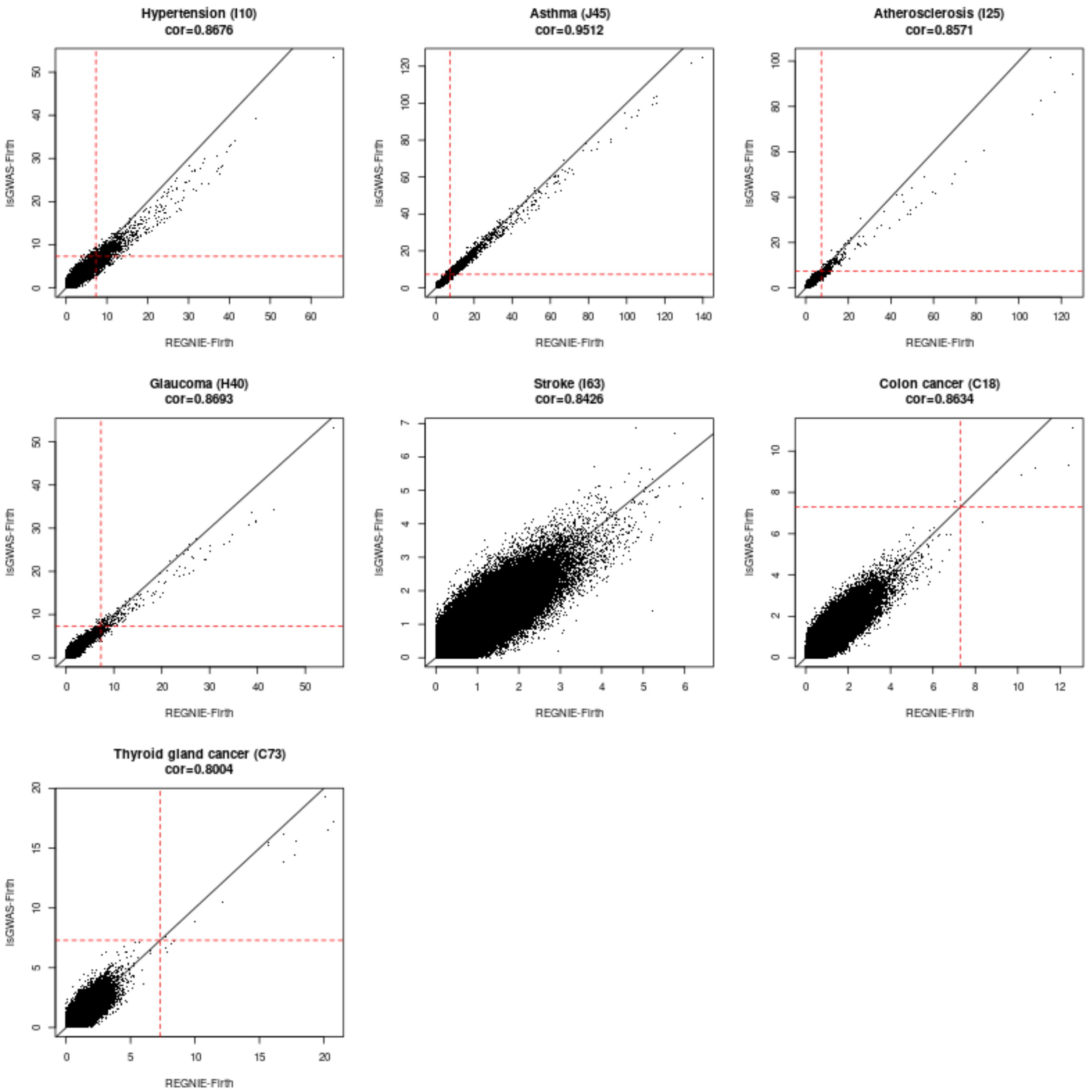

Supplementary Figure 13. REGENIE-Firth vs isGWAS-Firth p-value results for seven UK Biobank diseases for 1,333,940 variants for 486,378 UK Biobank subject with available covariate information. The subjects have not been filtered neither for ethnicity nor for relatedness.

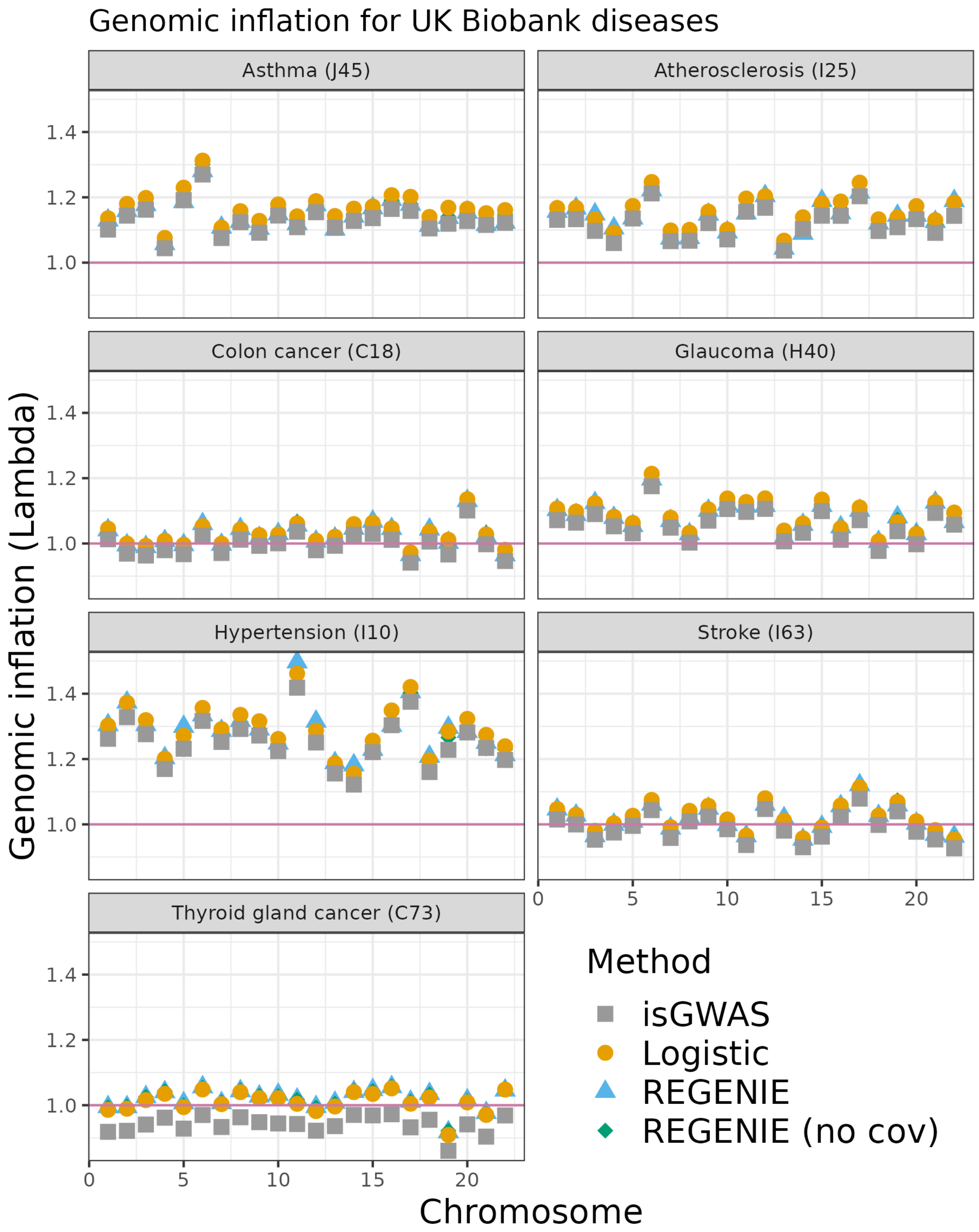

*Supplementary Figure 14. Genomic inflation per chromosome comparing isGWAS to REGENIE with and without covariate adjustment and a logistic model without covariate adjustment.*

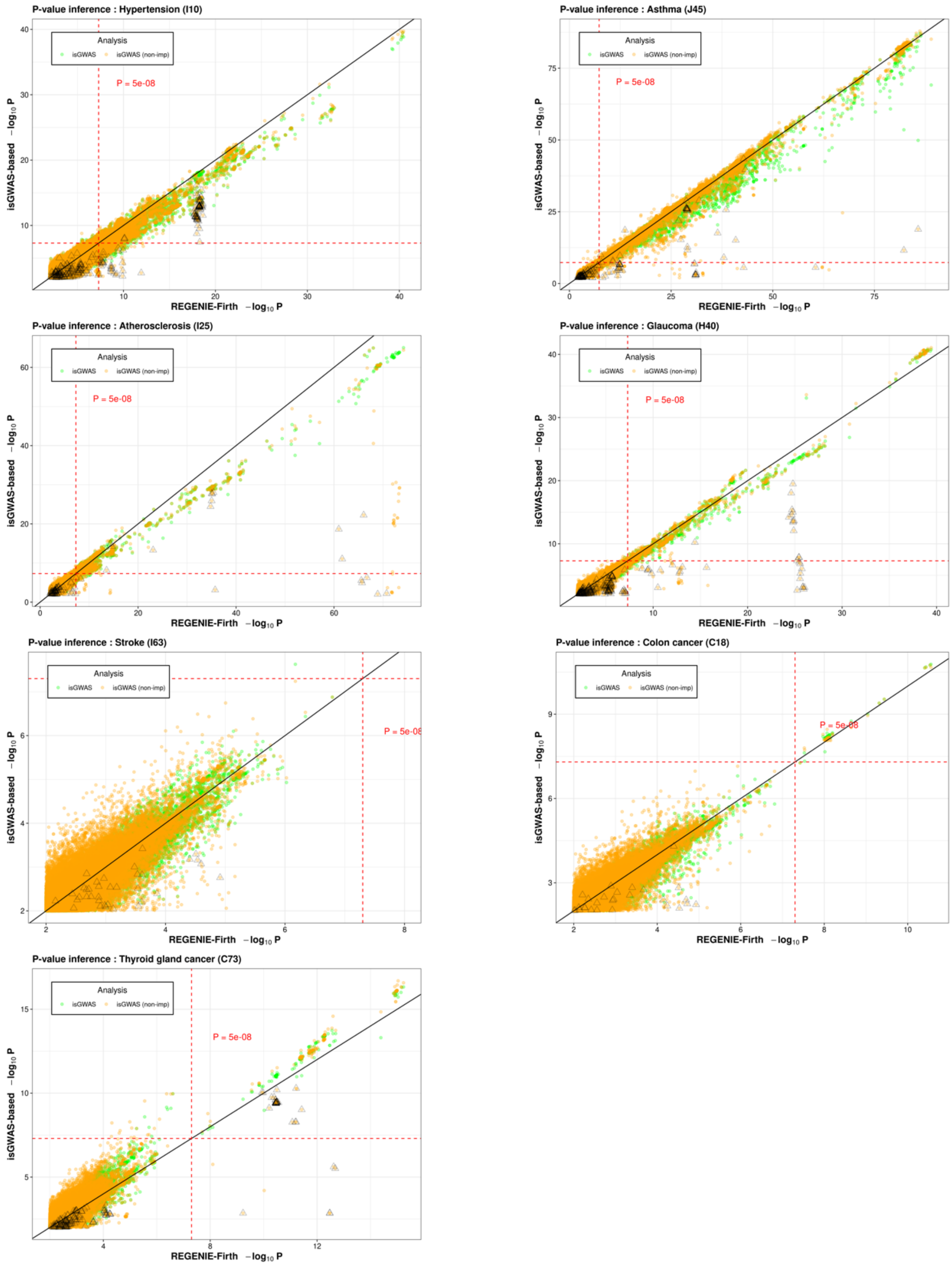

*Supplementary Figure 15. Performance of isGWAS with imputed and non-imputed MAFs. Representation of the values of $-\log_{10} P$ for isGWAS with imputed MAFs and REGENIE-Firth and the values for $-\log_{10} P$ for isGWAS with non-imputed MAFs and REGENIE-Firth. The classic threshold P-value is added for baseline. The values where imputed values lead to opposite beta direction are marked by triangle shape.*

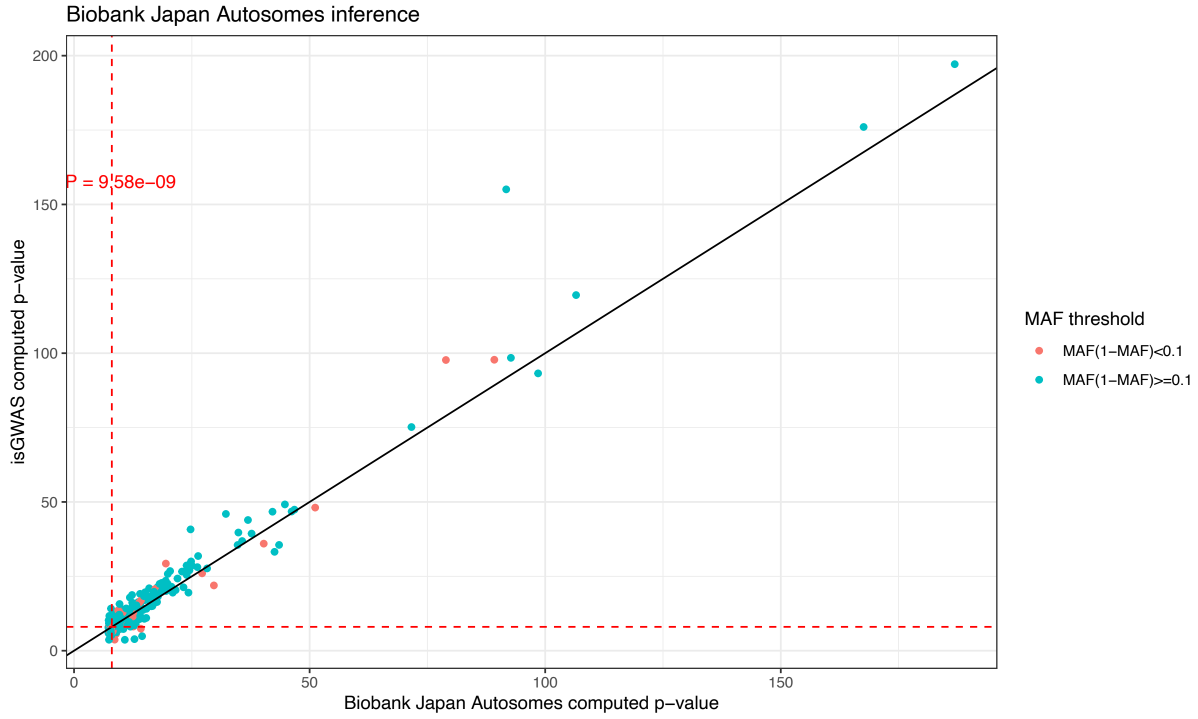

*Supplementary Figure 16. Inference results for Biobank Japan: population-level information for 309 significantly associated loci (P<5e-08) across 30 diseases has been used to infer p-values. The figure compares GWAS BBJ p-values and isGWAS computed p-values. The dashed red line represents the very stringent threshold P=9.58e-09 used by the authors in* [15]. *The results are coloured with respect to the MAF parameter to identify whether rare alleles influence the inferential nature of isGWAS.*

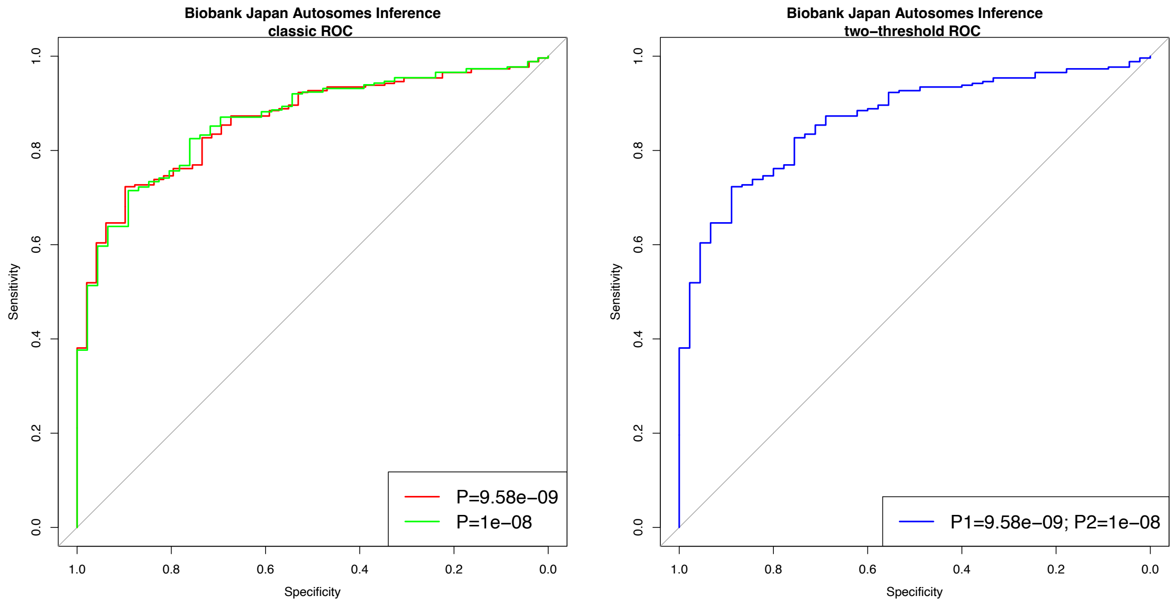

*Supplementary Figure 17. Inference results for Biobank Japan: classic ROC curve (left) and two-threshold ROC curve (right) on observed vs inferred values for autosomal variant results. The figure plots the ROC curves for different observed p-value*

*thresholds determining true/false label of autosomal variants' p-values. Lower threshold used for significance assessment provide better performance. The two-threshold ROC curve shows minimized false negatives and false positives even for tight difference between the thresholds.*

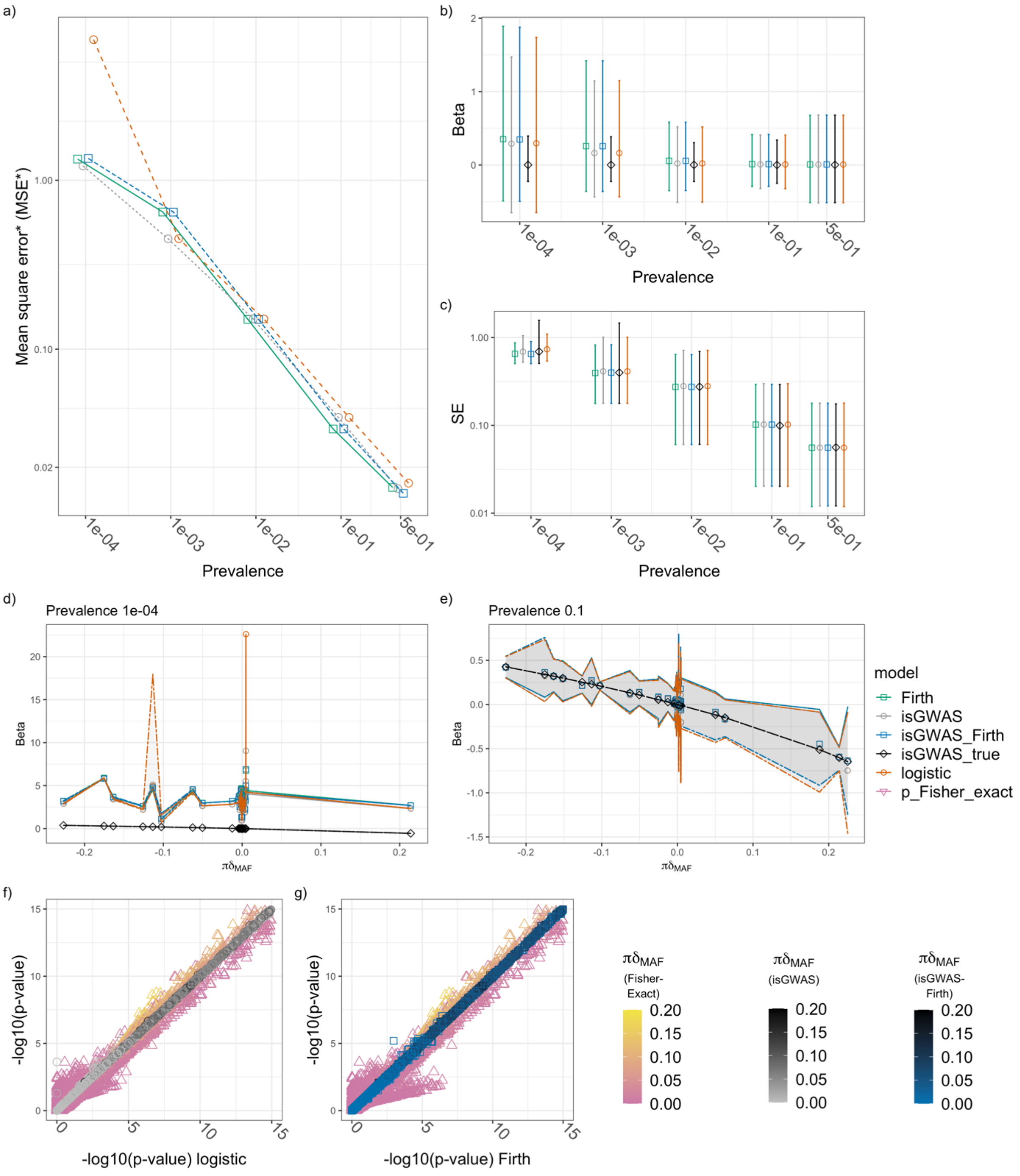

*Supplementary Figure 18.* ***Simulation II****: a) Mean square error (MSE*); b) distribution of estimated beta values and c) distribution of associated standard errors, for each model (logistic regression, firth regression, isGWAS and isGWAS-Firth) and specification of disease prevalence. Panels d)-e) illustrate the distribution of estimated beta values as a function of prevalence* $\pi$ *multiplied by the variable* $\delta_{MAF} = {(MAF^{*} - MAF)}\big/{\big(MAF(1 - MAF)\big)}$*, for* $\pi \in \{10^{-4}, 0.1\}$*. Panel f)-g) highlight the relationship between the logistic regression derived (f) and Firth regression derived (g)* -$log_{10}$(p-value) *along the horizontal axis and the corresponding isGWAS and Fisher Exact Test (FET) computed values on the vertical (note that we restricted the range of values presented to* $[0,15]$ *as FET regularly failed to converge for very small p-values). MSE* is computed assuming that the regression parameters derived from 'isGWAS_true' approximate the ground truth - isGWAS_true computes parameter estimates using the exact choice of population prevalence, MAF and MAF* used to generate a simulated dataset. In panels b)-e) a point denotes the median value and error-bars the first and ninth deciles of the range.*

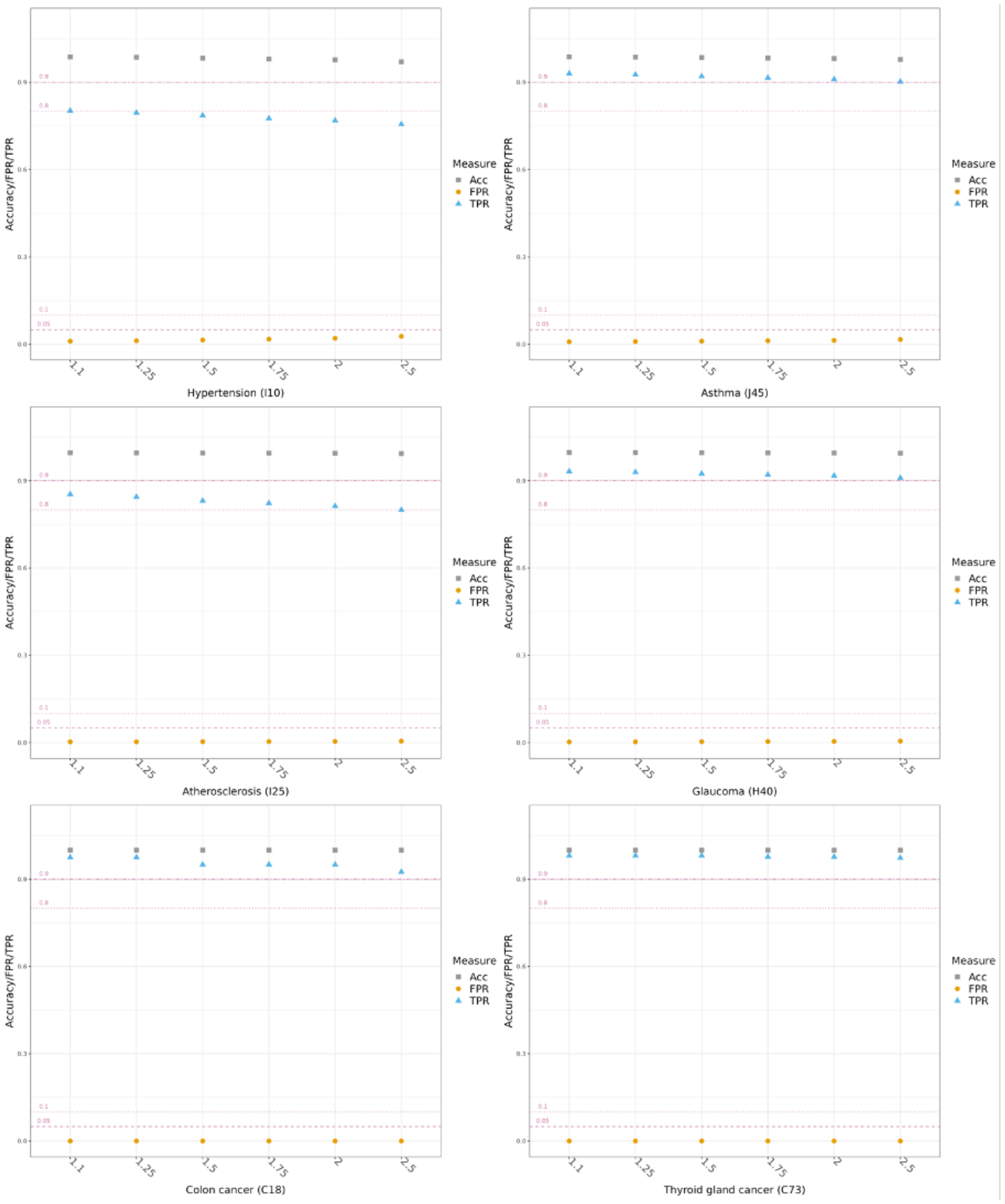

*Supplementary Figure 19. Performance of isGWAS in predicting future association results by increasing virtual sample size. Data are simulated using theoretic MAF ranges and prevalence scores from the UKB analysis for all variants with p<0.01. Using the isGWAS algorithm, the base sample size  from the UKB analysis has been virtually increased to  where , i.e., from a 10% to 150% increase in the number of 'samples'. Accuracy, true positive rate (TPR) and false positive rate (FPR) are presented. Stroke has been excluded from this assessment as no variants pass the significance threshold of* $p = 5e - 08$.

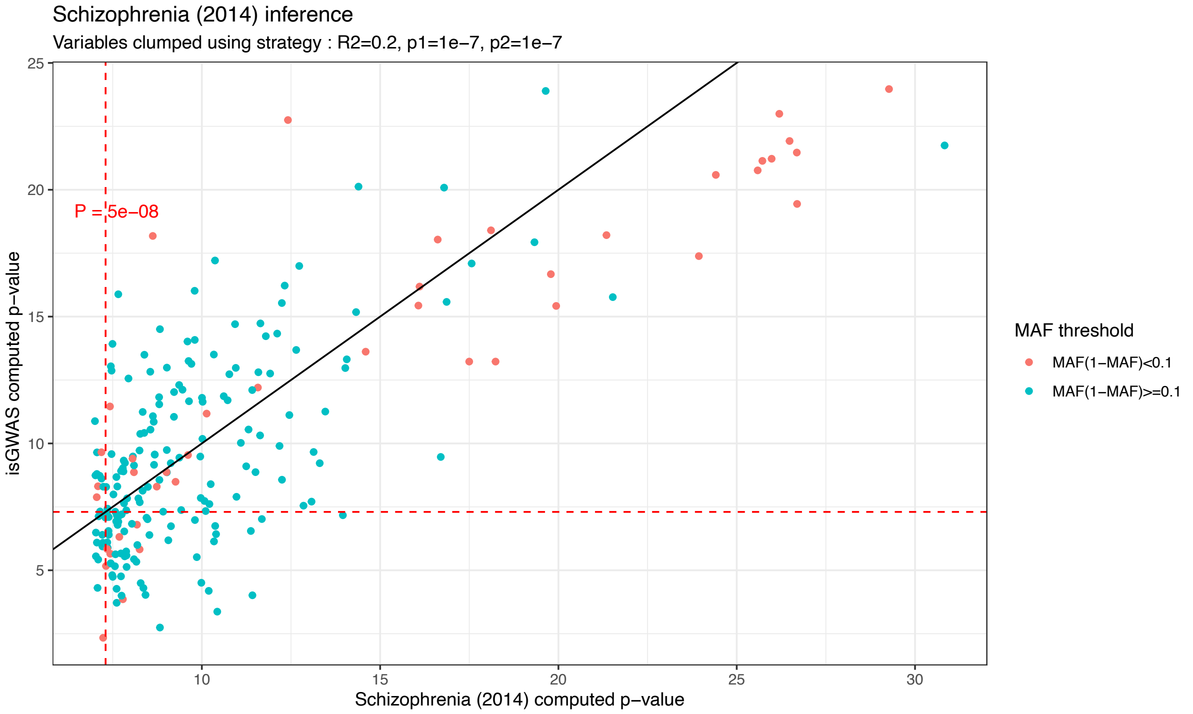

*Supplementary Figure 20. Inference results for Schizophrenia (2014): population-level information for 225 significantly associated loci (P<1e-07) has been used to infer p-values. The figure compares reported GWAS Schizophrenia (2014) p-values and isGWAS computed p-values. The dashed red line represents the threshold P=5e-08. The results are coloured with respect to the MAF parameter to identify whether rare alleles influence the inferential nature of isGWAS.*

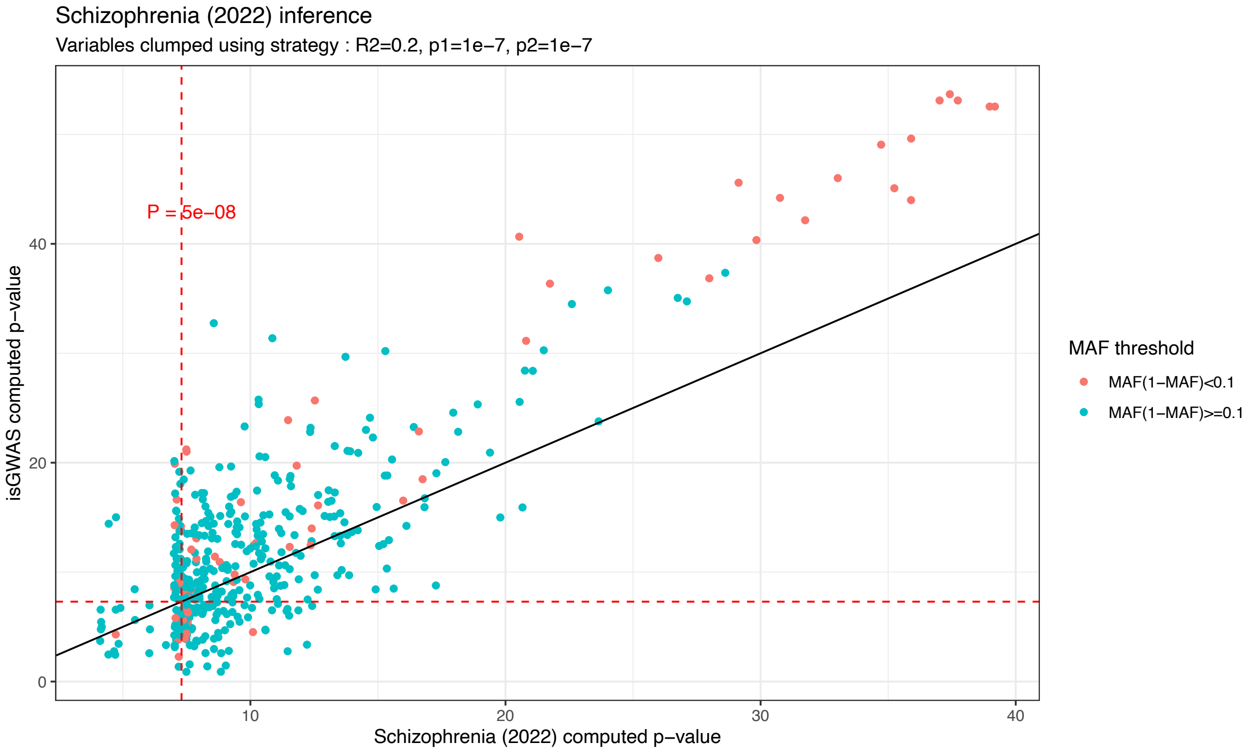

*Supplementary Figure 21. Inference results for Schizophrenia (2022): population-level information for 451 significantly associated loci (P<1e-07) has been used to infer p-values. The figure compares reported GWAS Schizophrenia (2022) p-values and isGWAS computed p-values. The dashed red line represents the threshold P=5e-08. The results are coloured with respect to the MAF parameter to identify whether rare alleles influence the inferential nature of isGWAS.*

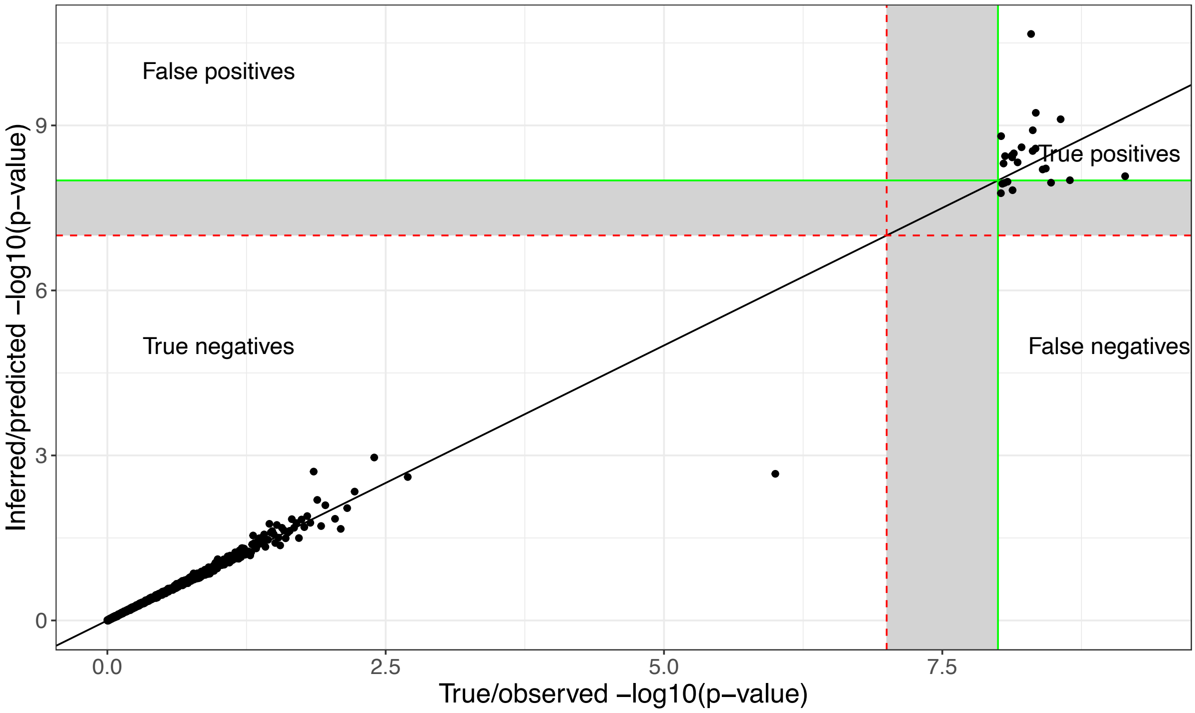

*Supplementary Figure 22. Classification performance of isGWAS: Visualization of confusion matrix on true vs predicted values. Given a significance assessment threshold p-value, we can identify the four regions of positives and negatives of the actual and predicted values. The shaded regions fall between different threshold regions: this shows that the choice of thresholds is an important one as it will have a large impact on the false positive and false negative rates.*
